# O-SNAP: A comprehensive pipeline for spatial profiling of chromatin architecture

**DOI:** 10.1101/2025.07.18.665612

**Authors:** Hannah H. Kim, Jose Angel Martinez-Sarmiento, Flavio R. Palma, Aayush Kant, Ellen Y. Zhang, Zixian Guo, Robert L. Mauck, Su Chin Heo, Vivek Shenoy, Marcelo G. Bonini, Melike Lakadamyali

**Affiliations:** Department of Physiology, Perelman School of Medicine, University of Pennsylvania, Philadelphia, PA 19104, USA; Biochemistry and Molecular Biophysics Graduate Group, Perelman School of Medicine, University of Pennsylvania, Philadelphia, PA 19104, USA; Center for Engineering Mechanobiology, University of Pennsylvania, Philadelphia, PA 19104, USA; Department of Metabolism and Physiology, H. Lee Moffit Cancer Center, Tampa, FL 33612, USA; Department of Materials Science and Engineering, School of Engineering and Applied Science, University of Pennsylvania, Philadelphia, PA, 19104, USA; Department of Bioengineering, School of Engineering and Applied Science, University of Pennsylvania, Philadelphia, PA, 19104, USA; McKay Orthopaedic Research Laboratory, Department of Orthopaedic Surgery, Perelman School of Medicine, University of Pennsylvania, Philadelphia, PA 19104, USA; Department of Mechanical Engineering and Applied Mechanics, School of Engineering and Applied Science, University of Pennsylvania, Philadelphia, PA 19104, USA

**Author notes:** Institute of Bioengineering, School of Life Sciences, École Polytechnique Fédérale de Lausanne, Lausanne, Switzerland (Jose Angel Martinez-Sarmiento). Center for Biomedical Engineering, Indian Institute of Technology Delhi, New Delhi - India (Aayush Kant).

## Abstract

We present O-SNAP (Objective Single-Molecule Nuclear Architecture Profiler), a comprehensive pipeline for the automated extraction, comparison, and classification of nuclear features from single-molecule localization microscopy (SMLM) data. O-SNAP quantifies 144 interpretable, biologically grounded spatial features describing chromatin organization or histone mark distributions at nanoscale resolution. The pipeline includes modules for pairwise comparison of features using volcano plots, feature set enrichment analysis, robust feature selection and classification of cell states, and pseudotime trajectory inference. We validate O-SNAP across diverse biological contexts, including fibroblast-to-stem cell reprogramming, tendon disease, histone variant sensitivity to oxidative stress, and chondrocyte de-differentiation, demonstrating its ability to detect subtle changes in nanoscale chromatin organization across diverse biological transitions.

## Introduction

The eukaryotic genome is hierarchically organized across multiple spatial and genomic scales, ranging from nanoscale chromatin structures to the global architecture of the nucleus^1,2^. At the most fundamental level, 146 base pairs of DNA are wrapped around histone octamers to form nucleosomes^3^ (10 nm), which compact further into higher-order structures including nucleosome clutches^4^ (20-100 nm), chromatin nanodomains^5,6^ (hundreds of nm), topologically associating domains (TADs)^7,8^, A/B compartments^9^, and ultimately chromosome territories^10^ (several micrometers). This multiscale organization is dynamically regulated by chemical modifications to DNA, such as methylation, and to histone tails, including site-specific methylation or acetylation, which modulate chromatin compaction and accessibility to transcriptional machinery^11–14^. Architectural proteins such as CTCF and cohesin further shape the genome by mediating long-range interactions and organizing it into domains such as TADs^15–18^. Additionally, associations with nuclear bodies, including the nuclear lamina^19^ and nuclear speckles^20,21^, contribute to functional compartmentalization, segregating the genome into transcriptionally repressive regions like lamina-associated domains (LADs) and active regions near nuclear speckles. This layered spatial compartmentalization is increasingly recognized as a key determinant of cell identity and function^22,23^. Understanding the relationship between the spatial organization of chromatin and cell state is therefore central to elucidating the molecular basis of developmental programming, cellular reprogramming, and disease-associated state transitions.

Recent advances have highlighted the importance of higher-order chromatin organization in shaping gene expression profiles across diverse cellular contexts. High throughput sequencing-based methods including chromatin immunoprecipitation sequencing (ChIP-seq), RNA sequencing (RNA-seq), assay for transposase-accessible chromatin using sequencing (ATAC-Seq) and chromosome conformation capture methods such as Hi-C, have been instrumental in revealing nuclear remodeling events in distinct cell states^24^. However, these approaches typically rely on population-aggregated information overlooking single-cell heterogeneity. Additionally, they often require chromatin extraction, resulting in the loss of native spatial context and obscuring the relationship between chromatin features and nuclear substructures. In contrast, microscopy-based approaches preserve nuclear architecture and enable direct visualization of chromatin and epigenetic modifications at single-cell resolution and in its native state. However, because chromatin structures span a broad size range, from nanometer-scale nucleosome clutches^4,25^ to micron-scale chromosome territories^26^, conventional light microscopy lacks the spatial resolution needed to resolve sub-diffraction chromatin features and visualize their remodeling dynamics.

Single-molecule localization microscopy (SMLM) encompasses a suite of super-resolution techniques that surpass the diffraction limit of light (∼200 nm), enabling visualization of sub-nuclear structures at nanoscale resolution^27,28^. These methods have been instrumental in uncovering previously inaccessible aspects of chromatin organization^24,29^. SMLM has revealed that chromatin is a disordered fiber composed of heterogenous clusters of nucleosomes or nucleosome clutches (also referred to as chromatin packing domains)^4,6,30^. Notably, clutch size has been shown to vary in a cell type–specific manner and correlates with the pluripotency status of both mouse embryonic stem cells and human induced pluripotent stem cells^4^. SMLM imaging further revealed progressive changes to chromatin organization and epigenetic modifications at different time points during somatic cell reprogramming^31^. Beyond stem cell contexts, SMLM has also demonstrated that chromatin architecture in mesenchymal stem cells (MSCs) is sensitive to mechanical cues, and chromatin remodeling is recapitulated in tissue-resident cells exposed to altered mechanical environments^32^. Additionally, SMLM imaging of tumor tissues has revealed fragmentation of heterochromatin in cancer cells^33^.

Collectively, these studies establish SMLM as a powerful tool to investigate chromatin and epigenetic remodeling during cell fate transitions and disease. However, a major limitation remains: the data generated by SMLM are highly complex and spatially rich, yet current analysis approaches often fail to capitalize on this wealth of information. Downstream analyses often rely on visual impression to guide quantification of a few simple parameters such as global chromatin compaction or chromatin domain size^4,32,34^. Such user-dependent decisions and reliance on simple measures of chromatin spatial organization are subjective and can lead to bias, and risk overlooking subtle, multidimensional patterns of chromatin reorganization.

Recent deep learning-based methods are enabling more sophisticated segmentation and quantification of super-resolution imaging data^35–42^ and these have been applied for accurate classification of cell states based on SMLM images of chromatin-associated components such as histones, DNA, or RNA polymerase II^43^. However, these methods rely on rasterizing the raw localization data into pixelated intensity maps, which can obscure fine-grained spatial information inherent to SMLM and introduce rendering bias as we previously demonstrated^44^. Additionally, the neural networks suffer from limited interpretability, making it difficult to pinpoint which spatial features drive classification decisions and impeding biological insight. Previously, our group developed ECLiPSE, a pipeline that extracts biologically meaningful morphological features from SMLM images and enables the quantitative classification of small subcellular structures such as protein aggregates and organelles^44^. While ECLiPSE provides interpretable outputs, its applicability is limited to relatively small and morphologically simple structures, and it is not optimized to analyze the spatially intricate architecture of the nucleus, which encompasses diverse and multiscale chromatin domains.

To address this gap, we introduce Objective, Single-molecule Nucleus Architecture Profiler (O-SNAP). O-SNAP begins by extracting a rich set of biologically grounded nuclear features from localization coordinates, preserving the resolution and spatial complexity of the original data. These features are then analyzed through an integrated suite of tools, including volcano plots, feature set enrichment analysis, pseudotime trajectory reconstruction, and classification models. Unlike the black-box approach of other methods, O-SNAP highlights which nuclear features strongly distinguish cell states, enabling users to trace classification outcomes back to specific spatial changes in chromatin architecture. This interpretability is critical for hypothesis generation and mechanistic understanding, empowering researchers to link nuclear organization with cellular identity and to uncover potential drivers of chromatin remodeling in development, reprogramming, and disease.

## Results

### O-SNAP enables automated, multiscale profiling of chromatin organization

O-SNAP begins by processing point cloud data (i.e. the raw x-y coordinates) of fluorophores detected via single-molecule localization microscopy (SMLM) for nuclear targets such as chromatin, histone modifications, or nuclear bodies, corresponding to high resolution spatial organization of these targets within the nucleus (**Figure 1**). The workflow integrates two distinct approaches to extract spatial features: (1) Voronoi tessellation is applied in a multiscale fashion to segment chromatin domains ranging from nanoscale to microscale^34,45,46^ (**Supplemental Figure 1A**), and (2) DBSCAN^47–50^ (Density-Based Spatial Clustering of Applications with Noise) is used to detect high-density, nanoscale chromatin packing domains (what we previously termed nucleosome clutches; **Supplementary Figure 1B**). Both approaches ultimately generate a set of domains that segment the localization data (see **Methods** for more details on clustering procedures and parameters). The domains are analyzed for key morphological properties including size, density (reflecting compaction), and radius of gyration (reflecting spatial dispersion). In addition to the average values, O-SNAP computes distributional statistics such as standard deviation (quantifying heterogeneity) and skewness (capturing asymmetry and deviations from normality).

**Figure 1.**
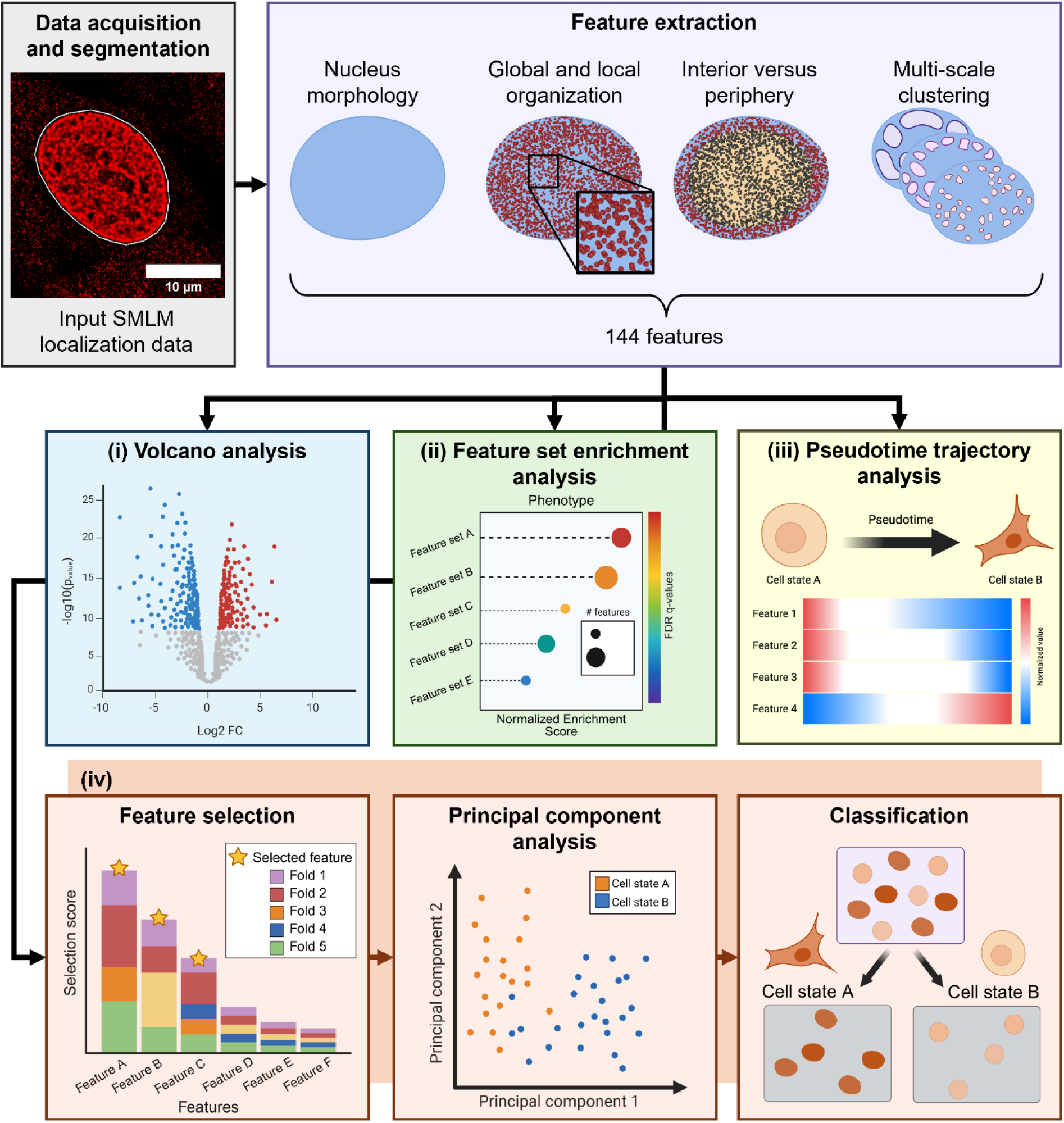
The O-SNAP workflow. The input data for O-SNAP consists of 2-D SMLM localization data of targets visualized in the nucleus. Following data acquisition and segmentation of the nucleus, O-SNAP calculates 144 features from the coordinate information of the SMLM data. Four types of analyses can then be independently conducted. (i) A volcano analysis identifies which features experience the greatest value change between two phenotypes. (ii) Feature set enrichment analysis determines how categories of features compare to each other from one phenotype to another. (iii) When relevant, pseudotime analysis builds a trajectory based on O-SNAP features to describe feature value changes as cells transition through multiple cell states over time. (iv) The final analysis in the pipeline identifies O-SNAP features with the highest potential to discriminate between different phenotypes, implements a principal component analysis (PCA) on these features, and generates a classification model to evaluate how sufficient these features are in distinguishing the conditions.

Given the importance of nuclear envelope to chromatin organization and function^19^, a compartmental analysis compares chromatin properties between the nucleus periphery, defined as the region near the nucleus boundary within the outermost 15% of the nucleus radius, and the nucleus interior which is the remaining innermost region (**Supplementary Figure 1B**). Additionally, O-SNAP incorporates a previously established modeling approach^51^ that quantifies the thickness and length of lamina-associated heterochromatin, which considers a more stringent subset of the dense chromatin compartment at the nuclear periphery, where 5% of the nucleus radius from the boundary is used as a distance cutoff to determine putative lamin associated domains (LADs). (**Supplementary Figure 1B**). To evaluate radial organization, O-SNAP divides the nucleus into ten concentric zones from the interior to the periphery and quantifies chromatin domain abundance and feature gradients across these zones (**Supplementary Figure 1C**). The zones are defined by evenly partitioning the major and minor axes of the nucleus into ten segments, which are then used to generate elliptical boundaries for each radial bin. (**Supplementary Figure 1C**). Due to the irregular shape of many nuclei, some peripheral localizations may fall outside the outermost ellipse. To ensure these edge-localizations are not excluded, the outermost zone is defined as including all localizations or domains that lie beyond the ninth ellipse (formed from 90% of the length of the major and minor axis, **Supplementary Figure 1C**).

In total, the O-SNAP pipeline extracts 144 biologically inspired spatial features encompassing nuclear morphology, global and radial chromatin organization, and multiscale domain architecture (**Figure 1** and **Supplementary Tables 1-3**). For more details on O-SNAP features, we direct readers to **O-SNAP feature generation** under the **Methods** section.

#### Downstream analysis modalities in O-SNAP

Once biologically inspired spatial features are extracted, O-SNAP offers multiple downstream analytical modalities for comparing chromatin architecture across distinct cell states (**Figure 1**). Drawing inspiration from transcriptomic workflows such as RNA sequencing (RNA-Seq), these include: (i) volcano plots to identify the most differentially changing spatial features between two groups; (ii) feature set enrichment analysis (FSEA) to reveal coordinated shifts within functionally related feature families; (iii) pseudotime trajectory analysis to temporally order cells based on progressive changes in chromatin features; and (iv) feature selection and classification using machine learning models trained to distinguish between cell states based on spatial patterns of chromatin organization (**Figure 1**). These complimentary analyses enable both hypothesis-driven and data-driven interrogation of nuclear architecture across cellular transitions.

##### Volcano Plot Analysis

The volcano analysis in O-SNAP is performed pairwise between specified phenotypes. For each feature, the fold change is calculated between the two phenotypes and statistical significance is assessed using a two-tailed t-test assuming unequal variance. The p-values are adjusted for multiple comparisons using the Benjamini-Hochberg method. Here, features with an absolute fold change greater than 2 and an adjusted p-value satisfying α < 0.05 were considered significant, although these thresholds are user-configurable and can be modified based on specific experimental expectations.

##### Feature Set Enrichment Analysis (FSEA)

The feature set enrichment analysis (FSEA) in O-SNAP is inspired by Gene Set Enrichment Analysis (GSEA)^52^, and builds upon the MrGSEA implementation^53^ (**Supplementary Figure 2**). FSEA determines which sets of features are more strongly associated with a given phenotype class. Feature sets are formed by manually categorizing features based on a general description of chromatin change they correspond to (**Supplementary Figure 2A** and **Supplementary Table 3**). For example, the feature “M.25” (nuclear radius) belongs to the set “larger nucleus”. Each set is evaluated to determine whether the features that comprise it are enriched in either of the two classes.

For a given feature set, *S*, the calculation of its enrichment score *ES*(*S*) begins with the normalization of the full dataset into z-scores (**Supplementary Figure 2A**). The features are then ranked according to their correlation with phenotype, which can be calculated using one of 8 possible ranking metrics (**Supplementary Figure 2A** and **Supplementary Table 4**). Here, we used the default metric, “Signal-to-noise” (“S2N”), which ranks features based on the difference between the means of each class and divides this by the sum of their standard deviations.

Unlike in sequencing data, where higher expression always indicates positive enrichment, an increase in an O-SNAP feature value may reflect a change in the opposite direction of its associated feature set. For example, the feature “M.1” (nucleus aspect ratio, the ratio between the major axis to the minor axis) has a value near 1 when the nucleus shape approximates a circle and increases in value the more elongated the nucleus becomes. M.1 is grouped under the “rounder nucleus” set, but an increase in aspect ratio corresponds to a more elongated, less round nucleus. To avoid redundancy in feature sets (e.g. including both “rounder nucleus” and “less round nucleus” as separate sets), a correction is applied to the ranking metric of the features whose increase opposes the direction of their respective feature set. For instance, with the “S2N” metric, the additive inverse is used in place of the original value when ranking the features whose increase opposes the described feature set. The correction for each ranking metric is detailed in **Supplementary Table 4**.

Once the ranking metric for all features is calculated, the features are sorted, where features near the top of the ranking correspond to greater enrichment in one of the classes. A cumulative score is obtained for a given feature set by traversing down the ranking. The score increases or decreases with each feature depending on whether the feature is a member of the set of interest (*P*_hit_ and *P*_miss_, respectively), where the magnitude depends on the correlation between the given feature and phenotype (**Supplementary Figure 2B**). The maximum value of the score over the course of the ranking is used as the reported *ES*(*S*) for a given feature set and resembles the weighted Kolmogorov-Smirnov statistic (**Supplementary Figure 2C**).

To test for statistical significance, a permutation method is used to obtain a null distribution of the enrichment score (**Supplementary Figure 2D**). The feature data undergoes 1,000 permutations of the phenotype labels, which maintains feature correlation structure within the data, and for each permutation, *π*, a new ranking is calculated with a new result for *ES*(*S*, *π*). This distribution is used to rescale the scores to normalized enrichment scores, *NES*(*S*), accounting for the size of the feature set, enabling the comparison of scores between different sets (**Supplementary Figure 2D**). The *NES* are analyzed against the normalized, permuted enrichment scores to obtain p- and q-values for statistical significance under multiple hypotheses (**Supplementary Figure 2D, E**, see Subramanian et al.^52^ for additional details).

##### Pseudotime Trajectory Analysis

O-SNAP’s pseudotime trajectory inference (TI) module (**Supplementary Figure 3**) similarly adapts established methods from single-cell sequencing trajectory analysis to operate on O-SNAP feature data. Here, we utilized the *dynverse* suite of R packages, which supports the exploration of multiple TI algorithms^54^. Three candidate TI methods were identified using the *dynguidelines* tool (SCORPIUS^55^, Embeddr^56^, and Slingshot^57^; **Supplementary Table 5**). The Slingshot method^57^ was selected for the results presented here, as it best separated the experimental timepoints of the heterokaryon reprogramming dataset used here along pseudotime. Based on findings from Saelens et al^54^, Slingshot’s superior performance on our reprogramming dataset may be attributed to its overall robustness across a range of criteria; including accuracy, stability, scalability, and usability; as well as its enhanced ability to model simple trajectory topologies, such as the linear progression assumed for the heterokaryon reprogramming system. Note that other biological systems may be better represented using alternative trajectories or inference methods, which are available through the *dynverse* package. Instead of gene expression data, O-SNAP feature values, which are exported from the MATLAB pipeline as a table in CSV format and normalized to z-scores, were used as input. Because *dynverse* requires raw sequencing count data for its input, we assigned a constant dummy value (0.5) to satisfy this requirement. This value does not influence the trajectory outcome and serves only to meet the software’s input format expectations. ^54^.^55–57^

A limited amount of prior information was provided to guide the inference. Given our expectation of a continuous, non-branching transition between cell states, the trajectory was constrained to a linear topology with defined start and end phenotypes. In scenarios where the transition might be more complex, the *dynverse* package offers alternative topologies and methods that can accommodate branching systems, provided the dataset is sufficiently large for each cell state. A basic understanding of R is required to modify and run scripts for *dynverse*, which provides its own tutorial; an example script is provided as *Pseudotime_Example_Heterokrayon.R*.

Overall, applying a pseudotime framework to SMLM data enables us to infer how chromatin features spatially reorganize along a continuum of cell states, thereby facilitating the analysis of long-term dynamic changes across the inferred trajectory.

##### Feature Selection, Principal Component Analysis, and Classification

The final analysis module in O-SNAP is the classification pipeline (**Supplementary Figure 4**), designed to ensure robust performance while minimizing the risk of data leakage. The dataset is partitioned into multiple pairs of training and test data. Five splits are created using five-fold cross-validation strategy, where each fold reserves a different one-fifth of the data as the test set while the remainder is used for training (**Supplementary Figure 4A**). The pipeline performs a feature selection to identify a subset of the original 144 features that best discriminates between cell states. The Minimum Redundancy Maximum Relevance (MRMR) algorithm ranks features to maximize their correlation with cell state while minimizing redundancy among them (**Supplementary Figure 4B**). The rankings from the five folds are summed together, and the top features with the largest cumulative score are selected. The cutoff for the number of selected features is determined automatically based on the “knee” point of the aggregate ranking curve or the point with the largest difference between the consecutive scores, whichever yields the smaller feature set (**Supplementary Figure 4C**). The filtered feature subset is then subject to principal component analysis (PCA) analysis, which serves both as dimensionality reduction step and as a visualization tool of the underlying structure in the data independent of class labels to assess the data (**Supplementary Figure 4D**). The PCA-transformed data is used to train a suite of classification model architectures and the model with the highest average accuracy across all folds is reported as the final output (**Supplementary Figure 4D**).

### The O-SNAP pipeline recapitulates the impact of histone deacetylase inhibition and tendinosis on chromatin spatial organization

To validate O-SNAP, we applied the pipeline to two well-characterized systems known to exhibit distinct and opposing changes in chromatin spatial organization: (1) human fibroblasts (hFbs) treated with the histone deacetylase inhibitor Trichostatin A (TSA; **Figure 2**), and (2) primary tenocytes isolated from healthy individuals versus individuals with tendinosis (**Figure 3**). TSA treatment induces global hyperacetylation of histone tails, weakening nucleosome-nucleosome interactions and leading to chromatin decompaction due to increased electrostatic repulsion between nucleosomes^34^. In prior work using SMLM imaging, we showed that TSA treatment results in the fragmentation of large, dense chromatin packing domains, which we referred to as nucleosome clutches, into smaller, less compact, and more spatially dispersed domains^4,34^. In contrast, tenocytes from tendinosis patients exhibit increased heterochromatin formation, with chromatin becoming sequestered at the nuclear lamina, a repressive nuclear compartment enriched for constitutive heterochromatin^32^. These two systems thus provide contrasting benchmarks for evaluating O-SNAP’s ability to detect spatial reorganization of chromatin.

**Figure 2.**
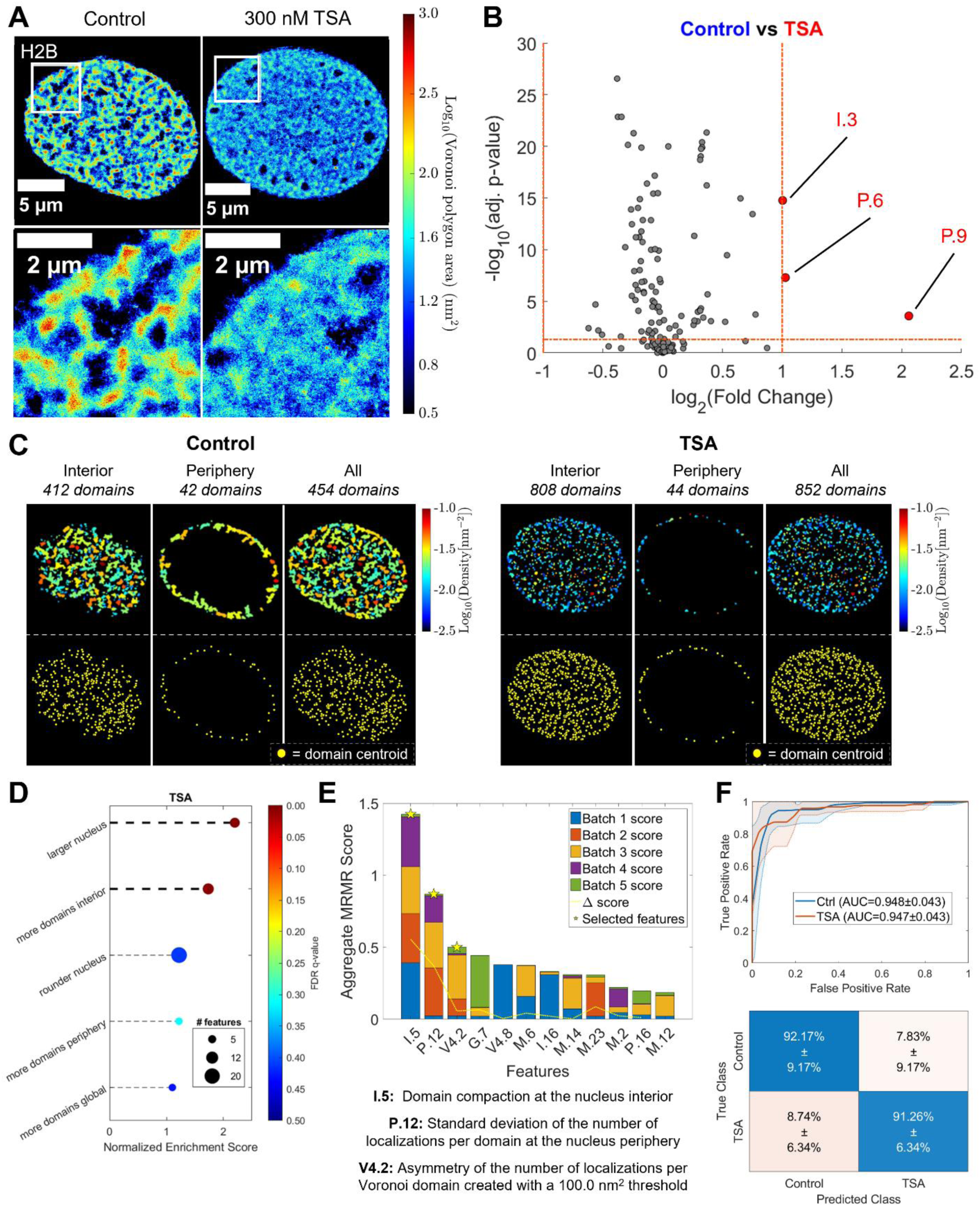
O-SNAP successfully separates TSA treated cells from control cells and identifies chromatin features of interest. (A) Representative Voronoi density map renderings of H2B STORM data for control and TSA-treated fibroblasts. (N=117 nuclei for the control and N=106 for the TSA-treated group). The color code indicates local chromatin compaction from low density (blue) to high density (red). (B) A Volcano plot visualizes fold changes in O-SNAP-generated features of control and TSA-treated fibroblasts. Three O-SNAP features increase in value in TSA-treated cells. The statistical significance was calculated using a two-sided t-test followed by Benjamini-Hochberg adjustment for multiple comparisons. (C) Visualizations of the chromatin packing domains, (i.e. the compact chromatin compartment segmented using DBSCAN), from the representative nuclei from panel A. Top: Packing domains are color-coded based on their respective localization density for control (left) or TSA-treated cells (right). Bottom: Each point represents the centroid position of a single chromatin packing domain. (D) FSEA indicates which overall nucleus characteristic TSA-treated samples trend towards, here being features related to a larger nucleus and those that indicate more domains at the nucleus interior. The color code corresponds to the statistical significance of the normalized enrichment score, measured by the False Discovery Rate (FDR) q-value, and the size of the icon corresponds to the number of features contained in each feature set. (E) Aggregated MRMR score across the five folds. Three features (I.5, P.12 and V4.2) were selected. (F) Classification results across five training/test folds for a gaussian SVM model trained to discriminate control from TSA-treated fibroblasts using the features selected in (E) and labels with the ground-truth cell state, which has an overall accuracy of 91.76 ± 5.22%. Top: ROC curve where the solid line indicates the average value across the folds, and the shaded region indicates the ±1 standard deviation interval. Bottom: A confusion matrix shows the model’s average accuracy for the control and TSA-treated conditions across the five folds.

**Figure 3.**
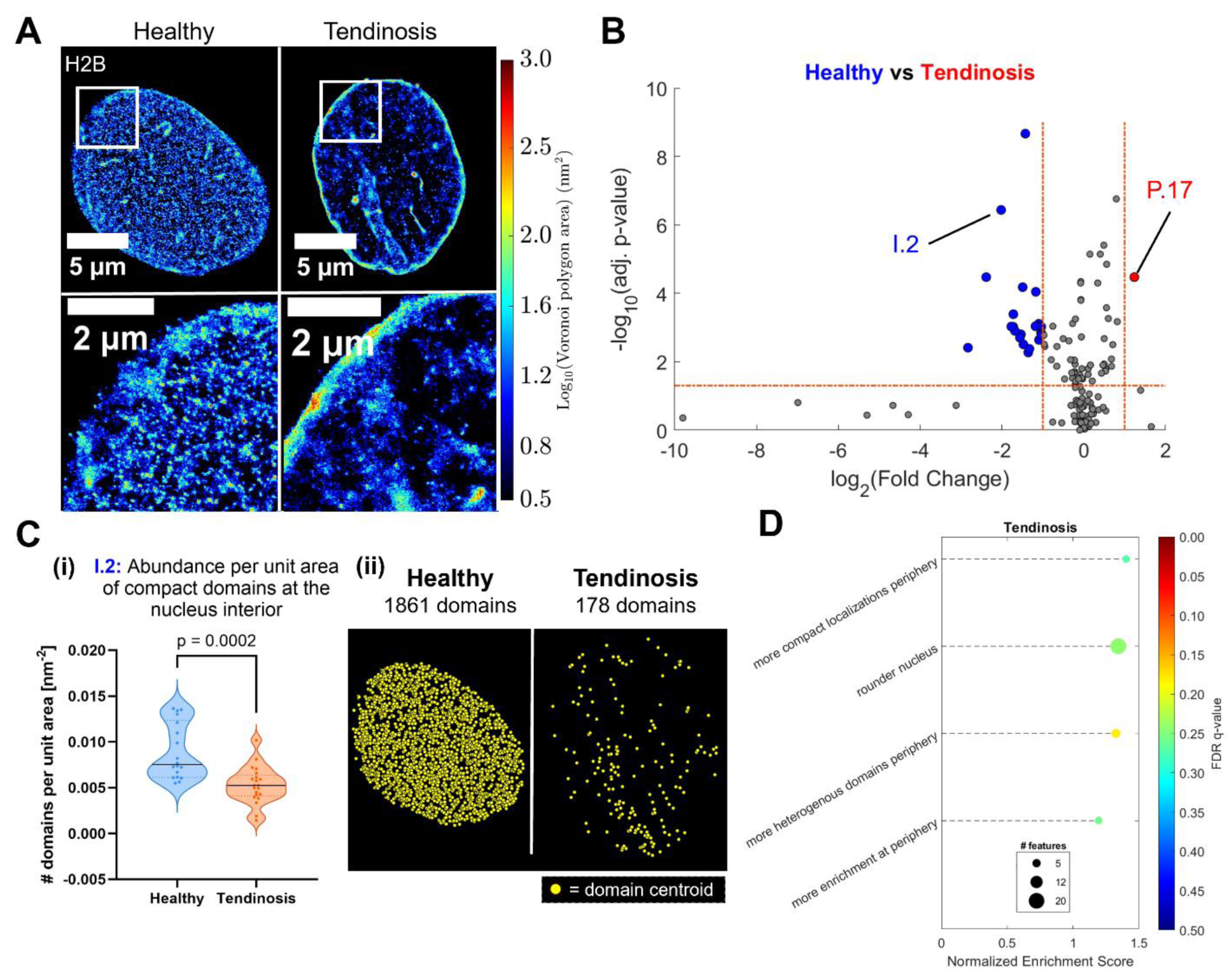
O-SNAP successfully separates tenocytes derived from tendinosis versus healthy patients and identifies chromatin features of interest. (A) Representative Voronoi density map renderings of H2B STORM data of tenocyte cells derived from healthy donors or those with tendinosis (N=19 for healthy cells and N=21 for tendinosis cells). The color code indicates local chromatin compaction from low density (blue) to high density (red). (B) A Volcano plot visualizes fold changes in O-SNAP-generated features of nuclei from healthy or tendinosis donors. One feature increased in value in tendinosis cells while 22 features increased in value in healthy cells. The statistical significance was calculated using a two-sided t-test followed by Benjamini-Hochberg adjustment for multiple comparisons. (C) (i) Violin plots showing the distribution of Feature I.2, the number of packing domains at the nucleus interior, for the healthy and tendinosis nuclei imaged, showing a decrease in tendinosis nuclei (ii) Representative nuclei, where each point represents the centroid position of a single packing domain for healthy (left) and tendinosis (right) nuclei. (D) FSEA indicates which overall nucleus characteristics tendinosis tenocytes trend towards. Features related to more heterogeneous domains at the periphery and rounder nuclei trend towards enrichment but are not statistically significant for α < 0.05. The color code corresponds to the statistical significance of the normalized enrichment score, measured by the False Discovery Rate (FDR) q-value, and the size of the icon corresponds to the number of features contained in each feature set.

We first applied O-SNAP to compare the global chromatin organization in wild-type and TSA-treated hFbs (**Figure 2A**) using volcano analysis to identify the most significantly altered spatial features (**Figure 2B**). Applying fold-change and significance cutoffs revealed three prominent feature changes, particularly within the nuclear interior (I) and periphery (P) (**Figure 2B, C** and **Supplementary Figure 5A-C**). At the interior, DBSCAN clustering detected a significant increase in the number of chromatin packing domains (nucleosome clutches) following TSA treatment (Feature I.3; **Figure 2B, C**, **Supplementary Figure 5A**, and **Supplementary Table 6**), consistent with prior findings that large, compact domains fragment into smaller, more numerous units upon chromatin decompaction^34^.

At the nuclear periphery, the two most significantly changing features related to the asymmetry in the distributions of chromatin domain compaction (Feature P.6) and radius of gyration (Feature P.9; **Figure 2B** and **Supplementary Figure 5A**). Closer examination of the DBSCAN-segmented chromatin packing domains revealed a general decrease in both density (**Figure 2C**) and radius of gyration (**Supplementary Figure 5B**) at the periphery, consistent with decompaction. However, a subset of peripheral domains remained highly compact or spatially extended (**Figure 2C** and **Supplementary Figure 5B**), leading to a heavy right tail in these distributions and contributing to increased asymmetry (**Supplementary Figure 5C**). These findings suggest that TSA treatment affects peripheral heterochromatin heterogeneously, inducing widespread decompaction while leaving a minority of domains resistant. These results validate O-SNAP’s ability to recapitulate known chromatin remodeling events (more numerous, less compact chromatin packing domains) and highlight its potential to uncover novel, interpretable spatial features that point to new hypotheses (heterogenous impact of TSA on peripheral heterochromatin compaction).

FSEA further supported these findings, revealing an increased number of chromatin domains globally, at the periphery, and in the interior (**Figure 2D**). Nuclear morphology features such as increased nuclear size and roundness also emerged, consistent with chromatin decompaction driving nuclear expansion^58,59^. Hence FSEA analysis provides both overlapping and complementary information to volcano plot analysis, highlighting the added value of distinct analysis modalities.

Finally, we applied MRMR feature selection and classification (**Figure 2E, F**). O-SNAP achieved 91.76 ± 5.22% accuracy in distinguishing TSA-treated from wild-type cells (**Figure 2F**), which decreased to 57.99 ± 9.04% upon random shuffling of the data labels (**Supplementary Figure 5D**). Details on how the confusion matrix accuracies and the overall accuracy are computed can be found in the **Classification** section under the **Methods**. The most predictive features involved changes in the compaction of interior chromatin and heterogeneity of histone amounts within the peripheral domains (**Figure 2E**), aligning with the volcano analysis results on asymmetry (**Figure 2B**). Plotting the three selected features revealed clear separation between wild-type and treated cells (**Supplementary Figure 5E**).

We next applied the O-SNAP pipeline to primary tenocytes isolated from healthy individuals and from patients with tendinosis (**Figure 3A**)^32^. Volcano analysis revealed 23 significantly altered spatial features (**Figure 3B** and **Supplementary Table 7**). Notably, among these changes was the increased ratio of chromatin at the nuclear periphery relative to the interior (Feature P.17; **Figure 3B** and **Supplementary Figure 6A**), recapitulating our prior observation that chromatin becomes sequestered at the nuclear lamina in tendinosis^32^. This peripheral accumulation was accompanied by a reduction in the number of chromatin domains at the nuclear interior (Feature I.2; **Figure 3B, C**), reflecting a shift in chromatin organization away from the interior toward the periphery. FSEA corroborated these findings, revealing increased chromatin enrichment and greater domain heterogeneity at the nuclear periphery, along with a trend toward rounder nuclear morphology (**Figure 3D**). These results provide complementary evidence of large-scale nuclear remodeling in disease. Finally, classification analysis distinguished healthy and tendinosis-derived tenocytes with an accuracy of 95.00 ± 6.85% (**Supplementary Figure 6B, C**). The most discriminative features included the number of chromatin domains at the periphery and the spatial distribution (spacing) of domains within the nuclear interior (**Supplementary Figure 6B**), again reflecting the redistribution of chromatin from interior to peripheral compartments.

Together, these analyses across two previously validated systems demonstrate that O-SNAP can reliably detect biologically meaningful changes in chromatin architecture and provide complementary, interpretable insights across multiple analytical modalities.

### O-SNAP reveals a detailed overview of how global chromatin and specific epigenetic marks remodel in early reprogramming

We next evaluated O-SNAP on a more complex and heterogeneous system: heterokaryon-mediated reprogramming of somatic human fibroblasts (hFbs) via fusion with mouse embryonic stem cells (mESCs). In this model, the fusion of somatic and pluripotent cells induce reprogramming that is initiated by diffusible mESC factors acting on the hFb nucleus^60–63^, and we previously showed that chromatin undergoes progressive decompaction over time following fusion^31^. Unlike TSA treatment or tendinosis, which induce relatively uniform and drastic chromatin changes, heterokaryon reprogramming generates more subtle changes particularly at early time points after cell fusion and a large amount of cell-to-cell heterogeneity^31^. These aspects of the heterokaryon dataset pose a more challenging test case for O-SNAP.

We first analyzed three cell types using SMLM images of histone H2B, corresponding to global chromatin organization (**Figure 4A**): control hFbs prior to fusion, control mESCs prior to fusion, and partially reprogrammed hFbs imaged 48 hours post-fusion^31^. Despite the inherent heterogeneity, O-SNAP achieved an average of 80.94 ± 5.28% classification accuracy in distinguishing these states, dropping to 41.08 ± 6.71% upon random label permutation (**Figure 4B, C**), demonstrating the pipeline’s robustness in complex biological settings.

**Figure 4.**
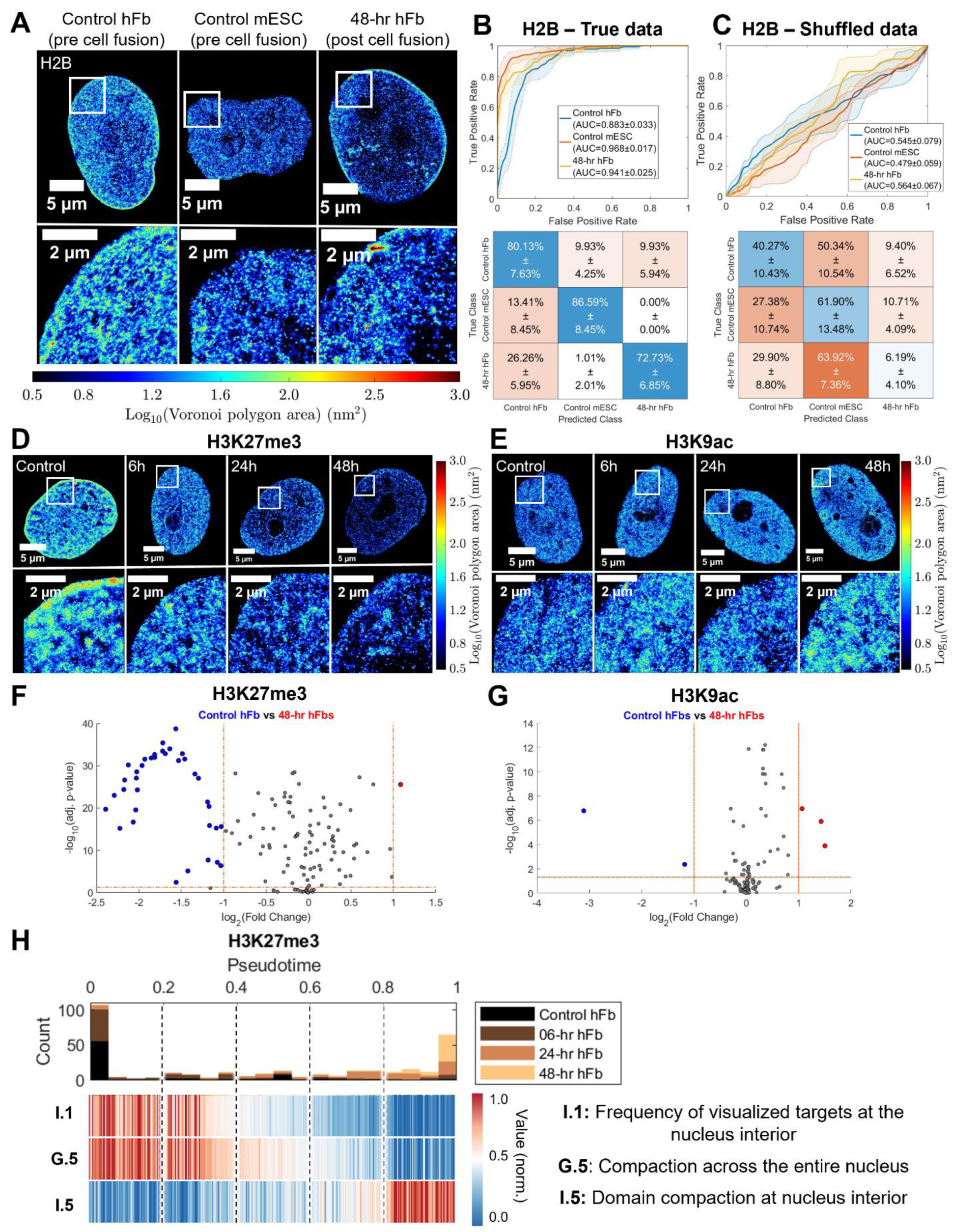
O-SNAP applied to the heterokaryon reprogramming dataset elucidates changes between multiple cells states and epigenetic marks. (A) Representative Voronoi density map renderings of H2B STORM images of control human fibroblasts (hFbs), mouse ESCs, and hFb nuclei 48-hr post cell fusion (N=151 nuclei for control hFbs, N=164 for control mESCs, and N=99 for 48-hr hFb nuclei post cell fusion). The color code indicates local chromatin compaction from low density in blue to high density in red. (B) A logistic regression classification model trained on the heterokaryon H2B data across five training/test folds using ground truth labels. The results of the classifier are shown with a ROC curve (top) and confusion matrix (bottom) across the five folds, where the overall classification accuracy is 80.94 ± 5.23%. (C) An ensemble boosted trees classification model across five training/test folds, shuffling the cell state labels of the heterokaryon H2B system. The average results of the ROC curve (top) and confusion matrix (bottom), where the overall classification accuracy across the five folds is 41.08 ± 6.71%. (D) Representative Voronoi density map renderings of H3K27me3 STORM data of somatic heterokaryon nucleus at different timepoints post-fusion (N=92 nuclei for control hFbs and N=61 for 48-hr hFb nuclei post cell fusion). (E) Representative Voronoi density map renderings of H3K9ac STORM data of somatic heterokaryon nucleus at different timepoints post-fusion (N=93 nuclei for control hFbs and N=62 for 48-hr hFb nuclei post cell fusion). (D,E) The color code indicates local chromatin compaction from low density in blue to high density in red. (F) Volcano plot visualizing fold changes in H3K27me3 O-SNAP-generated features of control fibroblast nuclei or fibroblast nuclei within hFb nuclei 48-hr post cell fusion. 36 features increased in value in control hFbs and one increased for 48-hr hFb nuclei post cell fusion. (G) Volcano plot visualizing fold changes in H3K9ac O-SNAP-generated features of control fibroblast nuclei or fibroblast nuclei within hFb nuclei 48-hr post cell fusion. Three features increase in value in the 48-hr hFb nuclei post cell fusion and two features increase in value in control hFbs. (F, G) The statistical significance was calculated using a two-sided t-test followed by Benjamini-Hochberg adjustment for multiple comparisons. (H) Top: Histogram of assigned pseudotimes calculated using the Slingshot trajectory inference method on O-SNAP features generated from the H3K27me3 heterokaryon SMLM data. Bottom: A heatmap of H3K27me3 O-SNAP features changing with pseudotime. The features displayed are a subset of those determined to be most predictive to pseudotime.

To assess how chromatin remodeling correlated with histone modifications, we additionally analyzed two marks, H3K27me3 representing a repressive mark (**Figure 4D**) and H3K9ac representing an active mark (**Figure 4E**), which we previously showed to be differentially regulated during reprogramming^31^. Volcano analysis revealed that 36 spatial features significantly decreased and one increased for H3K27me3 in hFb nuclei after 48 hours of reprogramming (**Figure 4F** and **Supplementary Table 8**), while only 5 features changed for H3K9ac (**Figure 4G** and **Supplementary Table 9**), consistent with our previous findings that the repressive heterochromatin is selectively depleted during early heterokaryon reprogramming while the levels of active euchromatin are maintained^31^. Accordingly, classification accuracy decreased from 97.40 ± 1.46% with H3K27me3 data to 83.87 ± 3.95% with H3K9ac data when distinguishing between two cell states: the somatic hFb nucleus of the heterokaryon at 48-hr post-fusion and the control hFbs prior to fusion (**Supplementary Figure 7A, B**).

To further probe dynamic chromatin remodeling, we applied pseudotime trajectory analysis using the R package, *dynverse*^54^, to the H3K27me3 images of hFb nuclei at 0-, 6-, 24-, and 48-hr post-fusion (**Figure 4H**). The resulting trajectory captured a continuous transition, with control hFbs prior to fusion (0 hr) clustering at the beginning and hFbs at 48-hr post fusion clustering at the end of the timeline (**Figure 4H**). In contrast, applying the same pseudotime analysis to global chromatin organization visualized via H2B yielded a less distinct trajectory, with cells from different time points more evenly distributed and inter-mixed along the pseudotime axis (**Supplementary Figure 7C**). These results suggest that O-SNAP-extracted spatial features can capture temporal dynamics in chromatin architecture, and that repressive H3K27me3-marked chromatin provides greater sensitivity for resolving these transitions than global chromatin structure alone. Examining the trajectory further revealed that H3K27me3 abundance declined early, followed by global decompaction of H3K27me3 marked chromatin (**Figure 4H** and **Supplementary Figure 7D**). Intriguingly, at the end of the trajectory, chromatin packing domains (or clutches) exhibited increased compaction (Feature I.5 in **Figure 4H** and **Supplementary Figure 7D**), suggesting the possible emergence of persistent, highly compact heterochromatin nanodomains that may be resistant to remodeling, a hypothesis that warrants further investigation in future studies. While these trends were also present in the H2B data, evidenced both along the pseudotime trajectory as well as the distributions according to the ground-truth labels, the magnitude of change for all features was much more modest compared to the trends in the H3K27me3 data (**Supplementary Figure 7C, D**).

These findings demonstrate that O-SNAP not only captures broad, time-evolving changes in chromatin state but also reveals residual chromatin features, which may be potentially linked to incomplete reprogramming.

### Impact of oxidative stress on chromatin spatial organization in human mammary epithelial cells

Chromatin spatial organization and transcriptional regulation are modulated in part by the incorporation of distinct histone variants, isoforms of canonical histones that differ slightly in amino acid sequence^64^. Aberrant deposition of these variants has been linked to pathological states including tumor initiation, therapy resistance, and metastasis^65,66^. Among these, the H3 variants (H3.1, H3.2, and H3.3), exhibit differential genomic distributions that contribute to the regulation of cellular identity and gene expression programs^67–70^. Notably, H3.1 is distinguished from H3.2 and H3.3 by a single unique cysteine residue at position 96 (Cys96)^71^ and is enriched in transcriptionally silent chromatin^72^. This cysteine confers susceptibility of H3.1 to oxidation by reactive oxygen species (ROS), such as hydrogen peroxide (H₂O₂), a property not shared by the other H3 variants^73,74^. Recent work leveraged a nuclear-localized D-amino acid oxidase (NLS-DAO) system to selectively elevate nuclear ROS (nROS) levels upon addition of D-alanine (D-Ala), enabling chromatin-specific redox manipulation without inducing oxidative DNA damage^75^. In breast cancer cells, oxidation of H3.1Cys96 under these conditions was shown to trigger its eviction from chromatin, thereby promoting decompaction of silenced domains and activation of epithelial-to-mesenchymal transition (EMT)-associated genes, as revealed by sequencing and electron microscopy analyses^75^.

To assess whether histone variant sensitivity to oxidative stress influences chromatin spatial organization, we carried out super-resolution imaging of histone H2B to visualize global chromatin structure (**Figure 5A, B**), in conjunction with O-SNAP analysis. We examined chromatin organization in MCF10A mammary epithelial cells co-expressing nuclear-targeted D-amino acid oxidase (NLS-DAO) and either the oxidation-sensitive histone variant H3.1 (**Figure 5A**) or the oxidation-insensitive variant H3.2 (**Figure 5B**). Cells were imaged under baseline conditions and following treatment with 10 nM D-Ala for 4 hours (**Figure 5A, B**). It was previously demonstrated that incubation of NLS-DAO enabled MCF10A cells treated with 10 nM D-Ala for 4 h elevates nuclear ROS (nROS) without inducing DNA damage^75^. Visual inspection of the super-resolution images of treated and untreated cells did not reveal obvious large-scale chromatin reorganization for either histone variant (**Figure 5A, B**), suggesting that any changes in spatial architecture are likely to be subtle and not visually apparent.

**Figure 5.**
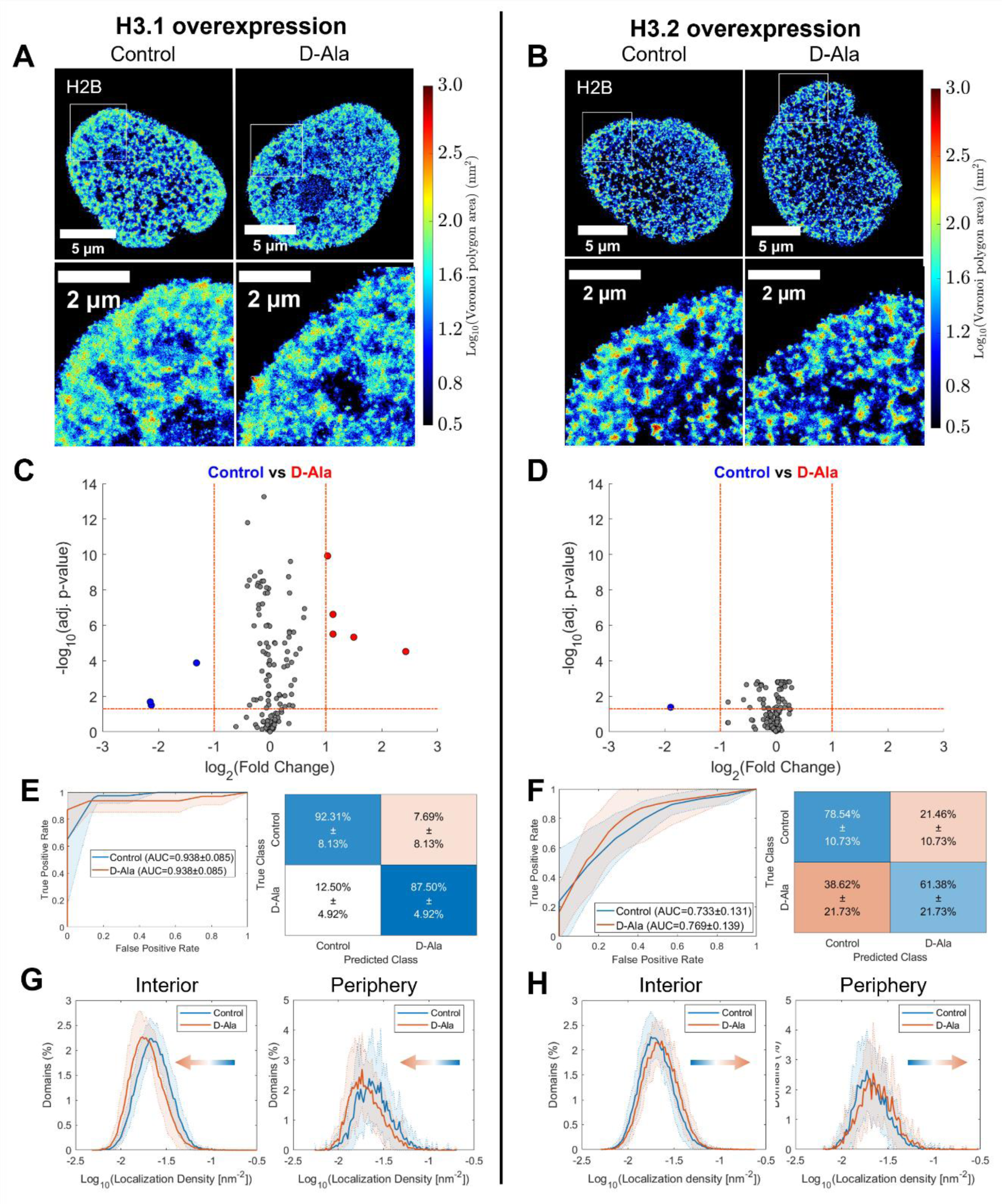
O-SNAP applied to the MCF10A/NLS-DAO H3 variant overexpression system suggests H3.1 is more sensitive to nROS compared to H3.2. (A) Representative Voronoi density map renderings of H2B STORM images of the MCF10A/NLS-DAO system with H3.1 overexpression in control cells and those treated with 10 nM D-Ala for 4 hours (N=39 nuclei for control and N=32 for D-Ala treated cells). (B) Representative Voronoi density map renderings of H2B STORM images of the MCF10A/NLS-DAO system with H3.2 overexpression in control cells and those treated with 10 nM D-Ala for 4 hours. (A, B) The color code indicates local chromatin compaction from low density in blue to high density in red. (C) Volcano plot visualizing fold changes in O-SNAP-generated features of H2B STORM data of MCF10A/NLS-DAO cells in control versus D-Ala treated cells. 5 features increase in value in the treatment group compared to the control, and 3 features increase in value in the control group (N=41 nuclei for control and N=29 for D-Ala treated cells). (D) Volcano plot visualizing fold changes in O-SNAP-generated features of H2B STORM data of MCF10A/NLS-DAO cells in control versus D-Ala treated cells. (C, D) The statistical significance was calculated using a two-sided t-test followed by Benjamini-Hochberg adjustment for multiple comparisons. (E) Classification results of a quadratic discriminant classification model trained to discriminate between control and D-Ala treated cells in the MCF10A/NLS-DAO, H3.1 overexpression system based on H2B STORM data with an overall validation accuracy of 90.19 ± 3.67%. (F) Classification results of a neural network model trained to discriminate between control and D-Ala treated cells in the MCF10A/NLS-DAO, H3.2 overexpression system based on H2B STORM data with an overall validation accuracy of 71.43 ± 7.95%. (E, F) The ROC curve (left) and a confusion matrix (right) indicate the performance for the control and D-Ala treatment conditions across the five folds. (G, H) The distribution of packing domain localization density of the MCF10A/NLS-DAO H3.1 (G) and H3.2 (H) overexpression system, where the solid line shows the percentage of packing domains that have a given localization density value for a given bin, averaged from over all nuclei of either the control or D-Ala treatment condition. The shaded region represents ±1 standard deviation interval from the mean for each bin.

We therefore turned to quantitative profiling using the O-SNAP pipeline. Volcano plot analysis revealed that eight chromatin features significantly changed upon D-Ala treatment in H3.1-overexpressing cells (**Figure 5C** and **Supplementary Table 10**), whereas only one feature passed the significance and fold-change thresholds in H3.2-overexpressing cells (**Figure 5D** and **Supplementary Table 11**). These findings are consistent with prior studies indicating that H3.1, due to its unique Cys96 residue, is selectively sensitive to nROS-induced oxidation^75^. Supporting this distinction, feature selection and classification analysis yielded a 90.19 ± 3.67% accuracy in distinguishing D-Ala-treated from untreated cells in the H3.1-overexpressing group (**Figure 5E**), compared to just 71.43 ± 7.95% accuracy in the H3.2-overexpressing group (**Figure 5F**).

Together, these results suggest that when H3.1 is the predominantly expressed histone variant, elevated nROS levels elicit detectable spatial remodeling of chromatin nanostructure, while H3.2-rich chromatin remains largely unaffected. Further analysis revealed that the features most impacted in the H3.1-overexpressing condition reflected decreased chromatin density in both internal and peripheral chromatin packing domains (nucleosome clutches; **Figure 5G**, **Supplementary Figure 8A**, and **Supplementary Table 10**), consistent with a shift toward chromatin decompaction. By contrast, the H3.2-overexpressing nuclei did not exhibit a decrease in compaction and the magnitude of the change that was present was less pronounced (**Figure 5H** and **Supplementary Figure 8B**). This observation aligns with earlier electron microscopy studies reporting similar chromatin opening in response to ROS-induced H3.1 eviction^75^.

These findings demonstrate that O-SNAP can sensitively detect subtle, histone variant-dependent changes in chromatin nanoscale architecture under redox conditions, reinforcing the emerging view that histone variant composition modulates chromatin’s structural plasticity in response to oxidative stress^75^ with important implications for understanding gene regulation in cancer, therapy resistance, and metastasis^76,77^.

### Spatial remodeling of chromatin in primary human chondrocytes cultured in vitro

Chondrocytes are the resident cells of articular cartilage, responsible for synthesizing and maintaining the extracellular matrix (ECM) that supports cartilage integrity and function. Injuries to articular cartilage are common and can lead to progressive erosion of joint surfaces, ultimately resulting in osteoarthritis^78–80^. One therapeutic approach for early-stage disease involves autologous cell-based repair, in which chondrocytes are harvested from the patient, expanded *in vitro*, and then re-implanted into the defect site using a biomaterial scaffold^81,82^. However, a major limitation of this strategy is the poor quality of the ECM produced by the re-implanted cells, a problem linked to the loss of chondrocyte phenotype during *in vitro* expansion^83,84^. Bulk RNA-sequencing studies have revealed widespread transcriptional dysregulation associated with passaging resulting from culture-induced chondrocyte dedifferentiation^85,86^.

To investigate whether the transcriptional dysregulation observed during chondrocyte expansion is accompanied by changes in chromatin spatial organization at the single-cell level, we performed super-resolution imaging of histone H2B in primary human chondrocytes isolated from two independent donors (**Figure 6A** and **Supplementary Figure 9A**). Cells were imaged at three timepoints during *in vitro* culture: Passage 0 (P0; baseline, no passaging), Passage 3 (P3; approximately 3 weeks of passaging), and Passage 6 (P6; approximately 6 weeks of passaging; **Figure 6A** and **Supplementary Figure 9A**). As with the histone variant datasets, visual inspection of the super-resolution images did not reveal obvious large-scale chromatin remodeling. We therefore applied the O-SNAP pipeline separately to cells from each donor to determine if it could detect changes in nanoscale chromatin organization.

**Figure 6.**
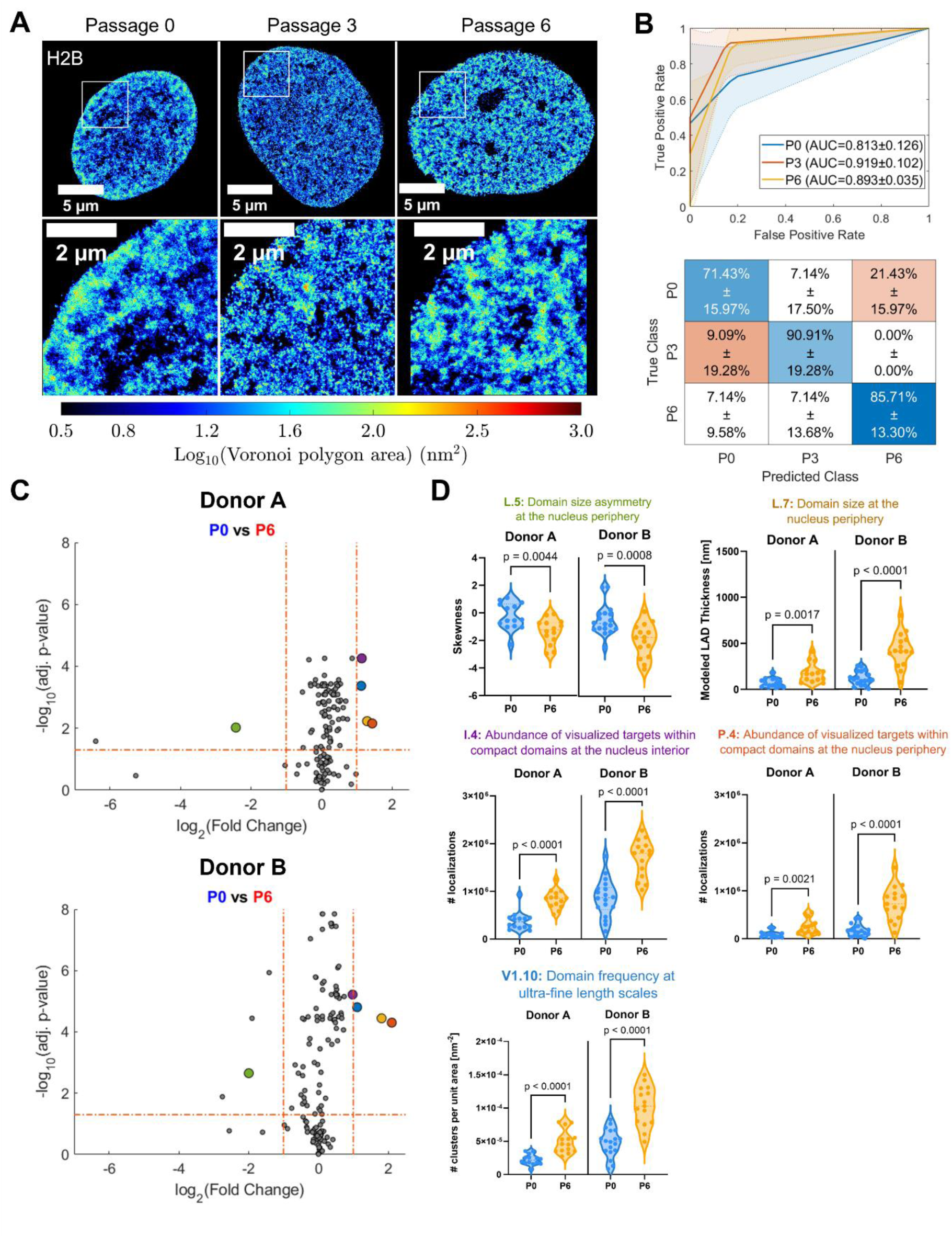
O-SNAP reveals consistent changes to chromatin organization in human chondrocytes over extended passaging. (A) Representative Voronoi density map renderings of H2B STORM images of human chondrocytes at Passage 0 (P0), Passage 3 (P3), and Passage 6 (P6) from Donor A (N=14 nuclei for P0, N=11 for P3, and N=14 for P6). The color code indicates local chromatin compaction from low density in blue to high density in red. (B) Classification results for a KNN model discriminating P0, P3, and P6 chondrocytes based on H2B STORM data across five training/test folds on Donor A. Top: The ROC curves across the five folds. Bottom: A confusion matrix demonstrating the classifier performance for each passage. The overall classification accuracy across the five folds is 82.14 ± 6.56%. (C) Volcano plot visualizing fold changes in O-SNAP-generated features of H2B STORM data of human chondrocytes at P0 compared to P6. Top: Donor A demonstrated 2 features significantly increase in value at P0 whereas 4 features increase in value at P6. Top: Donor A demonstrated 2 features significantly increased in value at P0 whereas 4 features increased in value at P6. Bottom: For Donor B, 4 features decrease in value for P0 and 3 increase at P6. The color-coded features (red, blue, yellow, green and purple) correspond to those that are the same between the two donors. The statistical significance was calculated using a two-sided t-test followed by Benjamini-Hochberg adjustment for multiple comparisons. (D) Violin plots of common features identified in (C) and **Supplementary Figure 9C** for P0 and P6 chondrocytes for each donor. Top row, left: The asymmetry of the domain size distribution at the nucleus periphery decreases in both donors. Top row, right: Model LAD thickness increases with passage. The number of localizations assigned to packing domains at both the nucleus interior (middle row, left) and periphery (middle row, right) increase by P6. Bottom row, left: The number of Voronoi clusters per unit area generated using a threshold of Voronoi area < 17.78 nm^2^ increases with passage.

Classification analysis using O-SNAP features achieved 82.14 ± 6.56% accuracy for Donor A (**Figure 5B**) and 82.89 ± 8.66% for Donor B (**Supplementary Figure 9B**) in distinguishing cells across the three passage conditions (P0, P3, and P6), indicating that chromatin spatial organization undergoes measurable and reproducible changes during *in vitro* expansion. To identify the specific features driving these changes, we performed volcano plot analyses comparing P0 and P6 cells (**Figure 5C**). Six features were significantly altered in Donor A and eight in Donor B, with five overlapping feature changes between the two donors, suggesting consistent patterns of chromatin remodeling associated with extended passaging. These shared features reflected increases in the size of chromatin packing domains and in the amount of histones within both peripheral and interior chromatin packing domains (**Figure 5C, D**, and **Supplementary Tables 12-13**), pointing toward the formation of larger chromatin domains indicative of increased heterochromatinization. In contrast, features that changed between P0 and P3 were largely donor-specific and did not persist at P6 (**Supplementary Figure 10A-D** and **Supplementary Tables 14-17**), suggesting that the intermediate P3 state may represent a transient and heterogeneous phase of chromatin reorganization, whereas the structural changes observed by P6 are more stable and convergent.

These results show that the O-SNAP pipeline can uncover subtle but persistent alterations to chromatin organization in chondrocytes during *in vitro* expansion. These new insights can provide a foundation for strategies aimed at preserving native chromatin states to support improved chondrocyte function for better therapeutic efficacy.

## Discussion

Here, we introduce O-SNAP, a comprehensive and interpretable analysis pipeline for profiling nuclear architecture from SMLM data. Our prior work with ECLiPSE demonstrated that dependence on rendered images introduces artifacts and information loss^44^; O-SNAP builds on this foundation by extending the analysis to the complex, multi-scale spatial features of the nucleus. Unlike previous machine learning approaches such as ASAP^87^ and AiNU^43^ that rely on rasterized image renderings, O-SNAP directly processes point cloud data, preserving the high-resolution spatial information critical for accurately characterizing chromatin organization. Moreover, O-SNAP incorporates downstream statistical tools inspired by RNA-Seq differential expression analysis, enhancing biological interpretability. This interpretability is a key advantage over black-box deep learning methods, as it allows for mechanistic insight into how specific chromatin features contribute to cell state identity and may guide targeted manipulation of chromatin states for therapeutic purposes.

Using O-SNAP, we validated known chromatin remodeling events in response to epigenetic perturbations (e.g., TSA treatment) and disease conditions (e.g., tendinosis), while also uncovering more subtle and heterogeneous alterations in chromatin spatial features. TSA treatment resulted in non-uniform decompaction of chromatin domains, particularly at the nuclear periphery, suggesting selective sensitivity of Lamina-Associated Domains (LADs). In heterokaryon reprogramming, O-SNAP revealed the persistence of a subpopulation of highly compact chromatin domains enriched in H3K27me3, indicative of facultative heterochromatin that may resist reprogramming. These observations open avenues for future studies using genomic methods such as ChIP-Seq to map LADs (via Lamin B1) or H3K27me3-enriched regions, enabling the translation of insights obtained from spatial chromatin architecture into genomic context to test specific hypothesis.

Furthermore, O-SNAP revealed differential sensitivity of chromatin to oxidative stress based on histone variant composition highlighting the utility of O-SNAP in detecting subtle metabolism-driven chromatin remodeling. Specifically, we found that chromatin containing the H3.1 variant was more susceptible to decompaction in response to elevated reactive oxygen species compared to H3.2-enriched chromatin, consistent with previous reports^75^. Beyond this, we applied O-SNAP to study dedifferentiation in cultured chondrocytes and observed progressive chromatin remodeling and heterochromatin formation with increasing passage number. Notably, persistent chromatin alterations emerged at Passage 6, whereas earlier passages exhibited more transient and heterogeneous changes. These observations suggest a critical temporal window prior to Passage 6 during which epigenetic interventions, such as treatments promoting chromatin decompaction (e.g., TSA), may be most effective in preventing unproductive chromatin remodeling. Future studies that systematically apply such interventions at defined time points, combined with super-resolution imaging and O-SNAP analysis, will be instrumental in testing this hypothesis.

Importantly, O-SNAP is not limited to global chromatin spatial organization analysis. As demonstrated in the heterokaryon reprogramming model, the pipeline can be readily adapted to study other nuclear features, including histone modifications. Its compatibility with different nuclear targets opens the door to analyzing the spatial organization of diverse nuclear structures, such as transcriptional machinery or nuclear bodies like speckles and nucleoli. Future work applying O-SNAP to multiplexed datasets^88^ will facilitate the assessment of how multiple nuclear features are remodeled in concert during cell state transitions and to identify a minimal yet informative set of nuclear markers for robust cell state classification. Such analyses could yield new insights into the coordination of nuclear architecture in development, disease, and reprogramming contexts.

O-SNAP incorporates strategies to address limited sample size and multiple hypothesis testing, such as cross-validation, dimensionality reduction, and statistical corrections. However, as for other supervised analysis methodologies, its performance relies on the quality of the input localization data, which is impacted by labeling density, segmentation accuracy, and the number of imaged nuclei for each phenotype. Inadequate quality of the input data may lead to false positives that do not reflect true biological patterns. Therefore, careful optimization of sample preparation and image acquisition is essential, as it underpins the reliability of any quantitative SMLM measurement. Both classification and pseudotime trajectory inference benefit from larger, balanced datasets, in which each phenotype is similarly represented. While SMLM imaging is currently time-intensive and low throughput, future advances in field-of-view expansion^89^ and automation, combined with O-SNAP analysis, may enable the detection of rare phenotypes. Nevertheless, users should exercise caution in such applications: although O-SNAP offers parameter adjustments to improve sensitivity in unbalanced datasets containing rare phenotypes, achieving balanced sampling is strongly recommended for robust results. Moreover, users should be familiar with the definitions and mathematical basis of the extracted features for careful interpretation of the feature outputs. For example, features based on distribution skewness may show disproportionately large fold changes in volcano plots, since a skewness value near zero represents a normal distribution, thereby amplifying deviations from this normality. Consulting both the feature definitions and associated violin plots is advised for accurate interpretation.

In summary, O-SNAP offers a powerful and versatile framework for analyzing both physiological and pathological transitions in nuclear architecture, providing mechanistic insights into the underlying chromatin dynamics and potentially enabling the rational design of interventions to modulate cell state transitions.

## Methods

### Cell culture and sample preparation

#### Human fibroblast sample preparation

Human BJ fibroblasts (ATCC, CRL-2522) were grown in media comprised of Dulbecco’s modified Eagle medium (DMEM; Sigma-Aldrich, #11965-084) supplemented with 10% v/v fetal bovine serum (HyClone, #SH30910.03), 1% v/v sodium pyruvate (100 mM; #11360-070, Gibco), 1% v/v antibiotic–antimycotic (Gibco, #15240-062), and 1% v/v GlutaMax (Gibco, #35050-061). Media was replaced every other day. For imaging, cells were seeded in Nunc™ Lab-Tek™ II 8-well chambered coverglass (ThermoFisher, #155409PK). For hyperacetylation STORM experiments, cells were treated with 300 nM of Trichostatin (TSA; Sigma-Aldrich, #T8552) in complete growth medium for 24-hr prior to experiments.

Cells were fixed in PBS with 4% v/v paraformaldehyde (Electron Microscopy Sciences, #15710) warmed to 37 °C for 10 min at 25 °C. Fixed cells were washed three times with PBS and permeabilized with 0.1% Triton X-100 (Fisher Scientific, #BP151-100) in PBS. Cells were incubated in blocking buffer containing 10% w/v bovine serum albumin (BSA; Fisher Scientific, #BP1600-100) and Triton X-100 at 0.2% v/v for 1 hour at 25 °C. Rabbit anti-histone H2B primary antibodies (Invitrogen, #PA5-115361) targeting were diluted in the blocking buffer at 1:50 for 15-18h at 4°C. Cells were rinsed three times in a wash buffer of 2% w/v BSA in 0.025% v/v Triton X-100 in PBS with gentle agitation, followed by incubation with anti-Rabbit secondary antibodies conjugated to Alexa Fluor 647 (Invitrogen, #A31573) diluted in blocking buffer at 1:50 for one hour at 25 °C. Cells were washed again three times in wash buffer at 25 °C.

#### Human tenocyte sample preparation

The human tenocyte SMLM data was originally generated and described in detail in Heo et al^32^; for a more in-depth protocol, readers are directed to the original publication. Briefly, human tendons were obtained from patients undergoing revision amputation for traumatic hand injury who were young (36 and 42 yr) or had been diagnosed with tendinosis (35 and 39 yr) using IRB-approved protocols (#13D.238). Tissues (2×2 mm) were digested in collagenase solution and centrifuged. Cell pellets were resuspended and grown in media containing high glucose DMEM medium, 10% penicillin–streptomycin, L-glutamine and 10% fetal bovine serum (FBS).

For STORM imaging, cells were fixed in methanol–ethanol (1:1) at −20 °C for 6 min, followed by incubation in blocking buffer containing 10% w/v BSA (Sigma) in PBS for 1 hour. The samples were incubated overnight with rabbit anti-H2B (Proteintech, #15857-1-AP) at 1:50; rabbit anti-H3K4me3 (Thermo, #MA5-11199) at 1:100; or rabbit anti-H3K27me3 (1:100; Thermo, #PA5-31817) at 4 °C. After repeated washing in PBS, the samples were incubated with secondary antibodies custom labelled with activator–reporter dye pairs (Alexa Fluor 405–Alexa Fluor 647; Invitrogen, #A30000 and #A20006).

#### Heterokaryon sample preparation

The heterokaryon SMLM data was originally generated and described in detail in Martinez-Sarmiento et al^31^; for a more in-depth protocol, readers are directed to the original publication. Primary human BJ fibroblasts (ATCC, CRL-2522) were maintained in DMEM 1X (Gibco, # 11965– 084) containing 10% of Fetal Bovine Serum (Cytiva, #SH30071.03), Penicillin/Streptomycin 1X (Gibco, #15240–062) and Sodium Pyruvate 1X (Gibco, #11360–070). Media was replaced every other day. *Tcf3^-/-^* mouse embryonic stem cells (mESCs)^90^ were cultured in KnockOut DMEM, 1X (Gibco, #10829–018), containing 15% FBS, Penicillin/ Streptomycin 1X, Sodium Pyruvate 1X, GlutaMAX 1X (Gibco, #35050–061), MEM amino acids 1X (Gibco, #11140–050), 2-Mercaptoethanol 50 mM (Gibco, #31350–010) and 106 units/mL of LIF (Leukemia Inhibitory Factor, EMD Millipore). Media was changed daily.

To generate heterokaryons, 5-10×10^6^ hFbs and a similar number of number of mESCs *Tcf3^-/-^* were harvested and mixed at a ratio of 1:1 using 1-2 mL of polyethylene glycol 1500 (PEG; Roche, #10783641001) at 37°C. Cells were centrifuged resuspended in PBS 1X (Gibco, # 14190–136) with 10% FBS and incubated with Phycoerythrin (BD Pharmingen, #561970) and Alexa Fluor 647-conjugated antibodies against the mutually exclusive surface antigens CD90 (BD Pharmingen, #17-0909-42) and E-Cadherin (BioLegend, # 147307) as 0.5 µL of antibody per mL per every 10×10^6^ cells. Cells were sorted with an Influx B sorter and double-positive (PE-CD90^+^, Alexa Fluor 647-E-Cadherin^+^) cells were considered to have undergone fusion. Cells were plated into Nunc™ Lab-Tek™ II 8-well chambered coverglass (ThermoFisher, #155409PK) coated with Laminin (Sigma-Aldrich, #114956-81-9) at a density of 3×10^4^ cells/cm^2^. Single positive cells were sorted as controls. Cells were cultured in mESC conditions for 48-hr post fusion. Media was changed at 24-hr.

Cells were washed three times with PBS 1X and fixed using an osmotically-balanced paraformaldehyde 4% w/v solution (PFA; Electron Microscopy Sciences, #1570) during 10 min at RT and washed with PBS 1X. Cells were washed with PBS 1X and permeabilized with Triton X-100 (Fisher BioReagents, #BP151-100) 0.1% v/v for 10 min, RT and washed with PBS 1X. Cells were then incubated in blocking buffer consisting of bovine serum albumin (BSA; Fisher, #BP1600-100), 10% w/v (Fisher, #BP1600-100) and Triton X-100 0.0025% in PBS 1X, for 1 h at RT. Primary antibodies were diluted in blocking buffer and used as follows: H2B (Proteintech, #15857-1-AP) 1:25; H3K27me3 (Thermo Fisher Scientific; #MA5-11198) 1:100; and H3K9ac (Thermo Fisher Scientific, #MA5-11195) 1:50; and incubated 12h–15 h at 4°C in a humidified chamber. Cells were washed three times in agitation at RT in washing buffer containing 2% w/v BSA and 0.01% v/v Triton X-100 in PBS 1X. Secondary antibody was added in blocking buffer containing DAPI (Thermo Scientific, #62248) at 5 mg/mL during 1 h at RT and then washed with PBS 1X. For STORM imaging, in-house made secondary antibodies conjugated with Alexa Fluor 405–647 organic dye pairs were used at the same dilution as primary antibodies. DAPI and Lamin/AC staining patterns were used to distinguish the human and mouse nucleus in heterokaryons during imaging.

#### MCF10A sample preparation

MCF10A cells (ATCC, CRL-10317) were cultured in DMEM/F12 (no phenol red) medium (Gibco, #21041025) supplemented with 10% FBS (Sigma Aldrich, #F2442), 20 ng/mL EGF (Sigma Aldrich, #SRP3027), 10 μg/mL insulin (Sigma Aldrich, #I3536), 0.5 μg/mL hydrocortisone (Sigma Aldrich, #H0135), 0.1 μg/mL cholera toxin (Sigma Aldrich, #C8052), and 1% penicillin-streptomycin (Sigma Aldrich, # P4333). For H3.1 and H3.2 overexpression in MCF10A cells, inducible TET-ON H3.1-FLAG and TET-ON H3.2-FLAG plasmids were obtained from cloning H3.1, H3.2 and FLAG tag sequences into pLVX-Tet3G blasticidin vector (Addgene, #128061). NLS-DAO plasmid was obtained from cloning the c-myc sequence into the D-Amino acid oxidase (DAO) vector (a generous gift from Thomas Michel, Harvard University) with a geneticin selection marker. MCF10 cells expressing NLS-DAO and either H3.1-FLAG or H3.2-FLAG variants were treated with D-Ala by replacing medium for 4h with new, fresh medium containing 10 nM D-Ala. Control cells received fresh medium only.

For imaging, cells were seeded in Nunc™ Lab-Tek™ II 8-well chambered coverglass (ThermoFisher, #155409PK). Cells were fixed in PBS 1X with 4% v/v paraformaldehyde (Electron Microscopy Sciences, #15710) warmed to 37 °C for 10 min at 25 °C. Fixed cells were washed three times with PBS 1X and permeabilized with 0.1% Triton (Fisher Scientific, #BP151-100) in PBS. Cells were incubated in blocking buffer containing 10% w/v bovine serum albumin (BSA; Fisher Scientific, #BP1600-100) and Triton X-100 at 0.0025% for 1 hour at 25 °C. Rabbit anti-histone H2B primary antibodies (Invitrogen, #PA5-115361) were diluted in the blocking buffer at 1:50 for 15h at 4°C in a humidified chamber. Cells were rinsed three times in a wash buffer of 2% w/v BSA in 0.01% v/v Triton X-100 in PBS 1X with moderate agitation, followed by incubation with anti-Rabbit secondary antibodies conjugated to Alexa Fluor 647 (Invitrogen, #A31573) diluted in blocking buffer at 1:50 for one hour at 25 °C. Cells were washed again three times in PBS 1X at 25 °C.

#### Human chondrocyte sample preparation

Primary human articular chondrocytes were isolated from cartilage obtained from two male donors (ages 46 and 55; LifeLink Tissue Bank) using enzymatic digestion. Briefly, cartilage fragments were incubated with LiberaseTM (a blend of highly purified Collagenase I, Collagenase II, and Thermolysin; Sigma Roche, SKU: #5401119001) at 37°C with gentle rocking for 18 hours. The resulting cell suspension was filtered, centrifuged, and resuspended in basal growth medium composed of Dulbecco’s Modified Eagle Medium (DMEM; ThermoFisher, #11965118) supplemented with 10% fetal bovine serum (FBS; R&D Systems, #S11150) and 1% penicillin-streptomycin (Corning, #30-002-CI). Cells were cultured at 37°C in a humidified incubator with 5% CO₂ and passaged upon reaching ∼70% confluency. For expansion, cells were seeded at a density of 9000 cells/cm^2^, with media changes every 2-3 days.

Passage 0, 3 and 6 chondrocytes were seeded in Nunc™ Lab-Tek™ II 8-well chambered coverglass (ThermoFisher, #155409PK) and cultured for 2 days. Fixation was performed using a methanol-ethanol (1:1) mixture at -20°C for 6 minutes, followed by three washes with PBS. Blocking was carried out with BlockAid solution (ThermoFisher, #B10710) for 1 hour at room temperature to reduce nonspecific binding. Cells were incubated overnight at 4°C with rabbit anti-histone H2B antibody (1:50; Thermo Scientific, PA5115361), followed by three PBS washes. Secondary antibodies were labeled with activator-reporter dye pairs (Alexa Fluor 405-Alexa Fluor 647; Invitrogen, #A30000 and #A20006) and incubated for 1 hour at room temperature. Cells were washed three times in agitation at RT in washing buffer containing 2% w/v BSA and 0.01% v/v Triton X-100 X-100 in PBS 1X. Secondary antibody was added in blocking buffer containing DAPI (Thermo Scientific, #62248) at 5 mg/mL during 1 h at RT and then washed with PBS 1X. For STORM imaging, in-house made secondary antibodies conjugated with Alexa Fluor 405–647 organic dye pairs were used at the same dilution as primary antibodies. DAPI and Lamin/AC staining patterns were used to distinguish the human and mouse nucleus in heterokaryons during imaging.

### STORM imaging and preprocessing

All images were acquired on an ONI Nanoimager (Oxford Nanoimaging microscope) using NimOS Nanoimager Software (Version: 1.19.4). The microscope is equipped with a 100×, NA 1.45, oil immersion lens (Olympus) and four lasers: 405, 488, 561 and 640 nm lasers, two channels: the first with 498–551 nm and 576–620 nm band-pass filters (Channel 0) and the second with 666–705–839 nm band-pass filters (Channel 1), and a Hamamatsu Flash 4 V3 scientific complementary metal-oxide-semiconductor (sCMOS) camera with a pixel size of 117 nm and field of view of 50 µm x 80 µm. Nuclei were randomly selected and imaged using Highly Inclined and Laminated Optical sheet illumination (HILO).

Cells were imaged in an imaging buffer of 0.1 M cysteamine (MEA; stock, 77 mg/mL #30070-10G, Sigma-Aldrich in 360 mM HCl; #A508-P500 Fisher), 5% w/v glucose (#A16828, Alfa Aesar), and 1% glucose oxidase (GLOX) solution. The GLOX solution was made using 14 mg Glucose Oxidase (#G2133, Sigma-Aldrich) in 50 µL of catalase (20 mg/mL; #106810, Roche Applied Science) in 200 µL of 10 mM Tris (#15568025, Invitrogen) at pH 8.0 with 50 mM NaCl (#S217-500 Fisher). The imaging buffer was replaced every 90 minutes. Images were acquired with a 15 ms exposure time for 30,000 frames at 30 °C with constant, high intensity laser power excited at 647 nm. To account for the high density of H2B signal in the nucleus, the sample was first pre-bleached for 1 minute with high laser power prior to image acquisition.

Following acquisition of the raw images, STORM localizations were generated and drift-corrected using the NimOS software with a localization precision of 30 nm. Additional filtering thresholds are listed in **Supplementary Table 18**. Localization data was exported from the NimOS software as CSV files. Using a custom-made MATLAB analysis software^91^, localizations were manually cropped to the shape of the nucleus. The segmented localization coordinate data of chromatin targets were then used as input for the O-SNAP analysis pipeline.

### O-SNAP analysis

#### Data requirements

O-SNAP was implemented in MATLAB (MathWorks). It accepts an array of 2-D coordinates and was developed to study localizations with variable spatial distribution about the nucleus. The data should be already segmented to the boundary of the nucleus. O-SNAP assumes the raw data is in pixels and prompts users for a conversion factor to nm. Beyond characterization of the nucleus features, the user must label each nucleus localization set with its associated class (e.g. treatment, timepoint, cell type) if they aim to utilize the classification features.

#### O-SNAP feature extraction

##### Nucleus morphometrics

From SMLM image data, features describing the 2-D shape of the nucleus can be calculated and these morphology-related features are prefixed in the O-SNAP shorthand system with “M.” The nucleus boundary is calculated on the localizations by using the MATLAB *boundary()* function to identify the outermost localizations that form the outline of the nucleus and using *smoothdata()* to smooth the boundary with a moving average.

With the nucleus boundary defined, several standard morphometric features can be calculated, such as the nucleus perimeter (M.26). The nucleus area is calculated by using the *polyarea()* function to measure the area defined by the boundary. The nucleus radius is calculated by measuring the radius of a circle with the same area as the nucleus (M.25). The major and minor axes of the nucleus are determined by performing Principal Component Analysis (PCA) on the boundary localizations, which rotates the point cloud so that the axis with the largest variation in localization position is aligned along the x-axis. The major axis (M.21) is the longest length along the first component, the minor axis (M.23) is the length along the second component, and the aspect ratio (M.1) is the ratio of the major to minor axis lengths. Features, such as boundary bending energy (M.3) and boundary elastic energy (M.12) are also derived by quantifying the curvature of the boundary. The boundary can be used to create a convex hull, the smallest convex polygon that contains the original shape that then provides additional features that compare the properties of the original shape to those of the convex hull, such as convexity (M.8). A detailed explanation for all morphometric features is described in Rows 1-28 of **Supplementary Table 3**.

##### Voronoi localization segmentation

To quantitatively measure both local and global spatial distribution of localizations, a previously described method of Voronoi tessellation was used to segment the coordinate data^45,46^. The Voronoi analysis was conducted using the custom MATLAB software (R2024b 24.2.0.2712019 64-bit, win64). This procedure assigns a polygonal cell for each localization whose area can then be quantified and enables clustering. Briefly, a Delaunay triangulation was generated using the built-in *delauneyTriangulation()* function, which serves as a basis for the built-in *voronoiDiagram()* function to segment the coordinate space into Voronoi cells, returning a list of vertices and regions corresponding to one polygon for each localization. The built-in MATLAB *polyarea()* function then determined the area for every localization’s Voronoi cell. To quantify chromatin compaction, the Voronoi density was then defined as the inverse of the Voronoi area, where higher values of Voronoi density correspond to localizations in regions of higher chromatin compaction. As a normalization to account for variation in labeling between nuclei, the reduced Voronoi density was calculated, where for each nucleus, the Voronoi area distribution is divided by the mean Voronoi area value, resulting in the reduced Voronoi area distribution. The inverse of the reduced Voronoi areas is the reduced Voronoi density distribution. This information was visually represented by color-coding the localization renderings from blue to red based on the value of the log value of the Voronoi density.

Features related to global localization density are prefixed in the O-SNAP shorthand system with “G.” The mean, standard deviation, and skewness were derived from the distribution of both the Voronoi density and reduced Voronoi density to include as O-SNAP features (G.5, G.7, and G.6, respectively). Only the mean of the reduced Voronoi density is included as a feature (G.10), as the standard deviation and skewness produce identical values to that of the non-normalized Voronoi density distribution. The 40^th^ percentile value of the Voronoi (G.3) and reduced Voronoi density (G.8) was used as an O-SNAP feature to represent sparse chromatin (euchromatin) while the 70^th^ percentile (G.4 and G.9, respectively) were used as a feature to represent dense chromatin (heterochromatin). In addition to these values, the total number of localizations (G.1) and the number of localizations per unit area (G.2) are also included as features. These features are also described in Rows 29-38 of **Supplementary Table 3**.

##### Nucleus periphery and interior segmentation

In addition to features that describe changes occurring to chromatin organization globally throughout the nucleus, it was of interest to understand how chromatin organization at the periphery of the nucleus differed from the nucleus interior. It was thus necessary to divide these two regions and designate separate O-SNAP features for each region. Features related to the nucleus interior are prefixed in the shorthand system with “I,” and features related to the nucleus periphery are prefixed with “P.” Localizations and high-density packing domains are characterized separately for these two regions (see **DBSCAN analysis** under **Methods** for details on analysis of high-density packing domains).

There are two ways in which the distinction between nucleus periphery and interior are implemented. One approach based on Heo et al^32^ defines the nucleus periphery as the outer 15% of the nucleus area and the nucleus interior as the inner 85%. This cutoff is used for features I.1 (localization density per unit area at the nucleus interior), I.2 (packing domain density per unit area at the nucleus interior), P.1 (localization density per unit area at the nucleus periphery), P.2 (packing domain density per unit area at the nucleus interior), P.16 (the ratio of P.1:I.1), and P.17 (the ratio of P.2:I.2). These features are also described in Rows 39-40, 57-58, and 72-73 of **Supplementary Table 3**. A second, more stringent cutoff that defines the periphery as the outermost 5% of the nucleus area was used to characterize the thickness and length of putative Lamin Associated Domains (LADs) and is based on the analysis pipeline developed for Kant et al^51^, which is further described in **Modeled LAD analysis** under **Methods**.

##### DBSCAN analysis

O-SNAP leverages two distinct clustering segmentation approaches to characterize multiple facets of chromatin organization throughout the nucleus. This section describes the first of these approaches, which characterizes dense regions of chromatin, or packing domains, based on a previously developed analysis pipeline^51^. The other approach is described in the following section, **Voronoi clustering analysis**.

First, localizations were filtered for those with Voronoi densities above the 70^th^ percentile value from all nuclei within a system. The DBSCAN algorithm segmented the filtered localizations into packing domains (or clutches) representing the highest density regions of chromatin. DBSCAN was implemented with two parameters: (1) ε, the threshold for the neighborhood search radius, here used with a value of 20 nm, and (2) the minimum number of neighbors in a cluster for a given core point, here set to 3. The resulting clusters are then filtered for those with greater than 35 localizations per cluster.

The segmented clusters were further separated into two categories: peripheral and interior clusters, where the peripheral clusters are within 5% of the nucleus radius from the nucleus boundary and the remaining clusters are considered as interior packing domains. The distribution and quantity of these domains are described with feature I.3 (number of localizations attributed to packing domains at the nucleus interior), I.4 (number of packing domains at the nucleus interior), P.3 (number of localizations attributed to packing domains at the nucleus periphery), and P.4 (number of packing domains at the nucleus periphery), which can be found in Rows 41-42 and 59-60 of **Supplementary Table 3**.

The properties of the domains at the periphery and interior were then separately characterized, where domains were measured for their size (i.e. equivalent radius), compactness (i.e. localization density per unit area), dispersion (i.e. radius of gyration), and number of localizations per domain. Additionally, for packing domains at the nucleus interior, the spacing between neighboring packing domains were calculated. For each property, the domains generated a distribution of values, which were log-transformed to obtain a distribution more closely resembling a normal distribution. The mean value, standard deviation, and skewness across these distributions for each were used as O-SNAP features for each nucleus. Features I.5-I.18 and P.5-P.15 represent these values, which can be found in Rows 43-56 and 61-71 of **Supplementary Table 3**.

##### Voronoi clustering analysis

The following section describes the second of the two clustering approaches implemented in O-SNAP. This approach achieves characterization of clusters at multiple length scales using Voronoi clustering.

The algorithm takes a parameter that defines the upper threshold for localizations to cluster based on their Voronoi area. The algorithm then clusters these filtered localizations by forming groups of contiguous Voronoi cells. Clusters with fewer than a minimum number of localizations (default = 5) are discarded. For this paper, we selected five area thresholds ranging from 17.78 nm^2^ to 100.0 nm^2^ evenly across a log scale to achieve broad coverage of cluster sets at different length scales (17.78 nm^2^, 32.61 nm^2^, 56.23 nm^2^, and 100.0 nm^2^), where smaller area thresholds lead to smaller, more dispersed (i.e., are more frequent and, generally, more evenly spaced out throughout the nucleus) clusters (**Supplementary Figure 1A**). Increasing the area threshold leads to segmentation of more contiguous chromatin domains at larger length scales. Features derived from these clusters are prefixed in the shorthand system with “V#,” where “#” ranges from 1-4 that correspond to the Voronoi clustering thresholds. Clusters were measured for their size (i.e. equivalent radius), compactness (i.e. localization density per unit area), and dispersion (i.e. radius of gyration). The mean value, standard deviation, and skewness across these distributions for each were used as O-SNAP features for each nucleus. The number of clusters per unit area was also measured as a feature. These features are listed in Rows 105-144 of **Supplementary Table 3**.

##### Model LAD analysis

The packing domains at the nuclear periphery (within 5% of nuclear boundary) produced by DBSCAN were used to create models of the unique chromatin pattern at the nucleus periphery as previously described in Kant et al^51^. Briefly, about fifty quadrilaterals, or “model LAD domains,” were created such that one side ran tangent to the nucleus boundary, creating a rough perimeter of the nucleus. The height of each quadrilateral perpendicular to this tangent was dependent on the size of the packing domain at the periphery. This set of quadrilaterals could then be assessed for additional features describing the organization of chromatin at the nucleus periphery. The mean, standard deviation, and skewness of the distribution of the quadrilateral side lengths both tangent and perpendicular to the nucleus boundary, as well as the value of the total modeled LAD area divided by the total tangent side lengths, were used as O-SNAP features. These features have the prefix “L” in the shorthand system and appear in Rows 74-80 of **Supplementary Table 3**.

##### Radial density analysis

Chromatin organization with respect to a given radial distance from the centroid of the nucleus was evaluated by modeling the nucleus as an ellipse and measuring the density of localizations or packing domains contained within each ring. These were constructed by dividing the major and minor axes lengths into ten equal segments and fitting ellipses for each major/minor axes segment pair. These ellipses define the ten regions for radial analysis. Because of the irregular shape of nuclei, localizations may lie beyond the outermost ellipse. These localizations are assigned to the outermost ring such that they can still be considered. While this approach results in ring areas varying on the nucleus area, this guarantees the same number of rings, and thus features, across all nuclei. For each ring, features are obtained by measuring either the localization density (features with prefix “R” in the shorthand system, features R.1-R.10) or the number of packing domains per unit area (features with prefix “RD” in the shorthand system, features RD.1-RD.10; **Supplementary Figure 1C**). Following this, the average change in these densities over the innermost eight rings was calculated, normalized over the length of the major or minor axis. For localization density, this corresponds to features R.11 (the change in localization density radially along the major axis) and R.12 (the change in localization density radially along the minor axis). Similarly, for the number of packing domains per unit area, this corresponds to features RD.11 (the change in packing domain density radially along the major axis) and RD.12 (the change in packing domain density radially along the minor axis). These features appear in Rows 81-104 in **Supplementary Table 3**.

#### Volcano analysis

The feature distributions were compared between each pair of conditions available for a given SMLM data set. The difference in mean values were compared between each pair, without outlier removal by default. The two-sample *t*-test was used to determine evaluate the difference between the distributions. To account for multiple hypothesis testing, p-values were then adjusted, where the default is set to the Benjamini-Hochberg procedure^92,93^.

#### Feature set enrichment analysis

Feature set enrichment analysis (FSEA) adapts the Gene Set Enrichment Analysis (GSEA) method introduced in Subramanian et al.^52^. Here, FSEA aims to achieve a similar goal with O-SNAP features rather than gene expression values. Features are ranked with an established ranking metric. The Enrichment Score (*ES*) quantifies the degree to which a given feature set differs from a uniform, null distribution. In addition, a test statistic is calculated that describes the level of significance of the feature set enrichment, which is adjusted to account for multiple hypothesis testing and differences in the number of features between sets.

##### Input

FSEA operates on a set of samples with *N* features (formerly *N* genes in GSEA). In O-SNAP, each sample represents a nucleus that is one of two possible phenotypes of interest, *A* or *B*. All *N* features are ranked to form *L* = {*f*_1_, … *f*_*N*_} based on a metric, *r*(*f*_*j*_) = *r*_*j*_, that relates the feature value to the observed phenotype. FSEA also requires a group of *M* feature sets, *S*, that represent the categorization of features into related trends, where each feature should appear exactly once across all the sets (**Supplementary Figure 2A**, **Supplementary Table 17**).

##### Calculation of enrichment score (ES)

An *ES* is calculated for each feature set *S* containing *N*_*H*_ features. Traversing the ranking *L* = {*f*_1_, … *f*_*N*_}, values for the fraction of features in *S*(“hits”), weighted by their correlation (see **Equation 1**), and *S*(“misses”) (**Equation 2**), are calculated at each position (**Supplementary Figure 2B**). The exponent *p* controls the weight of the step.

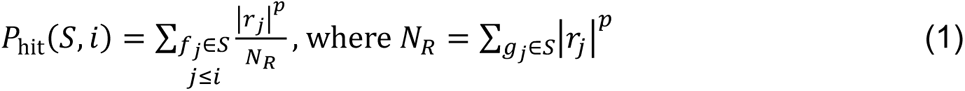

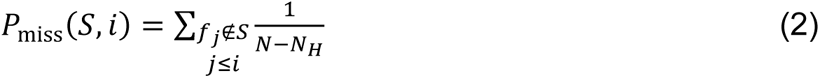

The *ES* is the maximum deviation of *P*_hit_ − *P*_miss_ from zero over all values of *i* (**Supplementary Figure 2C**). The value of *ES*(*S*) will be higher for sets whose features concentrate either at the top or bottom of *L*. Note that O-SNAP incorporates a correction absent from traditional GSEA that modifies the position of features in *L* whose value increase opposes that of the trend described in the feature set. Thus, the relationship between *ES*(*S*) and the distribution of features along *L* still holds. For *p* = 0, *ES*(*S*) reduces to the Kolmogorov-Smirnov statistic, and when *p* = 1 (the default value used in O-SNAP), the features in *S* are weighted by their correlation to phenotype *A* or *B*, normalized by the total sum of correlations over all genes in *S*. The latter option, according to Subramanian et al, results in more gene sets with less-than-perfect coherence to score well.

##### Estimating significance

Significance is measured by comparing *ES* to a set of scores, *ES*_NULL_, derived from a null distribution of randomly assigned phenotypes to the original feature data. Note that only the labels of the samples are shuffled to preserve existing correlations between features within samples. A new ranking *L* is created using the shuffled data and a corresponding value *ES*(*S*) is calculated. This permutation and recalculation of *ES*_NULL_ is performed, by default 1,000 times, creating a distribution of *ES*_NULL_ scores. A nominal *p*-value for *S* is determined by calculating the proportion of *ES*_NULL_ scores less than the calculated *ES* of the original data set, if *ES*(*S*) is positive, and the proportion greater than *ES*, if *ES*(*S*) is negative.

##### Multiple hypothesis testing: adjusting for variation in gene set size

To compare *ES* between multiple feature sets, the scores must be further normalized, and the significance testing should be corrected for multiple hypothesis testing. For each *S* and 1,000 fixed permutations π of the phenotype labels, a new ranking *L* and associated *ES*(*S*, *π*) are calculated. To adjust for differences in feature set size (i.e. the number of features within each set), the positive and negative values of *ES*(*S*) and *ES*(*S*, *π*) are rescaled separately, where the positive and negative scores are divided by the respective mean of *ES*(*S*, *π*) to obtain the normalized scores, *NES*(*S*) and *NES*(*S*, *π*) (**Supplementary Figure 2D**). Further detail regarding the reasoning behind this approach is discussed in more detail in Subramanian et al^52^.

After normalized scores are obtained, the False Discovery Rate (FDR) is computed. The ratio of false positives to the total number of sets is controlled by fixing the level of significance separately for the set of *NES*(*S*) and *NES*(*S*, *π*) with either positive or negative values. To do this, a null distribution is formed by considering all values of *NES*(*S*, *π*) for all *S* and *π*. Then, for a given *S*, in the case where *NES*(*S*) = *NES* ∗ ≥ 0 (i.e. the positive *NES* case), the *FDR* is calculated as the ratio of the percentage of all (*S*, *π*) with *NES*(*S*, *π*) ≥ *NES*^∗^ divided by the percentage of observed *S* with *NES*(*S*) ≥ 0 whose *NES*(*S*) ≥ *NES*^∗^. This procedure was performed similarly for *NES*(*S*) = *NES*^∗^ ≤ 0, where the FDR is instead the ratio of the percentage of all (*S*, *π*) with *NES*(*S*, *π*) ≤ *NES*^∗^ divided by the percentage of observed *S* with *NES*(*S*) ≤ 0 whose *NES*(*S*) ≤ *NES*^∗^. The final plot displays feature sets with largest *NES* values, where the color of the markers indicates the FDR q-values and the size of the marker is the number of features contained in the set (**Supplementary Figure 2E**).

#### Pseudotimeline analysis

Pseudotimeline analysis was performed by utilizing the R package collection, *dynverse*, which enables the exploration and implementation of trajectory inference (TI) methods^54^.

O-SNAP feature sets derived from SMLM data originally produced in Martinez-Sarmiento et al^31^ were adapted as input for trajectory inference analysis. The feature data was exported from MATLAB as a CSV and imported into R. Images from the hFb nuclei at 0-, 6-, 24-, and 48-hr post-fusion were considered for the pseudotime analysis, and the control mESC nucleus was excluded from the pseudotime analysis. The feature data was then z-score normalized and used as the expression data. Because the *dynverse* wrapper requires sequencing count data in the input dataset, this array was set to a single, trivial value of 0.5. Prior information on the sample phenotype labels, expected start and end phenotypes, and expected transitions between phenotypes (e.g. 0- to 6-hr, 6-hr to 24-hr, and 24-hr to 48-hr) was also provided since the reprogramming behavior of the somatic heterokaryon nucleus is expected to be continuous and not form multiple, distinct endpoint states. The prior information also served to improve the stability and simplicity of the inferred trajectory.

A shortlist of trajectory methods were obtained from *dynverse* based on dataset-dependent parameters, such as cell count and the expected trajectory topology (see **Supplementary Table 5**). Each of these candidate methods analyzed the processed feature data to create an initial trajectory, where samples are placed along a topology and assigned a value that quantifies the progression between cell transitions. If the samples form distinct subpopulations with time, the trajectory’s topology may feature branches that represent cell state decisions. To facilitate interpretation, the pseudotime calculated from the trajectory analysis was used to project the samples onto a new linear trajectory and samples on extraneous branches deviating from the main progression were omitted in subsequent analysis. The pseudotimes for every cell state were binned to evaluate to degree each trajectory method could separate the samples across pseudotime in a manner that reflected their ground truth labels.

Once the best performing method was determined, in this case Slingshot^57^, the most important features informing the trajectory were identified from the H3K27me3 data using functions provided in *dynverse*. With a heatmap, the values of the top three important features were plotted against pseudotime and compared between the H3K27me3 signal and global chromatin.

#### Feature selection and principal component analysis

In this work, O-SNAP data is used to train and test classification models designed to distinguish different cell states of interest to (1) identify which features and (2) evaluate the extent these features are sufficient to distinguish between phenotypes.

To rigorously evaluate the classification performance from a limited sample size, the data is split into five folds where a random 80% of the sample are used for model training and the remaining 20% is reserved to test for model performance. The data is split such that each fold contains a different subset of the training data with at least one member of each class. Within each fold, the training data is z-score normalized (**Supplementary Figure 4A**).

Feature selection is then performed, using the Minimum Redundancy – Maximum Relevance (MRMR) algorithm^94^, which was first introduced to identify features of interest from gene expression data. This approach aims to identify which subset of features maximizes the relevance to the phenotype labels while also minimizing the redundancy of the set that represents information shared between the selected features (e.g. features that have a strong correlation). Details on the MRMR procedure are fully described both in the original publication^94^ as well as the MATLAB documentation. Once features are ranked on the training set, a decision must be made on the number of features to include in the final set. This cutoff was automatically determined by examining the top 12 MRMR-ranked features and locating either the “knee” point in the MRMR scores of the ranked features^95^ or the feature at which there was the largest drop in MRMR scores occurs. Whichever of these options results in the smaller set of features is taken as the final selected set for that fold (**Supplementary Figure 4B**).

Although the feature selection occurs separately for each fold, it is difficult to interpret five separate feature sets. To address this, O-SNAP aggregates the feature sets together by summing the MRMR scores together from each fold to obtain an aggregated ranking. The same approach for finding a cutoff number of features is used on the aggregate feature selection (**Supplementary Figure 4C**). The final set of feature values are then normalized and PCA-transformed to reduce correlation within training values for classification. The PCA occurs independently between folds. By default, the number of components retained from the PCA-transformation is the smallest set of PCA components such that at least 75% of the variance in the feature set is explained. The PCA-transformed training data is then used to train a suite of models for classification (**Supplementary Figure 4D**). In this way, the models are generally formed independently from each other except for the aggregated feature selection, for the sake of interpretability.

#### Classification

A suite of different, standard classification models built into MATLAB were trained, including architectures such as decision trees, logistic regression classifiers, Support Vector Machines (SVM), and k-nearest neighbor (KNN) classifiers on the normalized, PCA-transformed, selected O-SNAP features as described under **Feature selection and principal component analysis** under **Methods**. Five replicates of the same model are trained. To evaluate the performance of the models, the 20% reserved data is independently normalized, followed by filtering for the selected features and undergoes a PCA-transformation applied to the respective training data. The trained models then predict on the test data, and the ground truth labels are compared to the predicted labels.

For each individual trained model, the overall accuracy is simply obtained from the proportion of samples correctly classified compared to the total test sample count. For each model architecture and within each fold, the accuracies of the five model replicates are averaged, also providing a standard deviation on the model training within the fold. These mean accuracies are then averaged again across the five folds to obtain a statistics on the overall accuracy of the model architecture for the full dataset. The overall accuracies across different models were compared, and the model with the highest mean accuracy was considered the best model for a given O-SNAP classification and was assessed further to better understand its performance.

Receiver Operator Characteristic (ROC) curves were one approach to evaluate the classification performance. ROC curves are obtained by plotting the true positive rate (TPR) against the false positive rate at different thresholds. These curves depict how two rates change depending on a threshold value that alters the sensitivity of the classifier to a specific phenotype at the cost of correctly identifying the others. The results shown represent the distribution of the ROC curves across the five folds for a given model architecture.

Confusion matrices offered another means to understand classifier performance. They display the number or proportion of correct identifications along the diagonal and misclassifications in the off-diagonal positions of the matrix. To derive these summary plots, confusion matrices are first generated for each of the five folds and the raw sample counts are used to populate the matrix, which are averaged within each fold over the five replicates of the same model type. The confusion matrices are then row-normalized so that the confusion matrix cells represent the proportion of true labels that were correctly or incorrectly classified. To obtain statistics of the classification results across the five folds, the weighted average and standard deviation were calculated for each normalized row, where the weights are the total number of test samples in the cell state the given row represents. This approach accounts for differences in the number of test sample counts of a given phenotype that may have been present for separate samples, as only the total test sample size was held constant but not the proportion of test samples from each class. The final values shown are these weighted means and standard deviations.

O-SNAP also performs a similar classification pipeline, but in place of the aggregated feature selection, maintains the individual fold-wise feature selections so that the full process of classification is fully independent between folds. The results of these are not shown but are available in the analysis. Another classification pipeline is also available, where normalization and feature selection occur on the entire data set and is followed by the five-fold cross-validation approach. However, this approach is prone to data leakage, where the trained classification models are indirectly informed about the test data set since the feature selection is performed on the entire data set. Hence, to achieve a robust approach while maintaining interpretability, O-SNAP implements the aggregated feature selection approach instead.

To determine whether O-SNAP was identifying true differences, as opposed to random variance, in chromatin organization between phenotypes, the classification performances for the TSA and reprogramming O-SNAP results were compared to that of models trained on the same dataset with shuffled phenotype labels (**Supplementary Figure 5D** and **Figure 4C**). In the shuffled dataset, only the class labels are permuted such that correlations between features in a sample are still preserved. The shuffling only occurs once prior to all subsequent O-SNAP analysis. Thus, normalization, feature selection, dimensionality reduction, and classification are applied to the shuffled data in the same fashion as the ground truth data.

## Acknowledgements

M.L. acknowledges funding from NIH Common Fund 4DNucleome Consortium under the grant number U01DA052715, NIH/NIAMS under the grant numbers R01 AR079224 and P50 AR080581, NIH/NIA under the grant number R01 AG082437, NIH/NIGMS under the grant number R35 GM152111 and NSF Center for Engineering Mechanobiology (CEMB) under the grant number CMMI-1548571. H.H.K acknowledges funding from the Structural Biology and Molecular Biophysics T32 training grant (5-T32-GM-132039-05) from the NIGMS of the NIH. We thank Siewert Hugelier (UPenn) for helpful reading and feedback on the manuscript.

## Author contributions statement

H.H.K. and M.L. conceived and designed the research study. H.H.K designed and implemented the O-SNAP framework. H.H.K., J.A.M.-S., F.R.P., E.Y.Z., and Z.G performed experimental work and provided data. H.H.K. performed O-SNAP analysis. H.H.K. and M.L. wrote the manuscript with input from all other authors.

## Data and Code Availability

Previously published data used to validate O-SNAP consist of the human tenocyte STORM dataset, publicly available at https://doi.org/10.1038/s41551-022-00910-5 (ref.^32^) and the fibroblast reprogramming STORM dataset, which is publicly available at http://doi.org/10.5061/dryad.931zcrjqp (ref.^31^). Datasets collected for this work are publicly available via figshare at https://doi.org/10.6084/m9.figshare.29533940. Example data for O-SNAP are included with the source code via https://github.com/LakGroup/O-SNAP (ref.^96^).

