## Supplementary Information for "O-SNAP: A comprehensive pipeline for spatial profiling of chromatin architecture"

### Supplementary Figures

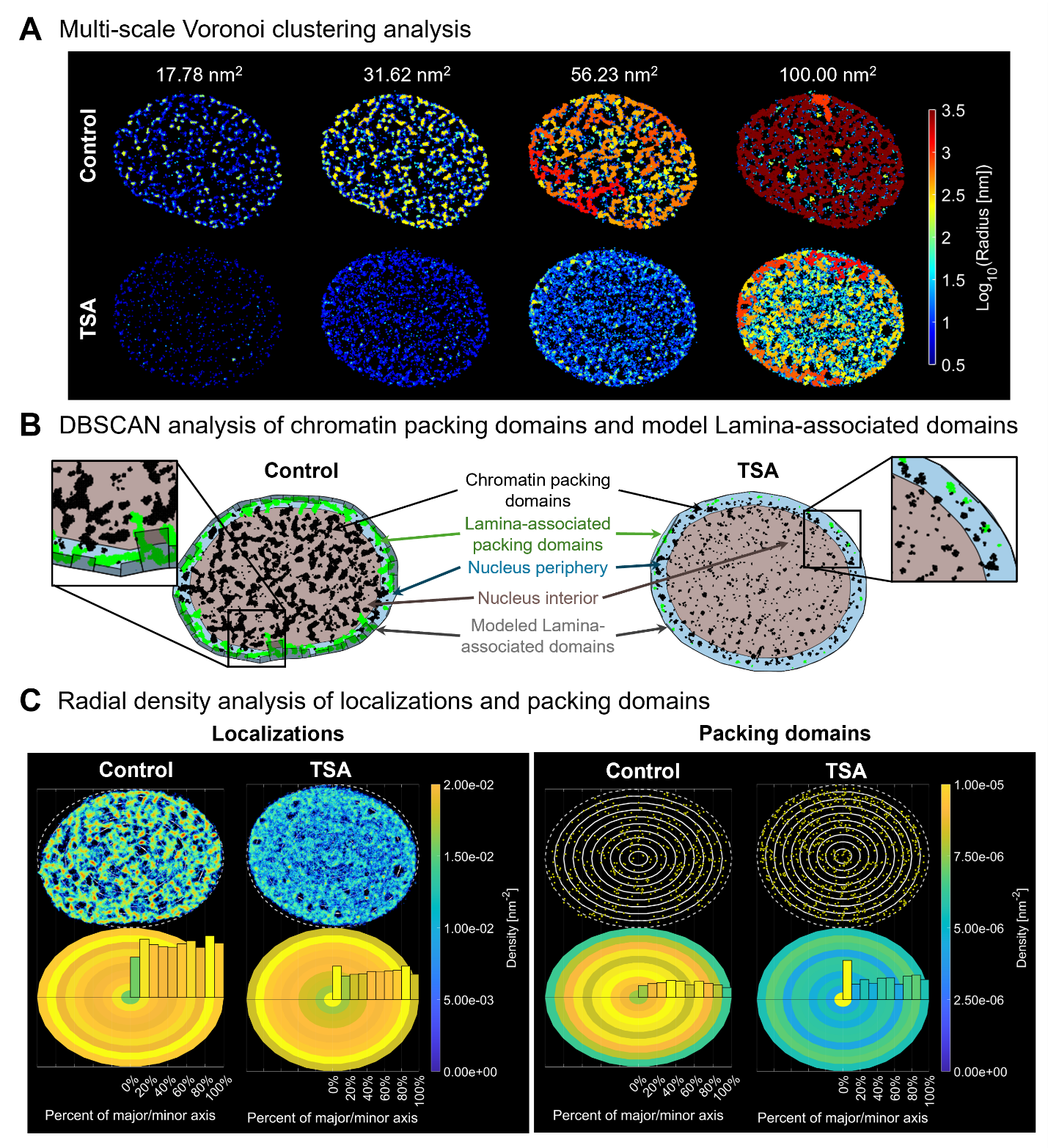

**Supplementary Figure 1. Multi-scale Voronoi clustering, DBSCAN, and partitioning of the nucleus**. Examples shown here are derived from control and TSA-treated human fibroblasts labeled for H2B (see **Figure 2**). (A) Voronoi clustering performed at multiple length scales (17.78 nm^2^, 32.61 nm^2^, 56.23 nm^2^, and 100.0 nm^2^; see **Voronoi clustering analysis** in **Methods**), where clusters are color-coded based on their size, demonstrating the increase in cluster size as the threshold is progressively relaxed. (B) Chromatin packing domains, identified by performing DBSCAN on localizations with Voronoi densities within the top 70^th^ percentile, shown in black and green. A subset of these domains at the periphery are identified as putative lamina-associated domains (green) if the cluster falls within 5% of the nucleus radius from the nucleus boundary. These domains inform the distribution of 50 quadrilaterals, the modeled lamina-associated domains (gray boxes). The nucleus area is separated into two regions, where the nucleus interior is the inner 85% of the total nucleus area (brown) whereas the periphery represents the outer 15% (blue; see **DBSCAN clustering analysis** in **Methods**). (C) Radial analysis models the nucleus area as a series of concentric rings where the number of localizations (left) or packing domains (right) per unit area is calculated for each ring (see **Radial density analysis** in **Methods**).

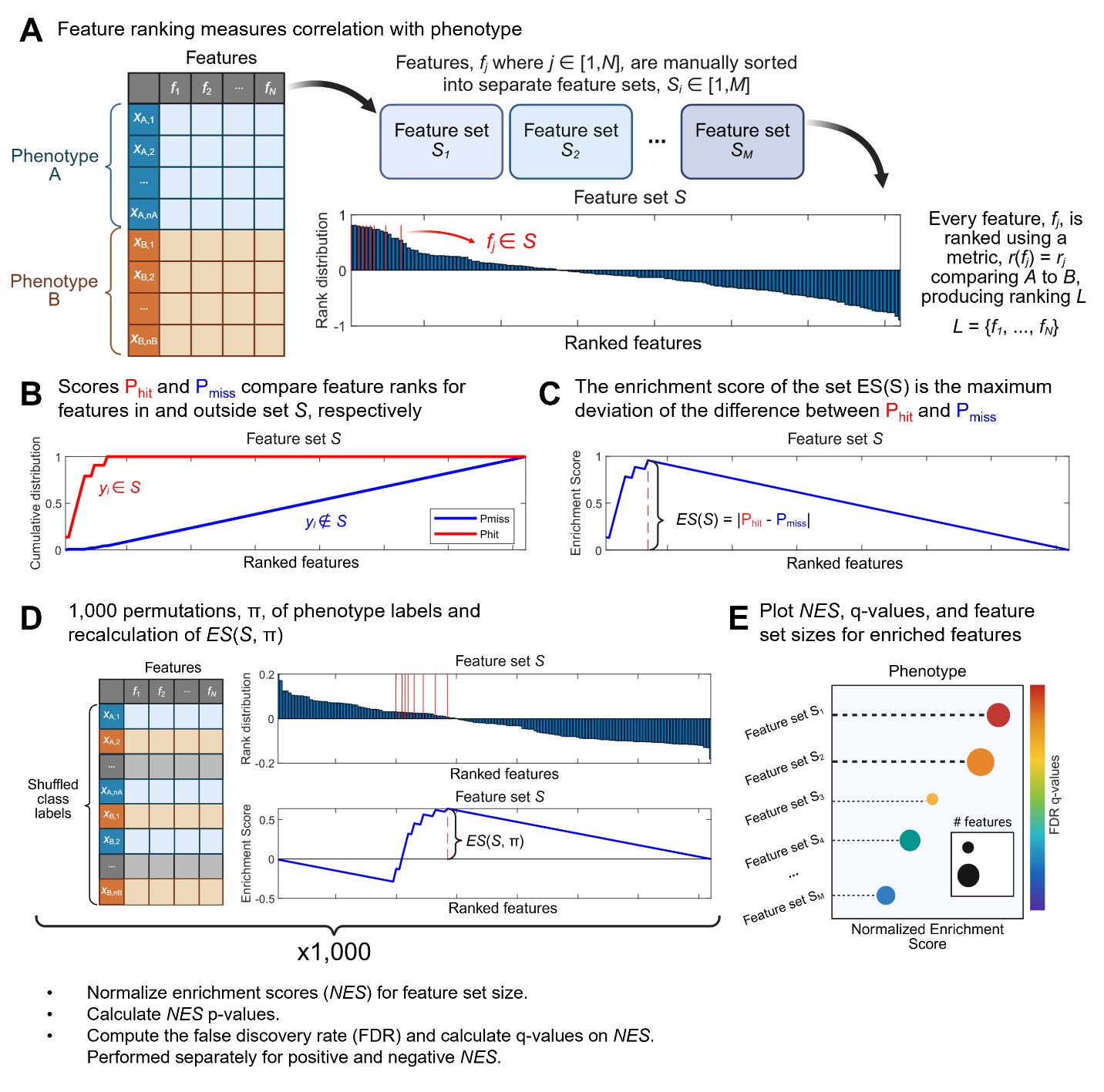

**Supplementary Figure 2. Feature set enrichment analysis workflow.** (A) Feature sets, $S$, are manually defined groups of features describing a similar, overall trend, such as “larger nucleus.” Following normalization to z-scores, the feature values, $f$, are sorted using one of 8 ranking metrics, $r(f)$, to create a ranking, $L$. (B) For each feature set, $S$, traversing the feature ranking, $L$, generates cumulative scores for features within ($P_{\text{hit}}$) and outside ($P_{\text{miss}}$) the feature set. (C) The maximum deviation of $P_{\text{hit}} - P_{\text{miss}}$ from zero along the ranking traversal is taken as the enrichment score $ES(S)$ of feature set, *S*. To calculate statistical significance, the data must be compared to a null distribution of the enrichment score. This distribution is formed from 1,000 permutations, π, of the phenotype labels of the data and recalculating a new enrichment score, $ES(S, \pi)$. The distribution of $ES(S, \pi)$ is used subsequently to normalize the enrichment scores, $NES(S)$, and perform significance testing. These are performed separately for positive and negative values of $ES(S)$. (E) A final plot is generated to compare the normalized enrichment scores, q-values, and the size between feature sets. (See **Feature set enrichment analysis** under **Methods**)

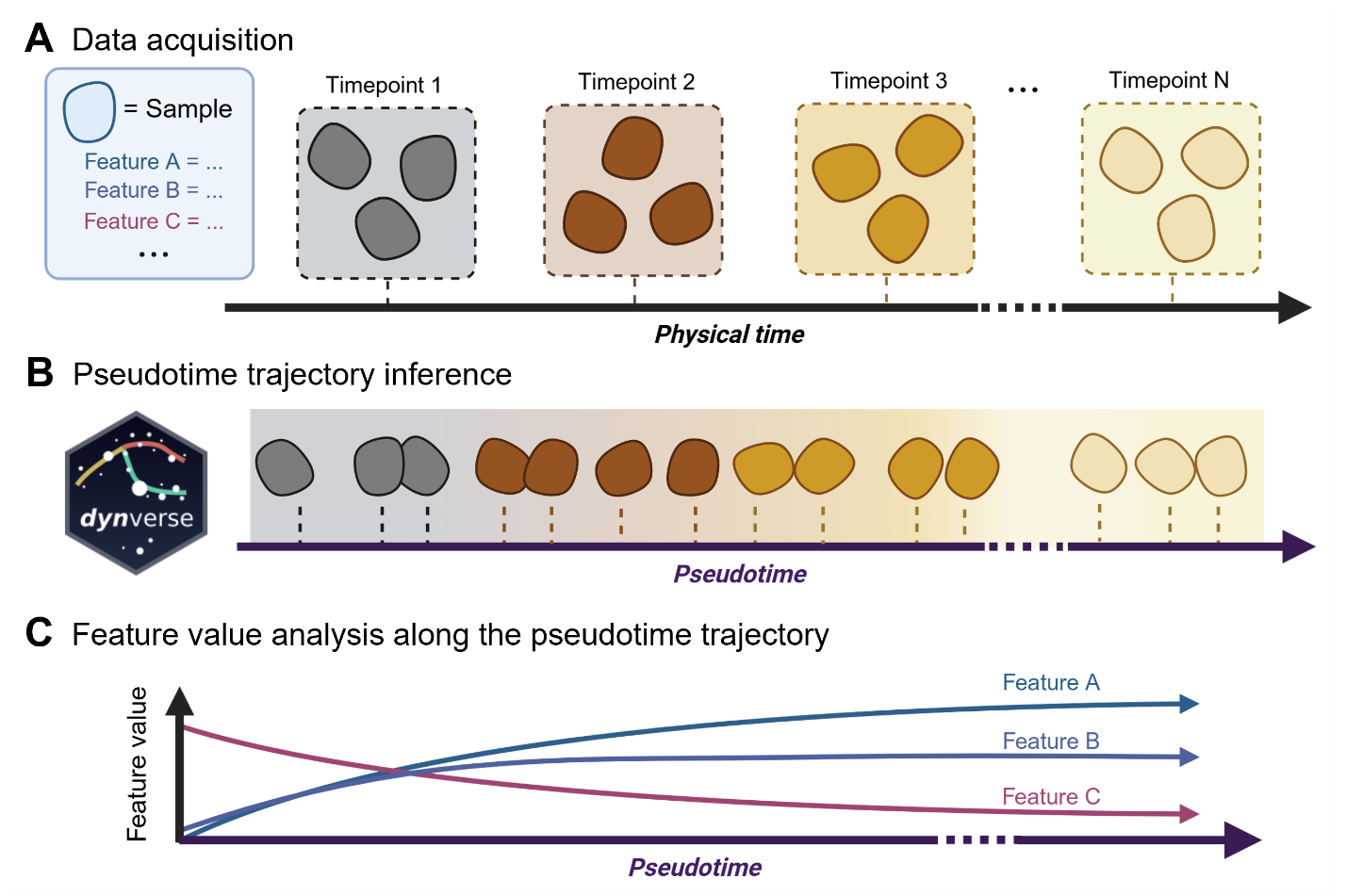

**Supplementary Figure 3. Pseudotimeline trajectory analysis workflow.** (A) Timepoints for experimental data are acquired at discrete intervals. (B) A trajectory inference method, implemented here using the *dynverse* package, estimates the position of each sample along a continuous timeline based on O-SNAP feature information. The details of how samples are assigned pseudotime values depends on the trajectory method. (C) Because samples are organized in a continuous manner, their respective feature values can be observed relative to pseudotime. (See **Pseudotime analysis** under **Methods**)

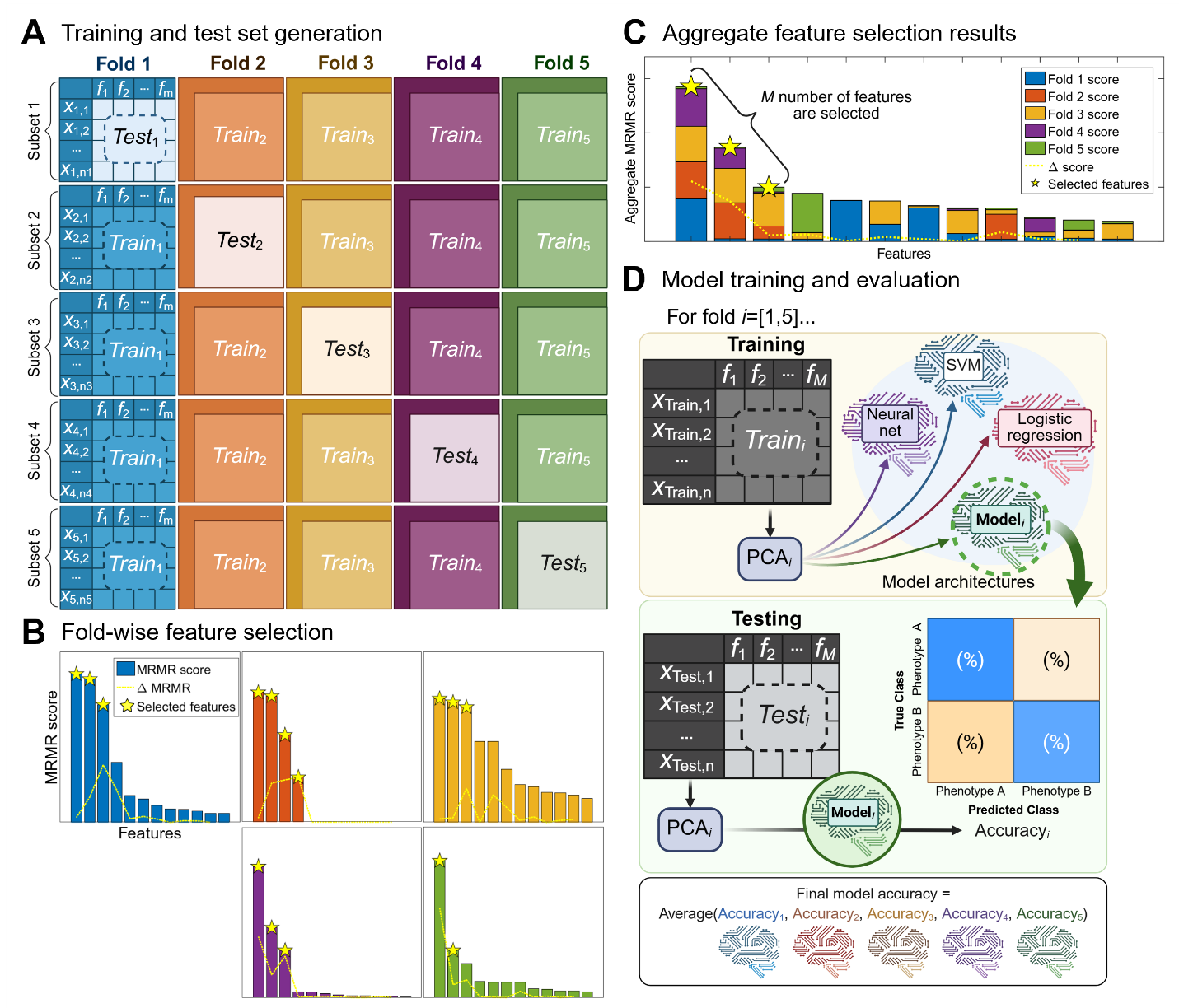

**Supplementary Figure 4. 5-fold cross validation pool generation, feature selection, and classification workflow.** To assess model performance, O-SNAP implements a 5-fold cross validation. (A) The entire dataset is pooled, and samples are shuffled. A different fifth of the samples is reserved to test the final model for each fold. The remaining four-fifths of the data is used for feature selection. (B) Based on the training data within each fold, the MRMR algorithm identifies the features with the greatest discrimination potential between the phenotypes. (C) The feature selection results from every fold are summed together to obtain the aggregate MRMR score. Identifying the knee point of the scores or the maximum change in consecutive features (whichever occurs first) produces the final feature subset that the models train on. (D) In each fold, the training data is filtered for the selected features and PCA-transformed for further dimensionality reduction. A suite of different model architectures trains on the PCA-transformed data. For each fold, the model is validated against the reserved test data, which undergoes the same filtering and PCA-transformation as the training data. The model with the highest average accuracy across the folds is used as the final model. (See **Feature selection and principal component analysis** and **Classification** under **Methods**)

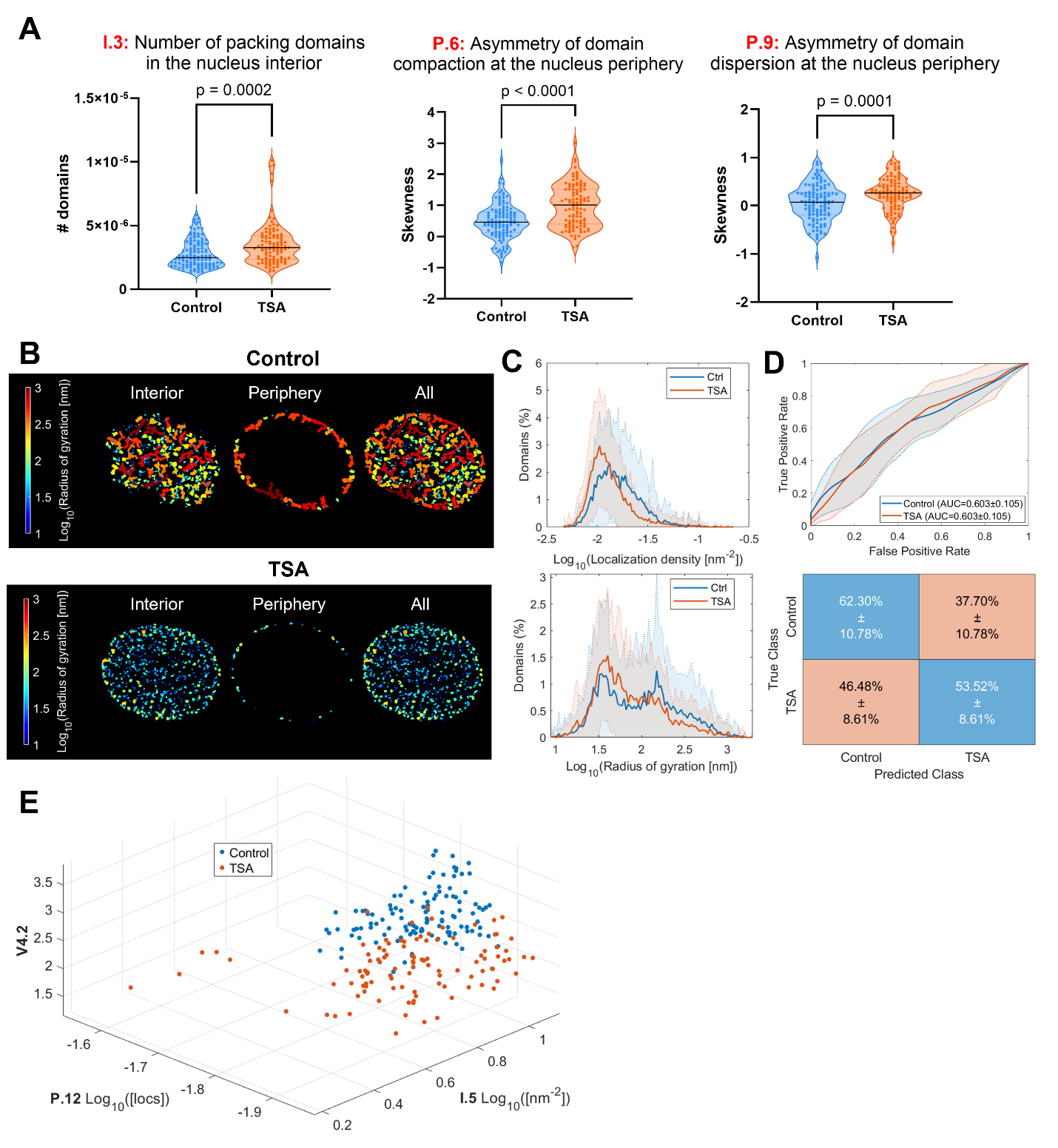

**Supplementary Figure 5. Additional results related to chromatin changes in response to histone deacetylase inhibition.** (A) Violin plots corresponding to the features identified as significantly changing in the volcano analysis: the abundance of packing domains at the nucleus interior (left), the asymmetry in the domain compaction (i.e. localization density) at the nucleus periphery (center), and the asymmetry in the dispersion (i.e. radius of gyration) of localizations in domains at the nucleus periphery (right). A two-way Welch’s t-test was used to calculate p-values and statistical significance. (B) Representative nuclei from **Figure 2A**, where packing domains are color-coded based on their respective radius of gyration for control (top) or TSA-treated cells (bottom). (C) Distributions of the packing domain properties from features identified from the volcano analysis comparing TSA-treated to control cells. The compactness of periphery domains from Feature P.6 (top) and the radius of gyration of the periphery domains from Feature P.9 (bottom). The solid line shows the percentage of packing domains that have a given value for a given bin, averaged over all nuclei of either the control or TSA treatment condition, and the shaded region represents ±1 standard deviation interval from the mean for each bin. (D) Classification on the TSA-treated system with shuffled class labels to assess the reliability of the ground-truth results. The ROC curve (top) and confusion matrix (bottom) of an ensemble bagged trees model trained to discriminate control from TSA-treated fibroblasts across five training/test batches on the feature data with shuffled cell labels with an overall validation accuracy of 57.99 ± 9.04%. (E) Scatter plot of the selected features with the highest aggregate MRMR rankings (I.5, P.12, and V4.2; features identified from **Figure 2E**). This data demonstrates that the features together demonstrate a distinct distribution of the control nuclei (blue) from the TSA-treated cells (orange).

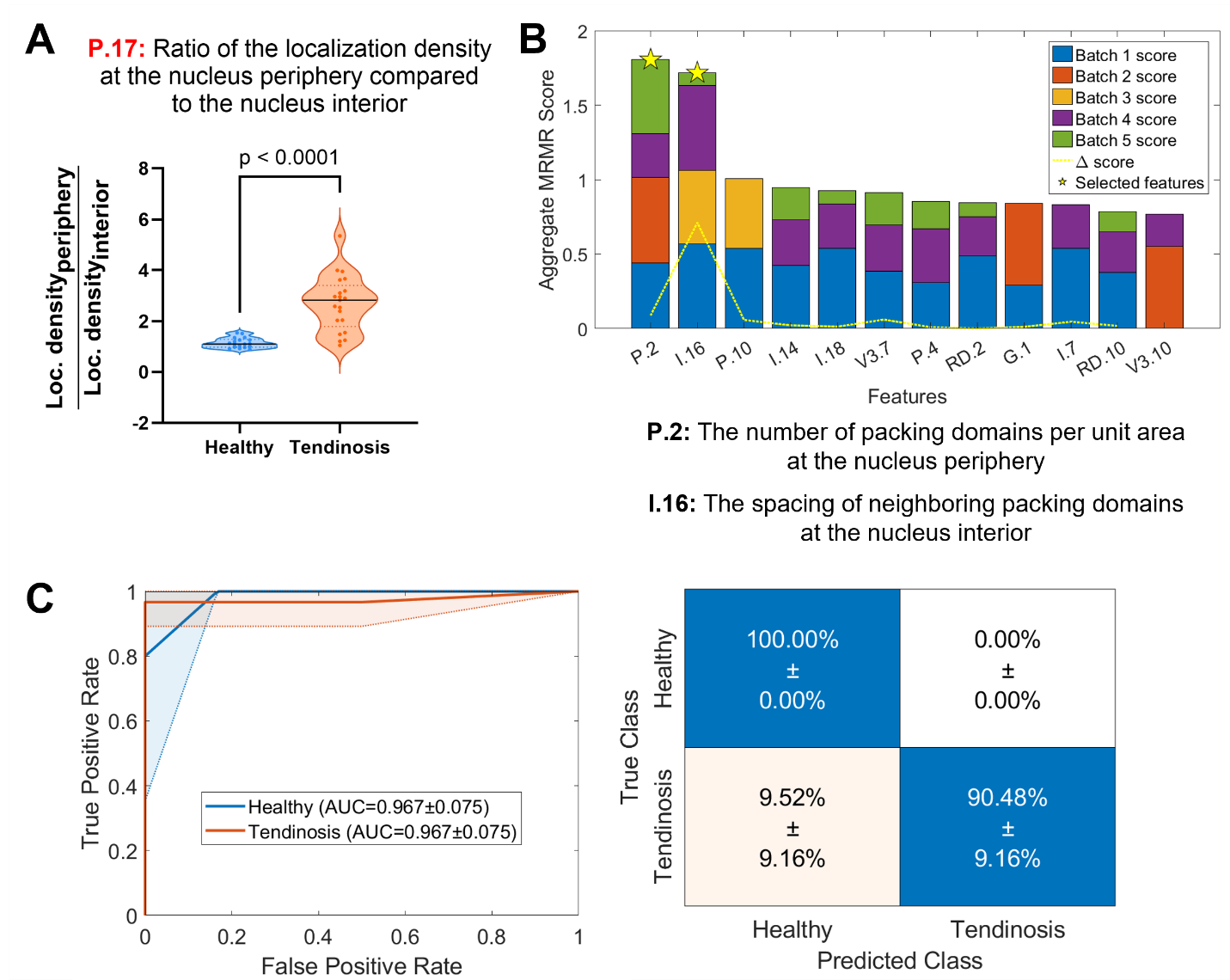

**Supplementary Figure 6. Additional results related to global chromatin organization of tenocytes derived from healthy and diseased (tendinosis) donors.** (A) The distribution of the ratio of the localization density of H2B signal to the nucleus periphery relative to the nucleus interior replicates the increase in relative density at the periphery reported in Heo et al, Nature BME, 2023*.* A two-way Welch’s t-test was used to calculate p-values and statistical significance. (B) The aggregated MRMR feature selection shows that Features P.2 (number of packing domains per unit area at the nucleus periphery) and I.16 (the spacing of neighboring packing domains at the nucleus interior) possess the highest potential to separate out healthy from tendinosis cells. (C) Classification results of a quadratic discriminant classifier trained on the PCA transformation of features selected from **Supplementary Figure 6B**, to discriminate control from healthy and tendinosis with an overall model accuracy is 95.00 ± 6.85%. The ROC curve (left) and confusion matrix (right) demonstrate the performance with respect to each condition across the five folds.

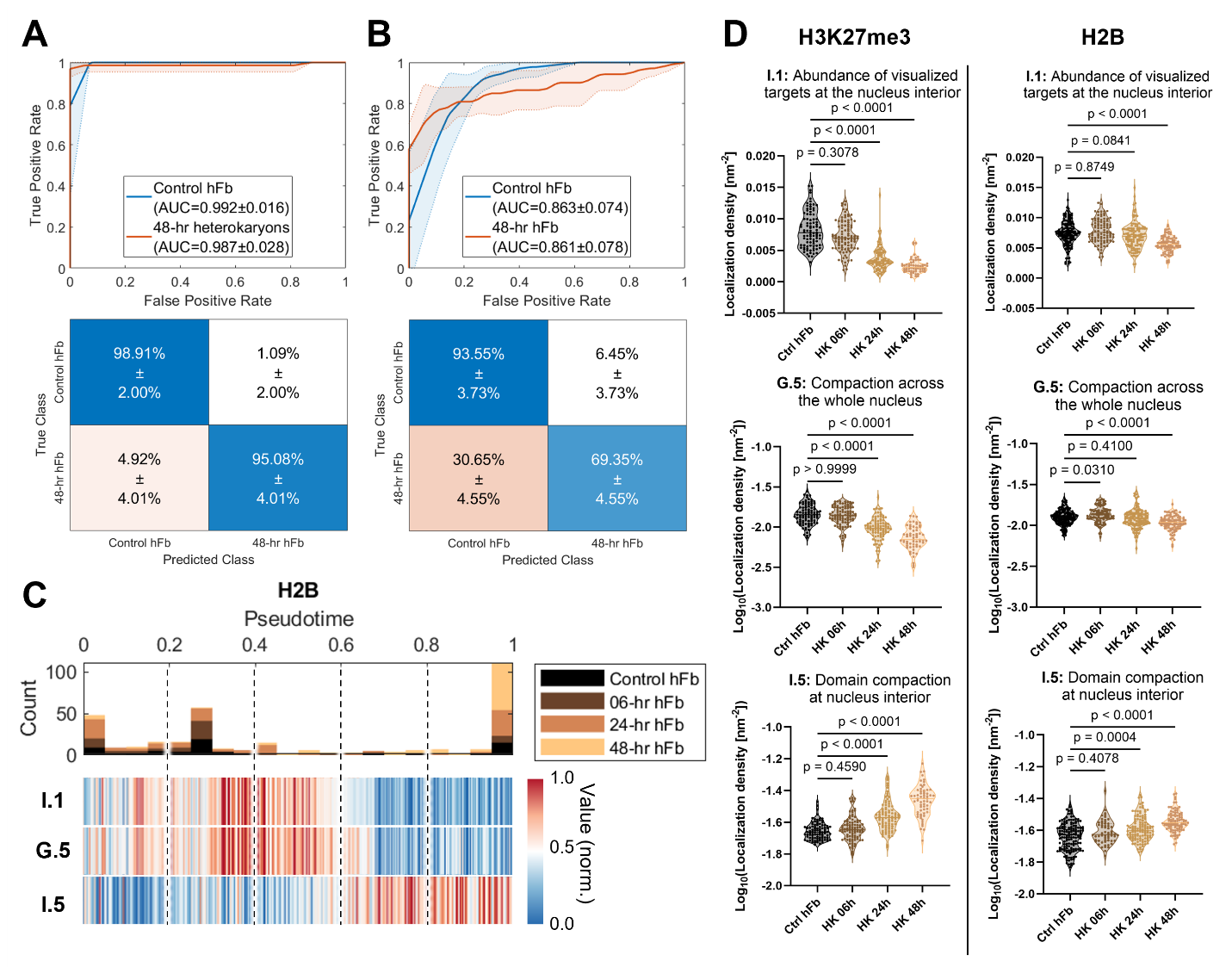

**Supplementary Figure 7. Additional results regarding changes to global chromatin and epigenetic markers in early heterokaryon reprogramming.** (A) Classification results of a linear SVM classifier trained to discriminate control fibroblast nuclei from fibroblast nuclei after 48-hr of cell fusion using O-SNAP features from H3K27me3 SMLM data, where the overall model accuracy is 97.40 ± 1.46%. The ROC curve (top) and confusion matrix (bottom) show the performance for each cell state across the five folds. (B) Classification results of a gaussian SVM classifier trained to discriminate control fibroblast nuclei from the fibroblast nuclei after 48-hr of cell fusion using O-SNAP features from H3K9ac SMLM data, where the overall model accuracy is 83.87 ± 3.95%. The ROC curve (top) and confusion matrix (bottom) show the performance for each cell state across the five folds. (C) Histogram of assigned pseudotime calculated using the Slingshot trajectory inference method on O-SNAP features generated from the H2B heterokaryon SMLM data. Bottom: A heatmap of H2B O-SNAP features changing with pseudotime. The features displayed are the same as those in **Figure 4H**, but do not demonstrate as smooth a transition with respect to pseudotime in contrast to the H3K27me3 data. (D) The spatial features identified in pseudotime analysis throughout heterokaryon reprogramming (**Figure 4H**) for H3K27me3 data (left) and H2B data (right): the nuclear interior localization density (I.1; top row), the global localization compaction (G.5; middle row), and the density of packing domains within the nuclear interior (I.5; bottom row). In the H3K27me3 data, both I.1 and G.5 showed a significant decrease and I.5 exhibited a significant increase with reprogramming by the 24-hour timepoint. In the H2B data, I.1 and G.5 show significant decrease by the 48-hr timepoint whereas I.5 shows increased density by the 24-hr timepoint. One-way ANOVA and Dunnett multiple comparisons test were used to calculate p-values and statistical significance.

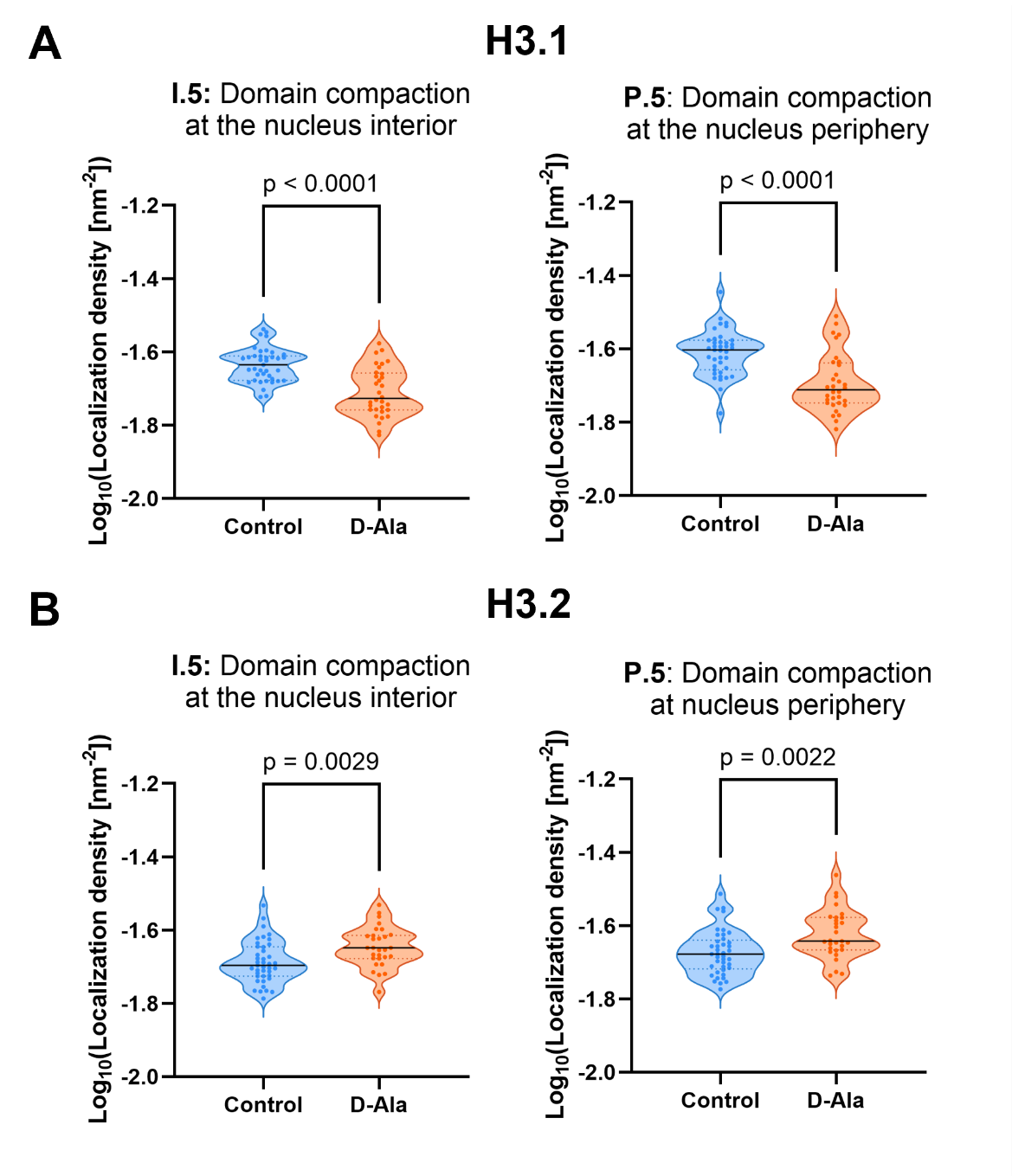

**Supplementary Figure 8. Changes to domain compaction in MCF10A/NLS-DAO overexpressing histone variant H3.1 or H3.2 upon treatment with D-Ala.** (A) Violin plots of mean domain compaction for SMLM data of H3.1 overexpression. The average localization density per unit area across packing domains at the nucleus interior (left) and the nucleus periphery (right). The D-Ala group shows a decrease in domain compaction for both regions of the nucleus. (B) Violin plots of mean domain compaction for SMLM data of H3.2 overexpression. The average localization density per unit area across packing domains at the nucleus interior (left) and the nucleus periphery (right). (A, B) A two-way Welch’s t-test was used to calculate p-values and statistical significance.

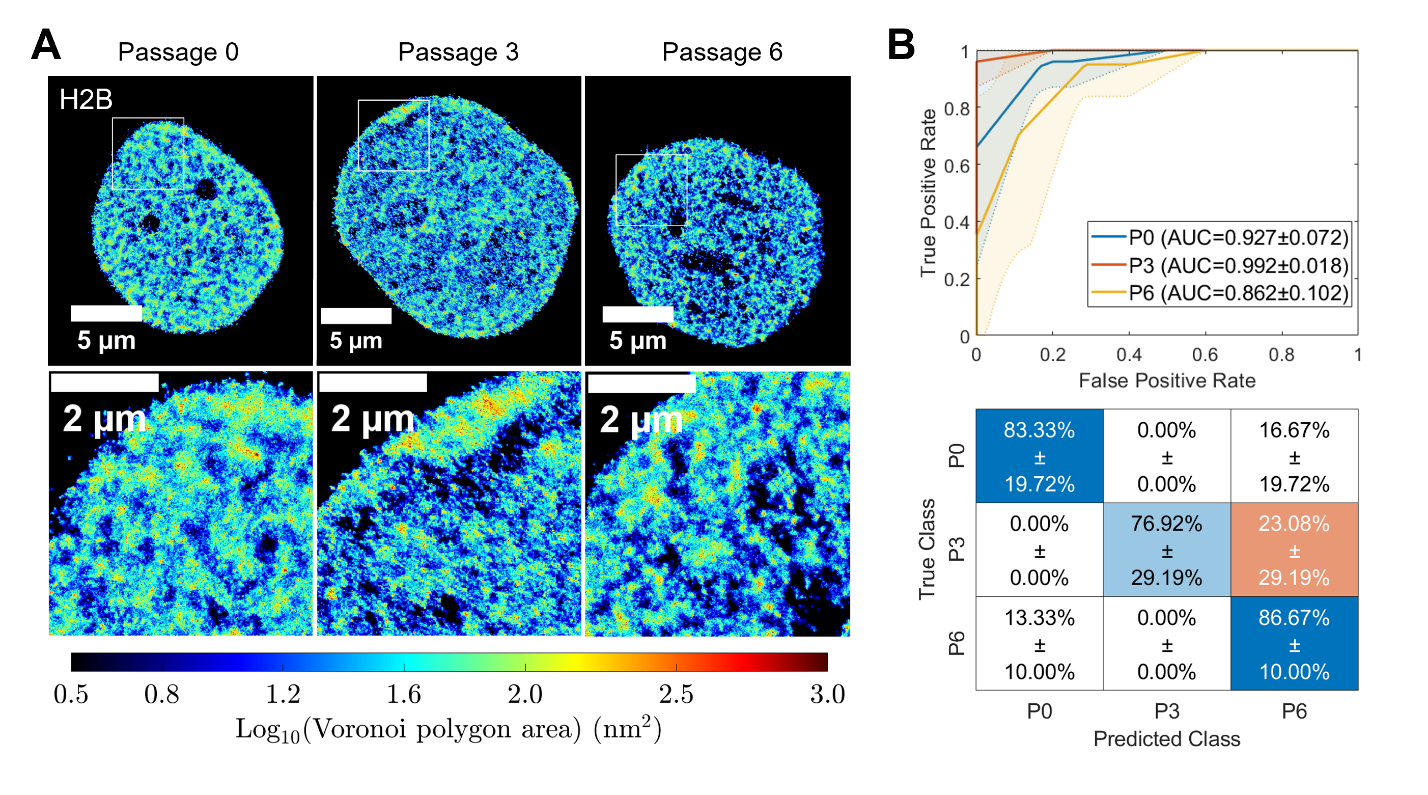

**Supplementary Figure 9. Representative images and classification for Donor B in the human chondrocyte system and distributions of features identified from volcano plot analysis.** (A) Representative Voronoi density map renderings of H2B STORM images of human chondrocytes at P0, P3, and P6 from Donor B (N=18 nuclei for P0, N=13 for P3, and N=15 for P6). The color code indicates local chromatin compaction from low density (blue) to high density (red). (B) Classification results of a Naïve Bayes gaussian classifier trained to discriminate P0, P3, and P6 nuclei of human chondrocytes derived from Donor B, demonstrating an overall accuracy of 82.89 ± 8.66%. The ROC (top) and confusion matrix (bottom) show the performance for each state across the five folds, which appears comparable to that of Donor A.

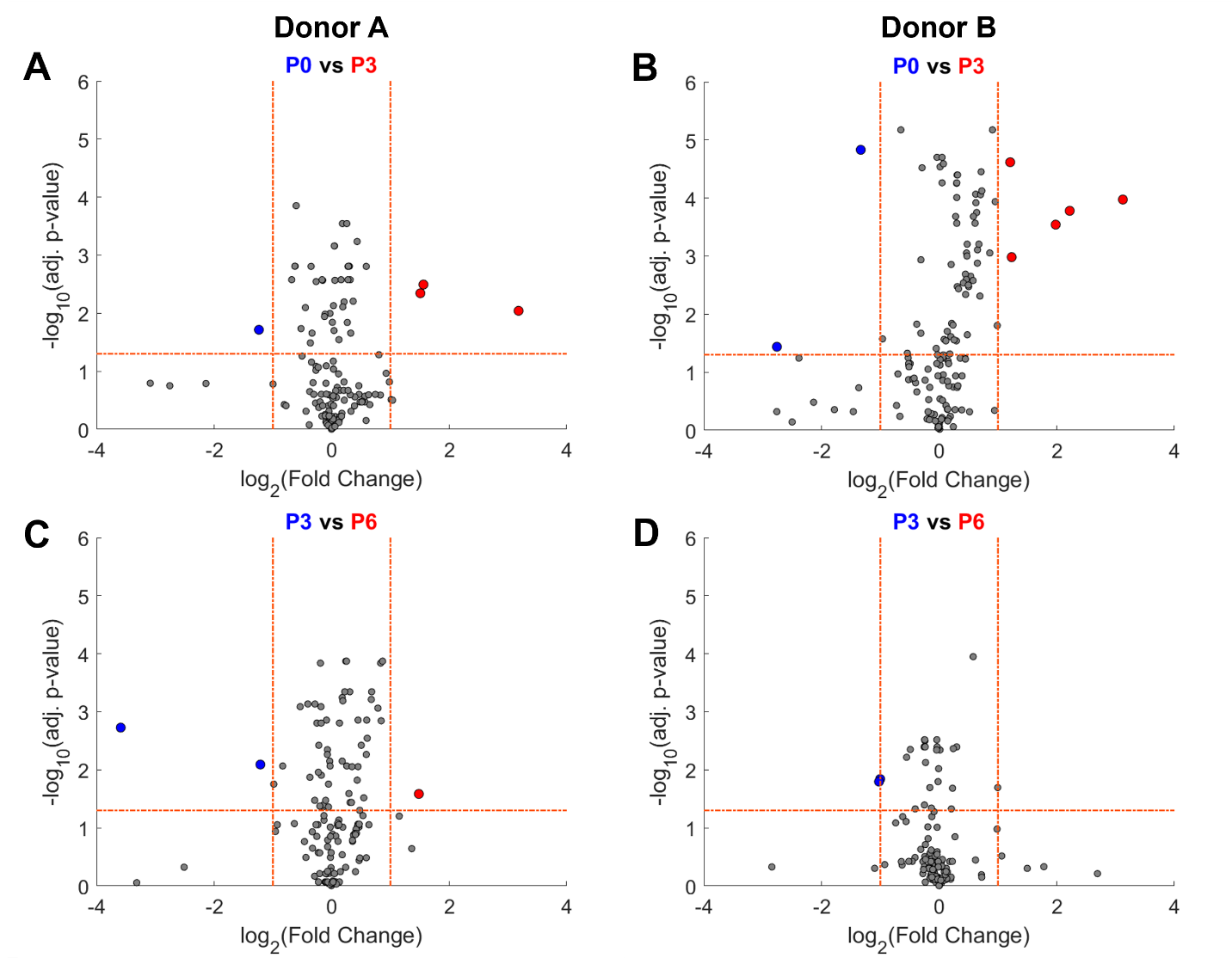

**Supplementary Figure 10. Volcano plots of changes from P0 to P3 and P3 to P6 for each donor.** The volcano plots demonstrate that features are changing significantly, both between the P0 and P3 (A, B) as well as the P3 to P6 (C, D) timepoints. In contrast to the P0 to P6 transition, there are no features in common between Donor A and B for these comparisons.

### Supplementary Tables

**Supplementary Table 1.** Recurring terms used in definitions of O-SNAP features.

| Term | Definition |
| --- | --- |
| Chromatin packing domain,  Nucleosome clutch,  DBSCAN domain | A single cluster of localizations. Generated by performing DBSCAN to segment the population of localizations within the top 70^th^ percentile of Voronoi density, which corresponds to the most compact compartment. |
| Nucleus interior | The innermost 85% (relaxed) or 95% (strict) region of the nucleus area |
| Nucleus periphery | The outermost 15% (relaxed) or 5% (strict) region of the nucleus area |
| Modeled Lamina-Associated Domain (LAD) | A quadrilateral formed based on the distribution of target clusters at a segment of the nucleus boundary. 50 total model LADs are generated to span the nucleus boundary. |
| Segment length | Length of modeled LAD tangent to the nucleus boundary |
| Segment thickness | Length of modeled LAD perpendicular to boundary |
| Spacing | The average nearest-neighbor distance between a DBSCAN domain and its nearest *K* neighbors (default *K* = 5), measured from centroid to centroid |
| Voronoi domain | A single cluster of localizations. Generated by performing Voronoi clustering to segment the population of localizations with a respective Voronoi area below a defined threshold. The clustering is performed by identifying regions of contiguous Voronoi cells that meet this criterion. |
| Voronoi density | The inverse of the Voronoi cell area for a given localization. |
| Reduced Voronoi density | The inverse of the Voronoi cell area for a given localization, normalized by the total number of localizations divided by the nucleus area |

**Supplementary Table 2.** Abbreviations of the categories the features broadly fall under, which is used to create the shorthand system used in the text.

| Shorthand | Category |
| --- | --- |
| M | Nucleus morphology |
| G | Global localization density |
| I | Nucleus interior |
| P | Nucleus periphery |
| L | Modeled LAD |
| R | Radial localization density |
| RD | Radial packing domain density |
| V(#) | Voronoi domain (indicates the upper area threshold in nm) |

**Supplementary Table 3.** Full description of O-SNAP features with the shorthand, feature variable name as it appears in the MATLAB scripts, putative biological interpretation of the feature, the precise definition of the feature, details on how the feature is calculated when relevant, the default units of the feature (note some features undergo a Log-transformation, but the original feature unit is included), and finally the set the feature belongs for feature set enrichment analysis (FSEA).

| # | Short-hand | Feature  *Label as it appears in the code and output* | Putative biological interpretation | Definition | Details | Units | Feature set |
| --- | --- | --- | --- | --- | --- | --- | --- |
| 1 | M.1 | aspect_ratio | Nucleus elongation | Ratio of the major to minor axis | Aspect Ratio $= a/b$ | *Dimensionless* | Rounder nucleus |
| 2 | M.2 | bending_energy | Nucleus boundary irregularity | Metric to quantify second-order boundary deformation | Bending Energy $=\frac{1}{2}\cdot\sum\Delta{\Delta x}^{2}$ | [nm^2^] | Rounder nucleus |
| 3 | M.3 | bending_energy_norm_area | Nucleus boundary irregularity | Bending energy normalized by the area of the nucleus | Bending Energy Normalized by Area $=\text{Bending Energy}\div A$ | *Dimensionless* | Rounder nucleus |
| 4 | M.4 | circularity | Nucleus roundness | Degree of similarity of nucleus to a true circle | Circularity $=\frac{4\pi A}{P^{2}}$ | *Dimensionless* | Rounder nucleus |
| 5 | M.5 | convex_area | Nucleus size | Area of a convex hull containing all the localizations | Convex Area $=A_{C}$ | [nm^2^] | Larger nucleus |
| 6 | M.6 | convex_circularity | Nucleus roundness | Circularity of the convex hull containing all localizations | Convex Circularity $=\frac{4\pi A_{C}}{P_{C}^{2}}$ | *Dimensionless* | Rounder nucleus |
| 7 | M.7 | convex_perimeter | Nucleus size | Perimeter of a convex hull containing all the localizations | Convex Perimeter $=P_{C}$ | [nm] | Larger nucleus |
| 8 | M.8 | convexity | Nucleus roundness | The ratio of the convex perimeter to the true perimeter of the nucleus | Convexity $=P_{C}/P$ | *Dimensionless* | Rounder nucleus |
| 9 | M.9 | curl | Nucleus boundary roundness | Ratio of the major axis to fiber length measuring degree that the object curls upon itself | Curl $=\frac{a}{\text{Fiber Length}}$ | *Dimensionless* | Rounder nucleus |
| 10 | M.10 | eigenentropy | Nucleus elongation | Entropy of the eigenvalues derived from a PCA transformation on the localizations | Eigen-Entropy $=-\frac{\lambda_{1}}{\lambda_{1}+\lambda_{2}}\cdot\ln\left( \frac{\lambda_{1}}{\lambda_{1}+\lambda_{2}} \right)-\frac{\lambda_{2}}{\lambda_{1}+\lambda_{2}}\cdot\ln\left( \frac{\lambda_{2}}{\lambda_{1}+\lambda_{2}} \right)$ | *Dimensionless* | Rounder nucleus |
| 11 | M.11 | eigenvalues_ratio | Nucleus elongation | The ratio of eigenvalues derived from a PCA transformation on the localizations | Eigenvalue Ratio $=\lambda_{1}/\lambda_{2}$ | *Dimensionless* | Rounder nucleus |
| 12 | M.12 | elastic_energy | Nucleus boundary irregularity | Metric to quantify first-order boundary deformation | Elastic Energy $=\frac{1}{2}\cdot\sum\Delta x^{2}$ | [nm^2^] | Rounder nucleus |
| 13 | M.13 | elastic_energy_norm_area | Nucleus boundary irregularity | Elastic energy normalized by the area of the nucleus | Elastic Energy Normalized by Area $=\text{Elastic Energy}\div A$ | *Dimensionless* | Rounder nucleus |
| 14 | M.14 | fiber_length | Nucleus size and elongation | Metric approximating the longest, continuous length within the shape | Fiber Length $=P+\frac{\sqrt{\left\vert P^{2}-16*A \right\vert}}{4}$ | [nm] | Larger nucleus |
| 15 | M.15 | fiber_width | Nucleus size and elongation | Ratio of the nucleus area to fiber length | Fiber Width $= \frac{A}{\text{Fiber Length}}$ | [nm] | Larger nucleus |
| 16 | M.16 | form_ratio | Nucleus elongation | Ratio of the nucleus area to a square with side length equal to the major axis | Form Ratio $= A/a^{2}$ | *Dimensionless* | Rounder nucleus |
| 17 | M.17 | gyration_radius | Target distribution and nucleus shape | Radius of gyration of the nucleus | Calculated using the second moment of localizations  Radius of Gyration ${= R}_{g}= \sqrt{\frac{\sum_{i=1}^{N} r_{i}^{2}}{N}}$ | [nm] | Rounder nucleus |
| 18 | M.18 | log_border_curvature_mean | Nucleus roundness | Average border curvature of the nucleus boundary | Border curvature is a distribution calculated from multiple segments along the nucleus boundary. | [nm^-1^] | Rounder nucleus |
| 19 | M.19 | log_border_curvature_skewness | Nucleus boundary irregularity | Skewness of the border curvature distribution of the nucleus boundary | Border curvature is a distribution calculated from multiple segments along the nucleus boundary. | *Dimensionless* | Rounder nucleus |
| 20 | M.20 | log_border_curvature_std | Nucleus boundary irregularity | Standard deviation of the border curvature distribution of the nucleus boundary | Border curvature is a distribution calculated from multiple segments along the nucleus boundary. The standard deviation describes the variance of the curvature distribution. | [nm^-1^] | Rounder nucleus |
| 21 | M.21 | major_axis | Nucleus size | Longest diameter across nucleus | Major Axis $=a$ | [nm] | Larger nucleus |
| 22 | M.22 | major_axis_norm_area | Nucleus elongation | Length of the major axis normalized by the area of the nucleus | Major Axis Normalized by Area $=a/A$ | [nm^-1^] | Rounder nucleus |
| 23 | M.23 | minor_axis | Nucleus size | Diameter perpendicular to major axis | Minor Axis $=b$ | [nm] | Larger nucleus |
| 24 | M.24 | minor_axis_norm_area | Nucleus elongation | Length of the minor axis normalized by the area of the nucleus | Minor Axis Normalized by Area $=b/A$ | [nm^-1^] | Rounder nucleus |
| 25 | M.25 | nucleus_radius | Nucleus size | Radius of a circle whose area is equal to that of the nucleus | Radius $=R=\sqrt{A/\pi}$ | [nm] | Larger nucleus |
| 26 | M.26 | perimeter | Nucleus size | Perimeter of the nucleus | Perimeter $=P$ | [nm] | Larger nucleus |
| 27 | M.27 | rectangularity | Nucleus roundness | Ratio of the nucleus area to a rectangle with side lengths equal to the major and minor axis | Rectangularity $= \frac{A}{a\cdot b}$ | *Dimensionless* | Rounder nucleus |
| 28 | M.28 | solidity | Nucleus roundness | Ratio of the true area of the nucleus to the convex area | Solidity $=A/A_{C}$ | *Dimensionless* | Rounder nucleus |
| 29 | G.1 | locs_density | Density of target across the entire nucleus | Total number of localizations divided by area of the nucleus | Localization Density$= N/A$ | [localizations/nm^2^] | More compact localizations global |
| 30 | G.2 | locs_number | Abundance of visualized target | Number of localizations in the nucleus | Number of Localizations $= N$ | [localizations] | More localizations |
| 31 | G.3 | log_voronoi_density_40 | Compaction of less dense compartment across entire nucleus  *For a nucleosome target, compartment associated to euchromatin* | 40^th^ percentile of the log-transformed Voronoi density distribution |  | Log_10_([localizations/nm^2^]) | More compact localizations global |
| 32 | G.4 | log_voronoi_density_70 | Compaction of denser compartment across entire nucleus  *For a nucleosome target, compartment associated to heterochromatin* | 70^th^ percentile of the log-transformed Voronoi density distribution |  | Log_10_([localizations/nm^2^]) | More compact localizations global |
| 33 | G.5 | log_voronoi_density_mean | Compaction across the entire nucleus | The average log-transformed Voronoi density of all localizations in the nucleus |  | Log_10_([localizations/nm^2^]) | More compact localizations global |
| 34 | G.6 | log_voronoi_density_skewness | Compaction across the entire nucleus | Skewness of the log-transformed Voronoi density distribution |  | *Dimensionless* | *N/A* |
| 35 | G.7 | log_voronoi_density_std | Heterogeneity of compaction across the entire nucleus | Standard deviation of the log-transformed Voronoi density distribution |  | Log_10_([localizations/nm^2^]) | *N/A* |
| 36 | G.8 | reduced_log_voronoi_density_40 | Normalized compaction of less dense compartment across entire nucleus  *For a nucleosome target, compartment associated to euchromatin* | 40^th^ percentile of the log-transformed reduced Voronoi density distribution |  | Log_10_([localizations/nm^2^]) | More compact localizations global |
| 37 | G.9 | reduced_log_voronoi_density_70 | Normalized compaction of less dense compartment across entire nucleus  *For a nucleosome target, compartment associated to euchromatin* | 70^th^ percentile of the log-transformed reduced Voronoi density distribution |  | Log_10_([localizations/nm^2^]) | More compact localizations global |
| 38 | G.10 | reduced_log_voronoi_density_mean | Compaction across entire nucleus | The average log-transformed reduced Voronoi density of all localizations in the nucleus | Compaction – The density of the Voronoi cell | Log_10_([localizations/nm^2^]) | More compact localizations global |
| 39 | I.1 | interior_loc_density | Frequency of visualized targets at the nucleus interior | Number of localizations at the nucleus interior divided by the area of the nucleus interior | Relaxed threshold (85% of the nucleus area) | [localizations/nm^2^] | More compact localizations interior |
| 40 | I.2 | interior_cluster_density | Frequency of packing domains at the nucleus interior per unit area | Number of DBSCAN domains at the nucleus interior divided by the area of the nucleus interior | Relaxed threshold (85% of the nucleus area) | [domains/nm^2^] | More domains interior |
| 41 | I.3 | interior_dbscan_cluster_n_clusters | Abundance of packing domains at the nucleus interior | Total number of DBSCAN domains in the nucleus interior | Strict threshold (95% of the nucleus area) | [localizations] | More domains interior |
| 42 | I.4 | interior_dbscan_cluster_n_locs | Abundance of visualized targets at the nucleus interior | Total number of localizations assigned to DBSCAN domains in the nucleus interior | Strict threshold (95% of the nucleus area) | [domains] | More localizations |
| 43 | I.5 | log_interior_dbscan_cluster_density_mean | Organization at nucleus interior: Domain compaction | Mean compaction of DBSCAN domains in the nucleus interior | Strict threshold (95% of the nucleus area)  Domain compaction – number of localizations in the domain divided by the domain area | Log_10_([localizations/nm^2^]) | More compact domains |
| 44 | I.6 | log_interior_dbscan_cluster_density_skewness | Organization at nucleus interior: Domain compaction | Skewness of the compaction of DBSCAN domains in the nucleus interior | Strict threshold (95% of the nucleus area)  Domain compaction – number of localizations in the domain divided by the domain area | *Dimensionless* | *N/A* |
| 45 | I.7 | log_interior_dbscan_cluster_density_std | Organization at nucleus interior: Heterogeneity of domain compaction | Standard deviation of the compaction of DBSCAN domains in the nucleus interior | Strict threshold (95% of the nucleus area)  Domain compaction – number of localizations in the domain divided by the domain area | Log_10_([localizations/nm^2^]) | More heterogenous domains interior |
| 46 | I.8 | log_interior_dbscan_cluster_gyration_radius_mean | Organization at nucleus interior: Packing domain dispersion | Mean radius of gyration of the DBSCAN domains in the nucleus interior | Strict threshold (95% of the nucleus area) | Log_10_([nm]) | *N/A* |
| 47 | I.9 | log_interior_dbscan_cluster_gyration_radius_skewness | Organization at nucleus interior: Packing domain dispersion | Skewness of the radius of gyration distribution of DBSCAN domains in the nucleus interior | Strict threshold (95% of the nucleus area) | *Dimensionless* | Rounder domains |
| 48 | I.10 | log_interior_dbscan_cluster_gyration_radius_std | Organization at nucleus interior: Heterogeneity of packing domain dispersion | Standard deviation of the radius of gyration distribution of DBSCAN domains in the nucleus interior | Strict threshold (95% of the nucleus area) | Log_10_([nm]) | More heterogenous domains interior |
| 49 | I.11 | log_interior_dbscan_cluster_n_locs_mean | Organization at nucleus interior: Abundance of visualized target within packing domains | Mean number of localizations in DBSCAN domains at the nucleus interior | Strict threshold (95% of the nucleus area) | [localizations] | More localizations |
| 50 | I.12 | log_interior_dbscan_cluster_n_locs_std | Organization at nucleus interior: Heterogeneity of the abundance of visualized target within packing domains | Standard deviation of the number of localizations in the DBSCAN domains at the nucleus interior | Strict threshold (95% of the nucleus area) | [localizations] | More heterogenous domains interior |
| 51 | I.13 | log_interior_dbscan_cluster_radius_mean | Organization at nucleus interior: Packing domain size | Mean radius of the DBSCAN domains in the nucleus interior | Strict threshold (95% of the nucleus area) | Log_10_([nm]) | Larger domains |
| 52 | I.14 | log_interior_dbscan_cluster_radius_skewness | Organization at nucleus interior: Packing domain size | Skewness of the radii of DBSCAN domains in the nucleus interior | Strict threshold (95% of the nucleus area) | *Dimensionless* | More heterogenous domains interior |
| 53 | I.15 | log_interior_dbscan_cluster_radius_std | Organization at nucleus interior: Heterogeneity of packing domain size | Standard deviation of the radii of DBSCAN domains in the nucleus interior | Strict threshold (95% of the nucleus area) | Log_10_([nm]) | *N/A* |
| 54 | I.16 | log_interior_dbscan_cluster_spacing_mean | Organization at nucleus interior: Spacing of neighboring packing domains | Mean spacing between neighboring DBSCAN domains in the nucleus interior | Strict threshold (95% of the nucleus area) | Log_10_([nm]) | More enrichment at periphery |
| 55 | I.17 | log_interior_dbscan_cluster_spacing_skewness | Organization at nucleus interior: Spacing of neighboring packing domains | Skewness of the spacing between neighboring DBSCAN domains in the nucleus interior | Strict threshold (95% of the nucleus area) | *Dimensionless* | *N/A* |
| 56 | I.18 | log_interior_dbscan_cluster_spacing_std | Organization at nucleus interior: Heterogeneity of the spacing of neighboring packing domains | Standard deviation of the spacing between neighboring DBSCAN domains in the nucleus interior | Strict threshold (95% of the nucleus area) | Log_10_([nm]) | More heterogenous domains interior |
| 57 | P.1 | periphery_loc_density | Frequency of visualized targets at the nucleus periphery | Number of DBSCAN domains at the nucleus periphery divided by the area of the nucleus periphery | Relaxed threshold (15% of the nucleus area) | [localizations/nm^2^] | More compact localizations periphery |
| 58 | P.2 | periphery_cluster_density | Frequency of packing domains at the nucleus periphery  *For a nucleosome target, these domains at the periphery may be associated to LADs* | Number of DBSCAN domains at the nucleus periphery divided by the area of the nucleus periphery | Relaxed threshold (15% of the nucleus area) | [domains/nm^2^] | More domains periphery |
| 59 | P.3 | periphery_dbscan_cluster_n_locs | Abundance of visualized targets within packing domains at the nucleus periphery  *For a nucleosome target, these domains at the periphery may be associated to LADs* | Total number of localizations assigned to DBSCAN domains at the nucleus periphery | Strict threshold (5% of the nucleus area) | [localizations] | More localizations |
| 60 | P.4 | periphery_dbscan_cluster_n_clusters | Abundance of packing domains at the nucleus periphery  *For a nucleosome target, these domains at the periphery may be associated to LADs* | Total number of DBSCAN domains at the nucleus periphery | Strict threshold (5% of the nucleus area) | [domains] | More domains periphery |
| 61 | P.5 | log_periphery_dbscan_cluster_density_mean | Organization at nucleus periphery: Domain compaction  *For a nucleosome target, these domains at the periphery may be associated to LADs* | Mean compaction of DBSCAN domains at the nucleus periphery | Strict threshold (5% of the nucleus area)  Domain compaction – number of localizations in the domain divided by the domain area | Log_10_([localizations/nm^2^]) | More compact domains |
| 62 | P.6 | log_periphery_dbscan_cluster_density_skewness | Organization at nucleus periphery: Domain compaction  *For a nucleosome target, these domains at the periphery may be associated to LADs* | Skewness of the compaction of DBSCAN domains at the nucleus periphery | Strict threshold (5% of the nucleus area)  Domain compaction – number of localizations in the domain divided by the domain area | *Dimensionless* | *N/A* |
| 63 | P.7 | log_periphery_dbscan_cluster_density_std | Organization at nucleus periphery: Heterogeneity of the domain compaction  *For a nucleosome target, these domains at the periphery may be associated to LADs* | Standard deviation of the compaction of DBSCAN domains at the nucleus periphery | Strict threshold (5% of the nucleus area)  Domain compaction – number of localizations in the domain divided by the domain area | Log_10_([localizations/nm^2^]) | More heterogenous domains periphery |
| 64 | P.8 | log_periphery_dbscan_cluster_gyration_radius_mean | Organization at nucleus periphery: Packing domain dispersion  *For a nucleosome target, these domains at the periphery may be associated to LADs* | Mean radius of gyration of the DBSCAN domains at the nucleus periphery | Strict threshold (5% of the nucleus area) | Log_10_([nm]) | Rounder domains |
| 65 | P.9 | log_periphery_dbscan_cluster_gyration_radius_skewness | Organization at nucleus periphery: Packing domain dispersion  *For a nucleosome target, these domains at the periphery may be associated to LADs* | Skewness of the radius of gyration distribution of DBSCAN domains at the nucleus periphery | Strict threshold (5% of the nucleus area) | *Dimensionless* | *N/A* |
| 66 | P.10 | log_periphery_dbscan_cluster_gyration_radius_std | Organization at nucleus periphery: Heterogeneity of the packing domain dispersion  *For a nucleosome target, these domains at the periphery may be associated to LADs* | Standard deviation of the radius of gyration distribution of DBSCAN domains at the nucleus periphery | Strict threshold (5% of the nucleus area) | Log_10_([nm]) | More heterogenous domains periphery |
| 67 | P.11 | log_periphery_dbscan_cluster_n_locs_mean | Organization at nucleus periphery: Abundance of visualized target within packing domains  *For a nucleosome target, these domains at the periphery may be associated to LADs* | Mean number of localizations in the DBSCAN domains at the nucleus periphery | Strict threshold (5% of the nucleus area) | Log_10_([localizations]) | More localizations |
| 68 | P.12 | log_periphery_dbscan_cluster_n_locs_std | Organization at nucleus periphery: Heterogeneity of the abundance of visualized target within packing domains  *For a nucleosome target, these domains at the periphery may be associated to LADs* | Standard deviation of the number of localizations in the DBSCAN domains at the nucleus periphery | Strict threshold (5% of the nucleus area) | Log_10_([localizations]) | More heterogenous domains periphery |
| 69 | P.13 | log_periphery_dbscan_cluster_radius_mean | Organization at nucleus periphery: Packing domain size  *For a nucleosome target, these domains at the periphery may be associated to LADs* | Mean radius of the DBSCAN domains at the nucleus periphery | Strict threshold (5% of the nucleus area) | Log_10_([nm]) | Larger domains |
| 70 | P.14 | log_periphery_dbscan_cluster_radius_skewness | Organization at nucleus periphery: Packing domain size  *For a nucleosome target, these domains at the periphery may be associated to LADs* | Skewness of the radii of DBSCAN domains at the nucleus periphery | Strict threshold (5% of the nucleus area) | *Dimensionless* | *N/A* |
| 71 | P.15 | log_periphery_dbscan_cluster_radius_std | Organization at nucleus periphery: Heterogeneity of the packing domain size  *For a nucleosome target, these domains at the periphery may be associated to LADs* | Standard deviation of the radii of DBSCAN domains at the nucleus periphery | Strict threshold (5% of the nucleus area) | Log_10_([nm]) | More heterogenous domains periphery |
| 72 | P.16 | periphery_to_interior_dbscan_cluster_density_ratio | Enrichment of packing domains at the nucleus periphery compared to the interior | Ratio of the number of DBSCAN domains in the nucleus periphery compared to the nucleus interior | Strict threshold (5% of the nucleus area) | *Dimensionless* | More enrichment at periphery |
| 73 | P.17 | periphery_to_interior_loc_density_ratio | Enrichment of target abundance at the nucleus periphery compared to the interior | Ratio of the number of localizations per unit area in the nucleus periphery compared to the nucleus interior | Strict threshold (5% of the nucleus area) | *Dimensionless* | More compact localizations periphery |
| 74 | L.1 | log_model_lad_segment_length_mean | Domain size at the nucleus periphery | Mean segment length distribution of modeled LAD clusters |  | [nm] | Rounder nucleus |
| 75 | L.2 | log_model_lad_segment_length_skewness | Domain size at the nucleus periphery | Standard deviation of the segment length distribution of modeled LAD clusters |  | *Dimensionless* | *N/A* |
| 76 | L.3 | log_model_lad_segment_length_std | Domain size heterogeneity at the nucleus periphery | Skewness of the segment length distribution of modeled LAD clusters |  | Log_10_([nm]) | More heterogenous domains periphery |
| 77 | L.4 | log_model_lad_segment_thickness_mean | Domain size at the nucleus periphery | Mean segment thickness distribution of modeled LAD clusters |  | Log_10_([nm]) | More enrichment at periphery |
| 78 | L.5 | log_model_lad_segment_thickness_skewness | Domain size asymmetry at the nucleus periphery | Standard deviation of the segment thickness distribution of modeled LAD clusters |  | *Dimensionless* | *N/A* |
| 79 | L.6 | log_model_lad_segment_thickness_std | Domain size heterogeneity at the nucleus periphery | Skewness of the segment thickness distribution of modeled LAD clusters |  | Log_10_([nm]) | More heterogenous domains periphery |
| 80 | L.7 | model_lad_thickness | Domain size at the nucleus periphery | Total modeled LAD area divided by the sum of length of the modeled LADs tangent to the nucleus boundary |  | Log_10_([nm]) | More enrichment at periphery |
| 81 | R.1 | radial_loc_density_ring_01 | Radially dependent organization | Localization density within an ellipse constructed with major/minor axes 10% those of the whole nucleus |  | [localizations/nm^2^] | More compact localizations interior |
| 82 | R.2 | radial_loc_density_ring_02 | Radially dependent organization | Localization density within a ring bounded by ellipses constructed with major/minor axes that span 10-20% those of the whole nucleus |  | [localizations/nm^2^] | More compact localizations interior |
| 83 | R.3 | radial_loc_density_ring_03 | Radially dependent organization | Localization density within a ring bounded by ellipses constructed with major/minor axes that span 20-30% those of the whole nucleus |  | [localizations/nm^2^] | More compact localizations interior |
| 84 | R.4 | radial_loc_density_ring_04 | Radially dependent organization | Localization density within a ring bounded by ellipses constructed with major/minor axes that span 30-40% those of the whole nucleus |  | [localizations/nm^2^] | More compact localizations interior |
| 85 | R.5 | radial_loc_density_ring_05 | Radially dependent organization | Localization density within a ring bounded by ellipses constructed with major/minor axes that span 40-50% those of the whole nucleus |  | [localizations/nm^2^] | More compact localizations interior |
| 86 | R.6 | radial_loc_density_ring_06 | Radially dependent organization | Localization density within a ring bounded by ellipses constructed with major/minor axes that span 50-60% those of the whole nucleus |  | [localizations/nm^2^] | More compact localizations interior |
| 87 | R.7 | radial_loc_density_ring_07 | Radially dependent organization | Localization density within a ring bounded by ellipses constructed with major/minor axes that span 60-70% those of the whole nucleus |  | [localizations/nm^2^] | More compact localizations interior |
| 88 | R.8 | radial_loc_density_ring_08 | Radially dependent organization | Localization density within a ring bounded by ellipses constructed with major/minor axes that span 70-80% those of the whole nucleus |  | [localizations/nm^2^] | More compact localizations interior |
| 89 | R.9 | radial_loc_density_ring_09 | Radially dependent organization | Localization density within a ring bounded by ellipses constructed with major/minor axes that span 80-90% those of the whole nucleus |  | [localizations/nm^2^] | More compact localizations periphery |
| 90 | R.10 | radial_loc_density_ring_10 | Radially dependent organization | Localization density outside an ellipse with major/minor axes 90% those of the whole nucleus |  | [localizations/nm^2^] | More compact localizations periphery |
| 91 | R.11 | radial_loc_density_ring_gradient_major_axis | Radial change in target abundance | Average change in localization density in the direction away from the centroid of the nucleus from R.1-R.8 normalized by the major axis of the nucleus |  | [localizations/nm^3^] | More heterogenous domains interior |
| 92 | R.12 | radial_loc_density_ring_gradient_minor_axis | Radial change in target abundance | Average change in localization density in the direction away from the centroid of the nucleus from R.1-R.8 normalized by the minor axis of the nucleus |  | [localizations/nm^3^] | More heterogenous domains interior |
| 93 | RD.1 | radial_dbscan_cluster_density_ring_01 | Radially dependent packing domain distribution | Number of DBSCAN domains per area within an ellipse with major/minor axes 10% that of the whole nucleus |  | [domains/nm^2^] | More domains interior |
| 94 | RD.2 | radial_dbscan_cluster_density_ring_02 | Radially dependent packing domain distribution | Number of DBSCAN domains per area within a ring bounded by ellipses constructed with major/minor axes that span 10-20% those of the whole nucleus |  | [domains/nm^2^] | More domains interior |
| 95 | RD.3 | radial_dbscan_cluster_density_ring_03 | Radially dependent packing domain distribution | Number of DBSCAN domains per area within a ring bounded by ellipses constructed with major/minor axes that span 20-30% those of the whole nucleus |  | [domains/nm^2^] | More domains interior |
| 96 | RD.4 | radial_dbscan_cluster_density_ring_04 | Radially dependent packing domain distribution | Number of DBSCAN domains per area within a ring bounded by ellipses constructed with major/minor axes that span 30-40% those of the whole nucleus |  | [domains/nm^2^] | More domains interior |
| 97 | RD.5 | radial_dbscan_cluster_density_ring_05 | Radially dependent packing domain distribution | Number of DBSCAN domains per area within a ring bounded by ellipses constructed with major/minor axes that span 40-50% those of the whole nucleus |  | [domains/nm^2^] | More domains interior |
| 98 | RD.6 | radial_dbscan_cluster_density_ring_06 | Radially dependent packing domain distribution | Number of DBSCAN domains per area within a ring bounded by ellipses constructed with major/minor axes that span 50-60% those of the whole nucleus |  | [domains/nm^2^] | More domains interior |
| 99 | RD.7 | radial_dbscan_cluster_density_ring_07 | Radially dependent packing domain distribution | Number of DBSCAN domains per area within a ring bounded by ellipses constructed with major/minor axes that span 60-70% those of the whole nucleus |  | [domains/nm^2^] | More domains interior |
| 100.0 | RD.8 | radial_dbscan_cluster_density_ring_08 | Radially dependent packing domain distribution | Number of DBSCAN domains per area within a ring bounded by ellipses constructed with major/minor axes that span 70-80% those of the whole nucleus |  | [domains/nm^2^] | More domains interior |
| 101 | RD.9 | radial_dbscan_cluster_density_ring_09 | Radially dependent packing domain distribution | Number of DBSCAN domains per area within a ring bounded by ellipses constructed with major/minor axes that span 80-90% those of the whole nucleus |  | [domains/nm^2^] | More domains periphery |
| 102 | RD.10 | radial_dbscan_cluster_density_ring_10 | Radially dependent packing domain distribution | Number of DBSCAN domains per area outside an ellipse with major/minor axes 90% those of the whole nucleus |  | [domains/nm^2^] | More domains periphery |
| 103 | RD.11 | radial_dbscan_cluster_density_ring_gradient_major_axis | Radial change in packing domain abundance | Average change in the number of DBSCAN domains per area in the direction away from the centroid of the nucleus from RD.1-RD.8 along the major axis of the nucleus |  | [domains/nm^3^] | More heterogenous domains interior |
| 104 | RD.12 | radial_dbscan_cluster_density_ring_gradient_minor_axis | Radial change in packing domain abundance | Average change in the number of DBSCAN domains per area in the direction away from the centroid of the nucleus from RD.1-RD.8 along the minor axis of the nucleus |  | [domains/nm^3^] | More heterogenous domains interior |
| 105 | V1.1 | voronoi_cluster_018nm^2_log_density_mean | Spatial organization at ultra-fine length scales: Domain compaction  *Threshold 1/4* | Mean number of localizations per Voronoi domain  *Clustered on localizations with a Voronoi area < 17.78 nm^2^* | Domain compaction – the number of localizations in the domain divided by the domain area | [localizations/nm^2^] | More compact domains |
| 106 | V1.2 | voronoi_cluster_018nm^2_log_density_skewness | Spatial organization at ultra-fine length scales: Domain compaction  *Threshold 1/4* | Skewness of the number of localizations per Voronoi domain  *Clustered on localizations with a Voronoi area < 17.78 nm^2^* | Domain compaction – the number of localizations in the domain divided by the domain area | *Dimensionless* |  |
| 107 | V1.3 | voronoi_cluster_018nm^2_log_density_std | Spatial organization at ultra-fine length scales: Domain compaction heterogeneity  *Threshold 1/4* | Standard deviation of number of localizations per Voronoi domain  *Clustered on localizations with a Voronoi area < 17.78 nm^2^* | Domain compaction – the number of localizations in the domain divided by the domain area | [localizations/nm^2^] | More heterogenous clusters global |
| 108 | V1.4 | voronoi_cluster_018nm^2_log_gyration_radius_mean | Spatial organization at ultra-fine length scales: Domain dispersion  *Threshold 1/4* | Mean radius of gyration of Voronoi domains  *Clustered on localizations with a Voronoi area < 17.78 nm^2^* |  | [nm] | Rounder domains |
| 109 | V1.5 | voronoi_cluster_018nm^2_log_gyration_radius_skewness | Spatial organization at ultra-fine length scales: Domain dispersion  *Threshold 1/4* | Skewness of Voronoi domains’ radius of gyration distribution  *Clustered on localizations with a Voronoi area < 17.78 nm^2^* |  | *Dimensionless* | *N/A* |
| 110 | V1.6 | voronoi_cluster_018nm^2_log_gyration_radius_std | Spatial organization at ultra-fine length scales: Domain dispersion heterogeneity  *Threshold 1/4* | Standard deviation of Voronoi domains’ radius of gyration distribution  *Clustered on localizations with a Voronoi area < 17.78 nm^2^* |  | [nm] | More heterogenous clusters global |
| 111 | V1.7 | voronoi_cluster_018nm^2_log_radius_mean | Spatial organization at ultra-fine length scales: Domain size  *Threshold 1/4* | Mean radius of Voronoi domains  *Clustered on localizations with a Voronoi area < 17.78 nm^2^* |  | [nm] | Larger domains |
| 112 | V1.8 | voronoi_cluster_018nm^2_log_radius_skewness | Spatial organization at ultra-fine length scales: Domain size  *Threshold 1/4* | Skewness of Voronoi domains’ radius distribution  *Clustered on localizations with a Voronoi area < 17.78 nm^2^* |  | *Dimensionless* | *N/A* |
| 113 | V1.9 | voronoi_cluster_018nm^2_log_radius_std | Spatial organization at ultra-fine length scales: Domain size heterogeneity  *Threshold 1/4* | Standard deviation of Voronoi domains’ radius distribution  *Clustered on localizations with a Voronoi area < 17.78 nm^2^* |  | [nm] | More heterogenous clusters global |
| 114 | V1.10 | voronoi_cluster_018nm^2_clusters_per_nucleus_area | Spatial organization at ultra-fine length scales: Domain frequency  *Threshold 1/4* | The number of Voronoi domains normalized by the nucleus area  *Clustered on localizations with a Voronoi area < 17.78 nm^2^* |  | [domains/nm^2^] | More domains global |
| 115 | V2.1 | voronoi_cluster_032nm^2_log_density_mean | Spatial organization at fine length scales: Domain compaction  *Threshold 2/4* | Mean number of localizations per Voronoi domain  *Clustered on localizations with a Voronoi area < 31.62 nm^2^* | Domain compaction – the number of localizations in the domain divided by the domain area | [localizations/nm^2^] | More compact domains |
| 116 | V2.2 | voronoi_cluster_032nm^2_log_density_skewness | Spatial organization at fine length scales: Domain compaction  *Threshold 2/4* | Skewness of the number of localizations per Voronoi domain  *Clustered on localizations with a Voronoi area < 31.62 nm^2^* | Domain compaction – the number of localizations in the domain divided by the domain area | *Dimensionless* | *N/A* |
| 117 | V2.3 | voronoi_cluster_032nm^2_log_density_std | Spatial organization at fine length scales: Domain compaction heterogeneity  *Threshold 2/4* | Standard deviation of number of localizations per Voronoi domain  *Clustered on localizations with a Voronoi area < 31.62 nm^2^* | Domain compaction – the number of localizations in the domain divided by the domain area | [localizations/nm^2^] | More heterogenous clusters global |
| 118 | V2.4 | voronoi_cluster_032nm^2_log_gyration_radius_mean | Spatial organization at fine length scales: Domain dispersion  *Threshold 2/4* | Mean radius of gyration of Voronoi domains  *Clustered on localizations with a Voronoi area < 31.62 nm^2^* |  | [nm] | Rounder domains |
| 119 | V2.5 | voronoi_cluster_032nm^2_log_gyration_radius_skewness | Spatial organization at fine length scales: Domain dispersion  *Threshold 2/4* | Skewness of Voronoi domains’ radius of gyration distribution  *Clustered on localizations with a Voronoi area < 31.62 nm^2^* |  | *Dimensionless* | *N/A* |
| 120 | V2.6 | voronoi_cluster_032nm^2_log_gyration_radius_std | Spatial organization at fine length scales: Domain dispersion heterogeneity  *Threshold 2/4* | Standard deviation of Voronoi domains’ radius of gyration distribution  *Clustered on localizations with a Voronoi area < 31.62 nm^2^* |  | [nm] | More heterogenous clusters global |
| 121 | V2.7 | voronoi_cluster_032nm^2_log_radius_mean | Spatial organization at fine length scales: Domain size  *Threshold 2/4* | Mean radius of Voronoi domains  *Clustered on localizations with a Voronoi area < 31.62 nm^2^* |  | [nm] | Larger domains |
| 122 | V2.8 | voronoi_cluster_032nm^2_log_radius_skewness | Spatial organization at fine length scales: Domain size  *Threshold 2/4* | Skewness of Voronoi domains’ radius distribution  *Clustered on localizations with a Voronoi area < 31.62 nm^2^* |  | *Dimensionless* | *N/A* |
| 123 | V2.9 | voronoi_cluster_032nm^2_log_radius_std | Spatial organization at fine length scales: Domain size heterogeneity  *Threshold 2/4* | Standard deviation of Voronoi domains’ radius distribution  *Clustered on localizations with a Voronoi area < 31.62 nm^2^* |  | [nm] | More heterogenous clusters global |
| 124 | V2.10 | voronoi_cluster_032nm^2_clusters_per_nucleus_area | Spatial organization at fine length scales: Domain frequency  *Threshold 2/4* | The number of Voronoi domains normalized by the nucleus area  *Clustered on localizations with a Voronoi area < 31.62 nm^2^* |  | [domains/nm^2^] | More domains global |
| 125 | V3.1 | voronoi_cluster_056nm^2_log_density_mean | Spatial organization at intermediate length scales: Domain compaction  *Threshold 3/4* | Mean number of localizations per Voronoi domain  *Clustered on localizations with a Voronoi area < 56.23 nm^2^* | Domain compaction – the number of localizations in the domain divided by the domain area | [localizations/nm^2^] | More compact domains |
| 126 | V3.2 | voronoi_cluster_056nm^2_log_density_skewness | Spatial organization at intermediate length scales: Domain compaction  *Threshold 3/4* | Skewness of the number of localizations per Voronoi domain  *Clustered on localizations with a Voronoi area < 56.23 nm^2^* | Domain compaction – the number of localizations in the domain divided by the domain area | *Dimensionless* | *N/A* |
| 127 | V3.3 | voronoi_cluster_056nm^2_log_density_std | Spatial organization at intermediate length scales: Domain compaction heterogeneity  *Threshold 3/4* | Standard deviation of number of localizations per Voronoi domain  *Clustered on localizations with a Voronoi area < 56.23 nm^2^* | Domain compaction – the number of localizations in the domain divided by the domain area | [localizations/nm^2^] | More heterogenous clusters global |
| 128 | V3.4 | voronoi_cluster_056nm^2_log_gyration_radius_mean | Spatial organization at intermediate length scales: Dispersion  *Threshold 3/4* | Mean radius of gyration of Voronoi domains  *Clustered on localizations with a Voronoi area < 56.23 nm^2^* |  | [nm] | Rounder domains |
| 129 | V3.5 | voronoi_cluster_056nm^2_log_gyration_radius_skewness | Spatial organization at intermediate length scales: Dispersion  *Threshold 3/4* | Skewness of Voronoi domains’ radius of gyration distribution  *Clustered on localizations with a Voronoi area < 56.23 nm^2^* |  | *Dimensionless* | *N/A* |
| 130 | V3.6 | voronoi_cluster_056nm^2_log_gyration_radius_std | Spatial organization at intermediate length scales: Dispersion heterogeneity  *Threshold 3/4* | Standard deviation of Voronoi domains’ radius of gyration distribution  *Clustered on localizations with a Voronoi area < 56.23 nm^2^* |  | [nm] | More heterogenous clusters global |
| 131 | V3.7 | voronoi_cluster_056nm^2_log_radius_mean | Spatial organization at intermediate length scales: Domain size  *Threshold 3/4* | Mean radius of Voronoi domains  *Clustered on localizations with a Voronoi area < 56.23 nm^2^* |  | [nm] | Larger domains |
| 132 | V3.8 | voronoi_cluster_056nm^2_log_radius_skewness | Spatial organization at intermediate length scales: Domain size  *Threshold 3/4* | Skewness of Voronoi domains’ radius distribution  *Clustered on localizations with a Voronoi area < 56.23 nm^2^* |  | *Dimensionless* | *N/A* |
| 133 | V3.9 | voronoi_cluster_056nm^2_log_radius_std | Spatial organization at intermediate length scales: Domain size heterogeneity  *Threshold 3/4* | Standard deviation of Voronoi domains’ radius distribution  *Clustered on localizations with a Voronoi area < 56.23 nm^2^* |  | [nm] | More heterogenous clusters global |
| 134 | V3.10 | voronoi_cluster_056nm^2_clusters_per_nucleus_area | Spatial organization at intermediate length scales: Domain frequency  *Threshold 3/4* | The number of Voronoi domains normalized by the nucleus area  *Clustered on localizations with a Voronoi area < 56.23 nm^2^* |  | [domains/nm^2^] | More domains global |
| 135 | V4.1 | voronoi_cluster_100nm^2_log_density_mean | Spatial organization at coarse length scales: Domain compaction  *Threshold 4/4* | Mean number of localizations per Voronoi domain  *Clustered on localizations with a Voronoi area < 100.0 nm^2^* | Domain compaction – the number of localizations in the domain divided by the domain area | [localizations/nm^2^] | More compact domains |
| 136 | V4.2 | voronoi_cluster_100nm^2_log_density_skewness | Spatial organization at coarse length scales: Domain compaction  *Threshold 4/4* | Skewness of the number of localizations per Voronoi domain  *Clustered on localizations with a Voronoi area < 100.0 nm^2^* | Domain compaction – the number of localizations in the domain divided by the domain area | *Dimensionless* | *N/A* |
| 137 | V4.3 | voronoi_cluster_100nm^2_log_density_std | Spatial organization at coarse length scales: Domain compaction heterogeneity  *Threshold 4/4* | Standard deviation of number of localizations per Voronoi domain  *Clustered on localizations with a Voronoi area < 100.0 nm^2^* | Domain compaction – the number of localizations in the domain divided by the domain area | [localizations/nm^2^] | More heterogenous clusters global |
| 138 | V4.4 | voronoi_cluster_100nm^2_log_gyration_radius_mean | Spatial organization at coarse length scales: Dispersion  *Threshold 4/4* | Average radius of gyration of Voronoi domains  *Clustered on localizations with a Voronoi area < 100.0 nm^2^* |  | [nm] | Rounder domains |
| 139 | V4.5 | voronoi_cluster_100nm^2_log_gyration_radius_skewness | Spatial organization at coarse length scales: Dispersion  *Threshold 4/4* | Skewness of Voronoi domains’ radius of gyration distribution  *Clustered on localizations with a Voronoi area < 100.0 nm^2^* |  | *Dimensionless* | *N/A* |
| 140 | V4.6 | voronoi_cluster_100nm^2_log_gyration_radius_std | Spatial organization at coarse length scales: Dispersion heterogeneity  *Threshold 4/4* | Standard deviation of Voronoi domains’ radius of gyration distribution  *Clustered on localizations with a Voronoi area < 100.0 nm^2^* |  | [nm] | More heterogenous clusters global |
| 141 | V4.7 | voronoi_cluster_100nm^2_log_radius_mean | Spatial organization at coarse length scales: Domain size  *Threshold 4/4* | Average radius of Voronoi domains  *Clustered on localizations with a Voronoi area < 100.0 nm^2^* |  | [nm] | Larger domains |
| 142 | V4.8 | voronoi_cluster_100nm^2_log_radius_skewness | Spatial organization at coarse length scales: Domain size  *Threshold 4/4* | Skewness of Voronoi domains’ radius distribution  *Clustered on localizations with a Voronoi area < 100.0 nm^2^* |  | *Dimensionless* | *N/A* |
| 143 | V4.9 | voronoi_cluster_100nm^2_log_radius_std | Spatial organization at coarse length scales: Domain size heterogeneity  *Threshold 4/4* | Standard deviation of Voronoi domains’ radius distribution  *Clustered on localizations with a Voronoi area < 100.0 nm^2^* |  | [nm] | More heterogenous clusters global |
| 144 | V4.10 | voronoi_cluster_100nm^2_clusters_per_nucleus_area | Spatial organization at coarse length scales: Domain frequency  *Threshold 4/4* | The number of Voronoi domains normalized by the nucleus area  *Clustered on localizations with a Voronoi area < 100.0 nm^2^* |  | [domains/nm^2^] | More domains global |

**Supplementary Table 4.** O-SNAP features set ranking methods for Feature Set Enrichment Analysis. By default, Signal-to-Noise is used. For a feature, $f$, from the entire set of features $\Omega$, the value $m(f)$ represents its corresponding ranking metric. The values $\mu_{A}(f)$ and $\mu_{B}(f)$ represent the average value, while $s_{A}(f)$ and $s_{B}(f)$ represent the standard deviations for $f$ for the samples of phenotypes $A$ and $B$, respectively. The sample counts for each phenotype are denoted by $n_{A}$ and $n_{B}$. The variable $x$ represents the feature value for an individual sample.

| Method | Option  *Argument value to pass* | Equation | Correction  *Applied when the increase in feature* $\boldsymbol{f}$ *is opposite to the pattern described by the feature set* |
| --- | --- | --- | --- |
| Signal-to-Noise | S2N | $m(f)=\frac{\mu_{B}(f)-\mu_{A}(f)}{s_{B}(f)+s_{A}(f)}$ | $-m(f)$ |
| *t*-test | ttest | $m\left( f \right)=\frac{\mu_{B}(f)-\mu_{A}(f)}{\left( s_{B}(f)+s_{A}(f) \right)\cdot\sqrt{\frac{s_{A}^{2}(f)}{n_{A}}+\frac{s_{B}^{2}(f)}{n_{B}}}}$ | $-{m(f)}_{f}$ |
| Ratio | ratio | $m\left( f \right)=\frac{\sum_{i=1}^{n_{A}} 2^{x_{A,i}}}{n_{A}}\div\frac{\sum_{i=1}^{n_{B}} 2^{x_{B,i}}}{n_{B}}$ | $m^{-1}(f)$ |
| Difference | diff | $m_{f}=\mu_{B}-\mu_{A}$ | $-m(f)$ |
| Log2 Ratio | log2_ratio | $m\left( f \right)=\log_{2} \frac{\left( \frac{\sum_{i=1}^{n_{A}} 2^{x_{A,i}}}{n_{A}} \right)}{\left( \frac{\sum_{i=1}^{n_{B}} 2^{x_{B,i}}}{n_{B}} \right)}$ | $-m(f)$ |
| Weighted average difference | WAD | $AD\left( f \right)= \mu_{B}-\mu_{A}$  $w=\frac{\frac{\mu_{A}(f)+\mu_{B}(f)}{2}-\min\left( \left\{ \mu_{A}(f) \vert f\in\Omega\right\} \right)}{\max\left( \left\{ \mu_{A}(f) \vert f\in\Omega\right\} \right)-\min\left( \left\{ \mu_{A}(f) \vert f\in\Omega\right\} \right)}$  $m\left( f \right)=w(f)*AD\left( f \right)$ | Use the additive inverse of the average difference:  $AD=-\left( \mu_{B}-\mu_{A} \right)$ |
| Fold change rank ordering statistics | FCROS | The truncated mean on the matrix of fold changes from pairwise comparisons between the two phenotypes^1^ | Prior to the calculation of the truncated mean, use the multiplicative inverse of the rank values |
| Moderated Welch test | MWT | The Moderated Welch Statistic^2^ | $-m(f)$ |

**Supplementary Table 5.** Criteria to generate candidate trajectory inference methods to train on O-SNAP feature data of the heterokaryon system.

| Question ID | Answer |
| --- | --- |
| multiple_disconnected | FALSE |
| expect_topology | TRUE |
| expected_topology | "linear" |
| n_cells | 350 |
| n_features | 144 |
| memory | "2GB" |
| prior_information | "start_id","start_n","end_n","groups_id","groups_network" |
| docker | TRUE |

**Supplementary Table 6.** O-SNAP features satisfying |fold change| > 2 and p-value < 0.5, following adjustment by the Benjamini-Hochberg method for multiple hypothesis testing between H2B-labeled control and TSA-treated cells.

| Short-hand | Feature | Log_2_(Fold Change) | Adj p-Value |
| --- | --- | --- | --- |
| I.3 | interior_dbscan_cluster_n_clusters | 1.00 | 1.73E-15 |
| P.6 | log_periphery_dbscan_cluster_density_skewness | 1.03 | 5.12E-08 |
| P.9 | log_periphery_dbscan_cluster_gyration_radius_skewness | 2.06 | 2.65E-04 |

**Supplementary Table 7.** O-SNAP features satisfying |fold change| > 2 and p-value < 0.5, following adjustment by the Benjamini-Hochberg method for multiple hypothesis testing between H2B-labeled cells derived from healthy and tendinosis donors.

| Short-hand | Feature | Log_2_(Fold Change) | Adj p-Value |
| --- | --- | --- | --- |
| I.4 | interior_dbscan_cluster_n_locs | -2.83 | 3.90E-03 |
| I.3 | interior_dbscan_cluster_n_clusters | -2.38 | 3.38E-05 |
| I.2 | interior_cluster_density | -2.02 | 3.68E-07 |
| RD.7 | radial_dbscan_cluster_density_ring_07 | -1.77 | 9.38E-04 |
| RD.5 | radial_dbscan_cluster_density_ring_05 | -1.72 | 9.56E-04 |
| V3.1 | voronoi_cluster_056nm^2_log_density_mean | -1.72 | 4.10E-04 |
| RD.6 | radial_dbscan_cluster_density_ring_06 | -1.68 | 1.26E-03 |
| RD.4 | radial_dbscan_cluster_density_ring_04 | -1.56 | 1.99E-03 |
| RD.3 | radial_dbscan_cluster_density_ring_03 | -1.54 | 1.58E-03 |
| P.4 | periphery_dbscan_cluster_n_clusters | -1.49 | 6.65E-05 |
| RD.1 | radial_dbscan_cluster_density_ring_01 | -1.48 | 3.11E-03 |
| P.2 | periphery_cluster_density | -1.43 | 2.17E-09 |
| G.2 | locs_number | -1.36 | 5.38E-03 |
| RD.2 | radial_dbscan_cluster_density_ring_02 | -1.33 | 4.22E-03 |
| R.7 | radial_loc_density_ring_07 | -1.18 | 9.19E-04 |
| I.1 | interior_loc_density | -1.17 | 9.04E-05 |
| R.6 | radial_loc_density_ring_06 | -1.11 | 9.19E-04 |
| R.5 | radial_loc_density_ring_05 | -1.10 | 7.81E-04 |
| R.2 | radial_loc_density_ring_02 | -1.10 | 2.31E-03 |
| R.4 | radial_loc_density_ring_04 | -1.04 | 1.45E-03 |
| R.3 | radial_loc_density_ring_03 | -1.04 | 9.56E-04 |
| R.8 | radial_loc_density_ring_08 | -1.03 | 1.68E-03 |
| P.17 | periphery_to_interior_loc_density_ratio | 1.24 | 3.38E-05 |

**Supplementary Table 8.** O-SNAP features satisfying |fold change| > 2 and p-value < 0.5, following adjustment by the Benjamini-Hochberg method for multiple hypothesis testing between control hFb and hFb nuclei 48-hr post cell fusion within the heterokaryon labeled for H3K27me3.

| Short-hand | Feature | Log_2_(Fold Change) | Adj p-Value |
| --- | --- | --- | --- |
| RD.1 | radial_dbscan_cluster_density_ring_01 | -2.39 | 1.80E-20 |
| RD.2 | radial_dbscan_cluster_density_ring_02 | -2.29 | 9.44E-24 |
| V1.10 | voronoi_cluster_018nm^2_clusters_per_nucleus_area | -2.23 | 6.11E-16 |
| RD.3 | radial_dbscan_cluster_density_ring_03 | -2.18 | 3.86E-25 |
| R.1 | radial_loc_density_ring_01 | -2.17 | 2.23E-27 |
| R.2 | radial_loc_density_ring_02 | -2.13 | 6.21E-31 |
| I.4 | interior_dbscan_cluster_n_locs | -2.07 | 1.94E-17 |
| I.6 | log_interior_dbscan_cluster_density_skewness | -2.04 | 2.65E-20 |
| RD.4 | radial_dbscan_cluster_density_ring_04 | -2.03 | 8.03E-28 |
| R.3 | radial_loc_density_ring_03 | -2.03 | 2.57E-29 |
| V2.10 | voronoi_cluster_032nm^2_clusters_per_nucleus_area | -2.02 | 4.85E-25 |
| RD.5 | radial_dbscan_cluster_density_ring_05 | -1.96 | 8.23E-31 |
| R.4 | radial_loc_density_ring_04 | -1.93 | 2.57E-32 |
| R.5 | radial_loc_density_ring_05 | -1.85 | 1.36E-32 |
| RD.6 | radial_dbscan_cluster_density_ring_06 | -1.82 | 2.02E-33 |
| R.6 | radial_loc_density_ring_06 | -1.81 | 9.72E-33 |
| I.1 | interior_loc_density | -1.72 | 3.23E-34 |
| I.2 | interior_cluster_density | -1.72 | 3.71E-36 |
| R.7 | radial_loc_density_ring_07 | -1.69 | 1.31E-33 |
| RD.7 | radial_dbscan_cluster_density_ring_07 | -1.64 | 9.45E-35 |
| V3.10 | voronoi_cluster_056nm^2_clusters_per_nucleus_area | -1.57 | 1.78E-39 |
| I.17 | log_interior_dbscan_cluster_spacing_skewness | -1.56 | 0.003387908 |
| R.8 | radial_loc_density_ring_08 | -1.53 | 5.63E-32 |
| RD.8 | radial_dbscan_cluster_density_ring_08 | -1.49 | 1.31E-33 |
| G.1 | locs_density | -1.46 | 2.47E-32 |
| P.3 | periphery_dbscan_cluster_n_locs | -1.42 | 6.97E-06 |
| R.9 | radial_loc_density_ring_09 | -1.34 | 8.35E-29 |
| RD.9 | radial_dbscan_cluster_density_ring_09 | -1.30 | 8.73E-28 |
| RD.10 | radial_dbscan_cluster_density_ring_10 | -1.19 | 3.45E-22 |
| L.7 | model_lad_thickness | -1.18 | 1.82E-08 |
| R.10 | radial_loc_density_ring_10 | -1.17 | 3.95E-21 |
| V2.5 | voronoi_cluster_032nm^2_log_gyration_radius_skewness | -1.17 | 1.30E-16 |
| V3.5 | voronoi_cluster_056nm^2_log_gyration_radius_skewness | -1.09 | 5.17E-16 |
| R.11 | radial_loc_density_ring_gradient_major_axis | -1.08 | 6.32E-08 |
| R.12 | radial_loc_density_ring_gradient_minor_axis | -1.03 | 4.04E-07 |
| P.1 | periphery_loc_density | -1.03 | 2.27E-16 |
| G.8 | reduced_log_voronoi_density_40 | 1.09 | 2.51E-26 |

**Supplementary Table 9.** O-SNAP features satisfying |fold change| > 2 and p-value < 0.5, following adjustment by the Benjamini-Hochberg method for multiple hypothesis testing between control hFb and hFb nuclei 48-hr post cell fusion within the heterokaryon labeled for H3K9ac.

| Short-hand | Feature | Log_2_(Fold Change) | Adj p-Value |
| --- | --- | --- | --- |
| L.5 | log_model_lad_segment_thickness_skewness | -3.11 | 1.69E-07 |
| G.6 | log_voronoi_density_skewness | -1.18 | 4.38E-03 |
| P.4 | periphery_dbscan_cluster_n_clusters | 1.06 | 1.11E-07 |
| L.7 | model_lad_thickness | 1.43 | 1.26E-06 |
| P.3 | periphery_dbscan_cluster_n_locs | 1.50 | 1.30E-04 |

**Supplementary Table 10.** O-SNAP features satisfying |fold change| > 2 and p-value < 0.5, following adjustment by the Benjamini-Hochberg method for multiple hypothesis testing between H2B-labeled control and D-Alanine treated MCF10A cells overexpressing the histone variant H3.1.

| Short-hand | Feature | Log_2_(Fold Change) | Adj p-Value |
| --- | --- | --- | --- |
| RD.12 | radial_dbscan_cluster_density_ring_gradient_minor_axis | -2.15 | 1.97E-02 |
| RD.11 | radial_dbscan_cluster_density_ring_gradient_major_axis | -2.13 | 3.07E-02 |
| L.5 | log_model_lad_segment_thickness_skewness | -1.32 | 1.21E-04 |
| I.14 | log_interior_dbscan_cluster_radius_skewness | 1.03 | 1.20E-10 |
| P.3 | periphery_dbscan_cluster_n_locs | 1.13 | 2.83E-06 |
| L.7 | model_lad_thickness | 1.13 | 2.12E-07 |
| I.6 | log_interior_dbscan_cluster_density_skewness | 1.50 | 4.24E-06 |
| I.17 | log_interior_dbscan_cluster_spacing_skewness | 2.43 | 2.79E-05 |

**Supplementary Table 11.** O-SNAP features satisfying |fold change| > 2 and p-value < 0.5, following adjustment by the Benjamini-Hochberg method for multiple hypothesis testing between H2B-labeled control and D-Alanine treated MCF10A overexpressing the histone variant H3.2.

| Short-hand | Feature | Log_2_(Fold Change) | Adj p-Value |
| --- | --- | --- | --- |
| I.17 | log_interior_dbscan_cluster_spacing_skewness | -1.89 | 3.97E-02 |

**Supplementary Table 12.** O-SNAP features satisfying |fold change| > 2 and p-value < 0.5, following adjustment by the Benjamini-Hochberg method for multiple hypothesis testing between P0 and P6 human chondrocytes labeled for H2B derived from Donor A.

| Short-hand | Feature | Log_2_(Fold Change) | Adj p-Value |
| --- | --- | --- | --- |
| RD.12 | radial_dbscan_cluster_density_ring_gradient_minor_axis | -6.40 | 2.62E-02 |
| L.5 | log_model_lad_segment_thickness_skewness | -2.43 | 9.50E-03 |
| V1.10 | voronoi_cluster_018nm^2_clusters_per_nucleus_area | 1.13 | 4.27E-04 |
| I.4 | interior_dbscan_cluster_n_locs | 1.14 | 5.51E-05 |
| L.7 | model_lad_thickness | 1.30 | 5.90E-03 |
| P.3 | periphery_dbscan_cluster_n_locs | 1.44 | 6.99E-03 |

**Supplementary Table 13.** O-SNAP features satisfying |fold change| > 2 and p-value < 0.5, following adjustment by the Benjamini-Hochberg method for multiple hypothesis testing between P0 and P6 human chondrocytes labeled for H2B derived from Donor B.

| Short-hand | Feature | Log_2_(Fold Change) | Adj p-Value |
| --- | --- | --- | --- |
| I.17 | log_interior_dbscan_cluster_spacing_skewness | -2.75 | 1.30E-02 |
| L.5 | log_model_lad_segment_thickness_skewness | -2.00 | 2.23E-03 |
| V3.1 | voronoi_cluster_056nm^2_log_density_mean | -1.90 | 3.60E-05 |
| V4.1 | voronoi_cluster_100nm^2_log_density_mean | -1.41 | 1.17E-06 |
| V1.10 | voronoi_cluster_018nm^2_clusters_per_nucleus_area | 1.11 | 1.56E-05 |
| L.7 | model_lad_thickness | 1.81 | 3.60E-05 |
| P.3 | periphery_dbscan_cluster_n_locs | 2.10 | 4.99E-05 |

**Supplementary Table 14.** O-SNAP features satisfying |fold change| > 2 and p-value < 0.5, following adjustment by the Benjamini-Hochberg method for multiple hypothesis testing between P0 and P3 human chondrocytes labeled for H2B derived from Donor A.

| Short-hand | Feature | Log_2_(Fold Change) | Adj p-Value |
| --- | --- | --- | --- |
| P.6 | log_periphery_dbscan_cluster_density_skewness | -1.24 | 1.93E-02 |
| P.4 | periphery_dbscan_cluster_n_clusters | 1.51 | 4.53E-03 |
| V3.1 | voronoi_cluster_056nm^2_log_density_mean | 1.56 | 3.21E-03 |
| V4.1 | voronoi_cluster_100nm^2_log_density_mean | 3.18 | 9.09E-03 |

**Supplementary Table 15.** O-SNAP features satisfying |fold change| > 2 and p-value < 0.5, following adjustment by the Benjamini-Hochberg method for multiple hypothesis testing between P0 and P3 human chondrocytes labeled for H2B derived from Donor B.

| Short-hand | Feature | Log_2_(Fold Change) | Adj p-Value |
| --- | --- | --- | --- |
| R.12 | radial_loc_density_ring_gradient_minor_axis | -2.76 | 3.65E-02 |
| L.6 | log_model_lad_segment_thickness_std | -1.33 | 1.48E-05 |
| G.2 | locs_number | 1.21 | 2.42E-05 |
| V1.10 | voronoi_cluster_018nm^2_clusters_per_nucleus_area | 1.23 | 1.04E-03 |
| I.4 | interior_dbscan_cluster_n_locs | 1.98 | 2.88E-04 |
| L.7 | model_lad_thickness | 2.22 | 1.66E-04 |
| P.3 | periphery_dbscan_cluster_n_locs | 3.12 | 1.06E-04 |

**Supplementary Table 16.** O-SNAP features satisfying |fold change| > 2 and p-value < 0.5, following adjustment by the Benjamini-Hochberg method for multiple hypothesis testing between P3 and P6 human chondrocytes labeled for H2B derived from Donor A.

| Short-hand | Feature | Log_2_(Fold Change) | Adj p-Value |
| --- | --- | --- | --- |
| V4.1 | voronoi_cluster_100nm^2_log_density_mean | -3.59 | 1.87E-03 |
| V3.1 | voronoi_cluster_056nm^2_log_density_mean | -1.21 | 8.10E-03 |
| M.3 | bending_energy_norm_area | 1.48 | 2.60E-02 |

**Supplementary Table 17.** O-SNAP features satisfying |fold change| > 2 and p-value < 0.5, following adjustment by the Benjamini-Hochberg method for multiple hypothesis testing between P3 and P6 human chondrocytes labeled for H2B derived from Donor B.

| Short-hand | Feature | Log_2_(Fold Change) | Adj p-Value |
| --- | --- | --- | --- |
| P.3 | periphery_dbscan_cluster_n_locs | -1.02 | 1.59E-02 |
| I.4 | interior_dbscan_cluster_n_locs | -1.00 | 1.44E-02 |

**Supplementary Table 18.** Filters applied to localization data processed in NimOS.

| Method | Min | Max |
| --- | --- | --- |
| Photon count | 300 | 100000000 |
| Localization precision X (nm) | 0.00 | 30.00 |
| Localization precision Y (nm) | 0.00 | 30.00 |
| Sigma X (nm) | 22.25 | 445.07 |
| Sigma Y (nm) | 22.25 | 445.07 |
| p-value | 0.0000 | 1.0000 |
